# Genome-scale perturb-seq in primary human CD4+ T cells maps context-specific regulators of T cell programs and human immune traits

**DOI:** 10.64898/2025.12.23.696273

**Authors:** Ronghui Zhu, Emma Dann, Jun Yan, Justine Reyes Retana, Ryunosuke Goto, Reese C. Guitche, Lillian K. Petersen, Mineto Ota, Jonathan K. Pritchard, Alexander Marson

**Author notes:** These authors contributed equally (Ronghui Zhu, Emma Dann).

## Abstract

Gene regulatory networks encode the fundamental logic of cellular functions, but systematic network mapping remains challenging, especially in cell states relevant to human biology and disease. Here, we perturbed all expressed genes across 22 million primary human CD4+ T cells from four donors and developed a probe-based perturb-seq platform to measure the transcriptome effects in cells at rest and after stimulation. These data allow us to map genes that regulate known and novel pathways, including novel regulators of cytokine production. Importantly, active regulators and the gene programs they control change dramatically across stimulation conditions. Perturbation signatures enabled us to model T cell states observed in population-scale transcriptomic atlases, nominating regulators of Th1 and Th2 polarization and of age-related T cell phenotypes. Finally, we leveraged perturb-seq to implicate context-specific gene regulatory pathways in autoimmune disease risk. Our data provide a foundational resource to decode human immune function and genetic variation and for new approaches to study gene regulatory networks.

## Introduction

Gene regulatory networks (GRNs) constitute the fundamental logic by which cells organize biological functions. Mapping these networks uncovers circuit design principles [1], guides precise cellular engineering [2], and mechanistically links genetic variations to complex human traits [3]. While advances in single-cell transcriptomics have allowed for the detailed observation of cellular states, inferring causal regulatory relationships from observational data remains challenging [4].

Large scale CRISPR screens have ushered in a new era of functional genetics, where the consequences of perturbing each gene can be tested systematically. We and others have coupled CRISPR screens with fluorescence-activated cell sorting (FACS)-based selection strategies to enable comprehensive mapping of upstream regulators of key genes [5,6]. This approach provides a foothold to discover gene regulators but is laborious, limited to studying one target at a time, and does not immediately reveal the downstream transcriptional programs controlled by each regulator. More recently, perturb-seq (perturbation combined with single-cell RNA sequencing) has emerged as a powerful method to overcome these limitations. By coupling genetic perturbations with high-dimensional transcriptomic readouts, perturb-seq allows for systematic measurement of regulatory relationships and the unbiased discovery of gene functions [7–9].

Previous genome-scale studies have mapped the regulatory logic of general cellular processes in immortalized model cell lines [9,10]. However, much of human biology is specific to specialized cell types and their dynamic responses to defined stimuli, and influenced by inter-individual variability. Perturbations of immortalized cell lines often fail to recapitulate these features. Perturb-seq in primary human cells has generally been restricted to targeted libraries or arrayed formats. This targeted approach inherently biases discovery toward known biology, preventing the identification of novel regulators or unexpected connections between distinct biological processes that can only be captured by unbiased, genome-scale interrogation.

Here we developed a scalable, probe-based perturb-seq platform to extend perturbation measurements to genome-scale in primary human cells with high resolution measurements of the transcriptional consequences of each perturbation. We focused on CD4+ T cells due to their central role in orchestrating adaptive immunity and their relevance in many autoimmune diseases [11]. In response to stimulation via their T cell receptor (TCR), co-stimulation (via CD28) and additional extracellular signals, CD4+ T cells polarize into distinct effector states defined by transcription factor, surface receptors and cytokine expression signatures. Well-characterized polarized cell states, including Th1 and Th2 CD4+ T cells, drive distinct modes of immune responses and contribute to different autoimmune inflammatory conditions [12]. Despite decades of work on CD4+ T cell differentiation, we still lack a comprehensive map of the dynamic GRNs that shape the state of CD4+ T cells as they respond to stimulation.

We employed genome-scale CRISPR interference (CRISPRi) to systematically knock down all expressed genes in 22 million primary human CD4+ T cells. By profiling these cells at rest and after stimulation, we provide a dynamic atlas of each perturbation’s effects on human T cell state. We demonstrate the utility of this resource through three complementary applications: 1) systematic discovery of context-dependent regulators: by leveraging the scale and multi-condition nature of our data, we identify novel regulators of key T-cell specific and stimulation condition-specific gene programs that were previously undetectable in targeted or static screens; 2) modeling of natural cell states via perturbation signatures: we demonstrate that perturbation signatures derived *ex vivo* can be used to predict regulators of natural cell states, captured in observational data from population-scale cell atlases; and 3) mechanistic interpretation of gene-trait associations: by intersecting our perturb-seq measurements with population studies of human disease genetics, we link known and novel pathways to complex traits, moving beyond single gene-locus associations to define the functional pathways driving disease susceptibility. Taken together, this study offers new approaches to integrate genome-scale perturb-seq studies of human primary cells with human cell atlases and human genetics and also provides a foundational resource for human immunology.

## Results

### A scalable platform for perturb-seq in primary CD4+ T cells

To systematically map functional gene regulatory networks in human primary T cells, we compiled a genome-scale library targeting a union of all expressed genes in human CD4+ T cells and all transcription factors (12,748 genes in total, see Methods). Primary naive CD4+ T cells isolated from four human donors were stimulated, transduced with CRISPRi machinery and the genome-scale library via an optimized lentiviral protocol [6], and expanded *ex vivo* (Figure 1A, Suppl. Table 1, Methods). Cells were then allowed to rest without CD3/CD28 stimulation. To capture the dynamic regulatory circuits during T cell activation [13], we collected perturbed cells in three different experimental conditions – rested (Rest), 8 hours after re-stimulation (Stim8hr), and 48 hours after re-stimulation (Stim48hr).

**Figure 1.**
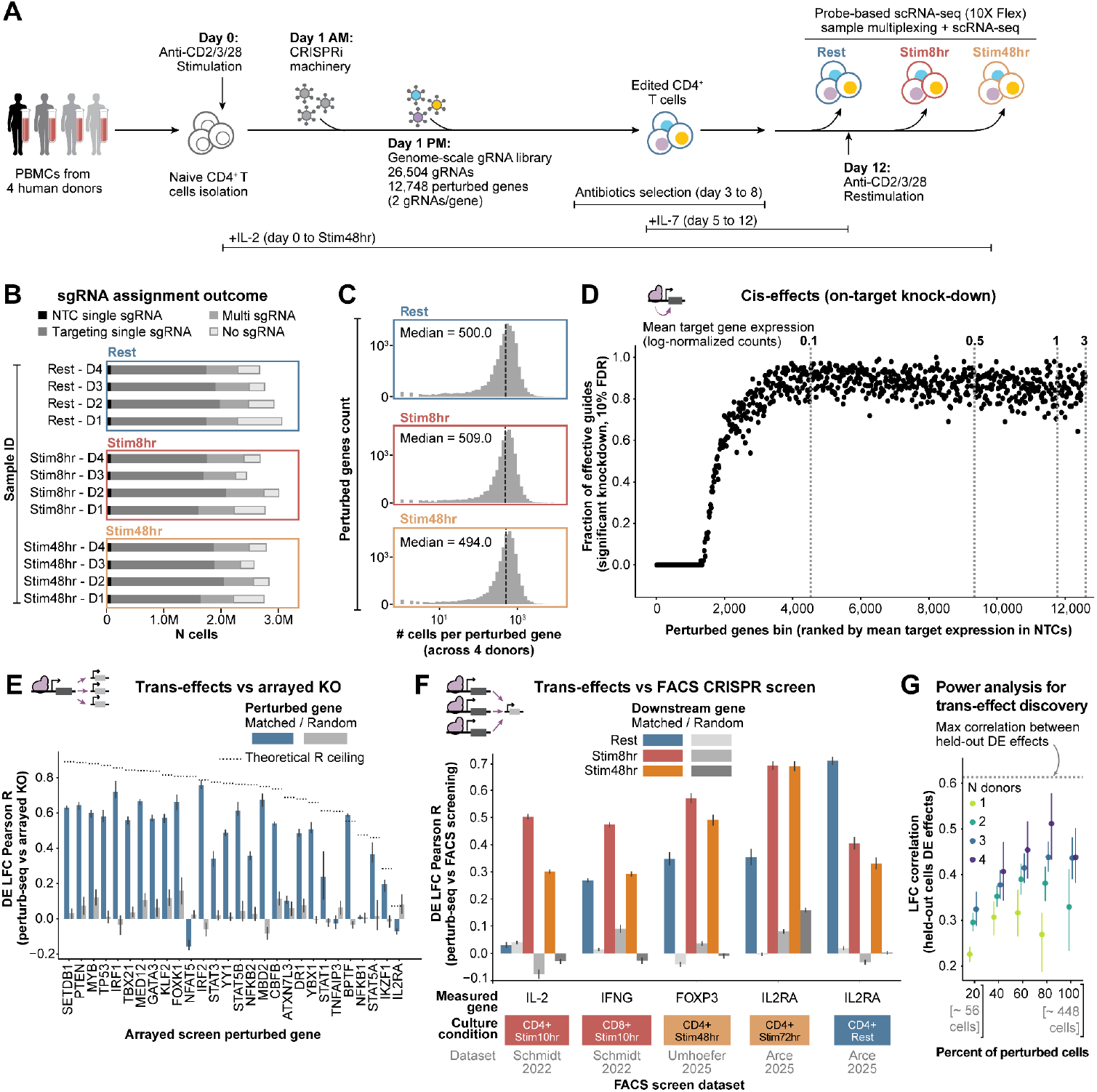
Genome-scale perturb-seq experimental design and quality control. (**A**) Experimental design of genome-scale perturb-seq in CD4+ T cells from multiple human donors across conditions (see Methods). (**B**) Number of cells (y-axis) for each biological sample (x-axis, condition-donor) with transcriptome passing quality control. Bars are colored by the outcome of guide RNA assignment (NTC: non-targeting controls). (**C**) Distribution of the number of cells harboring each gene perturbation (x-axis, log10 scale) across culture conditions (rows). The dotted line denotes the median. (**D**) Fraction of guides inducing significant on-target knock-down (y-axis) versus baseline expression of perturbed genes (x-axis). Each point represents 100 genes grouped by similar expression levels. Dotted vertical lines denote mean expression (log-normalized counts) per bin. (**E**) Comparison of trans effects between perturb-seq and arrayed knockout (KO) RNA-seq [5,16]: for each perturbed gene (x-axis), Pearson correlation of gene expression Log fold changes (LFCs) across all measured genes (y-axis) is shown. Blue bars: comparison of same gene perturbed in both platforms; grey bars: comparison on different perturbed genes. Error bars: 95% confidence interval (CI) from bootstrapping. Dotted line: theoretical maximum correlation given LFC noise. (**F**) Comparison of trans effects between perturb-seq and published CRISPRi FACS-based screens [6,13,17]. For each measured gene (x-axis), Pearson correlation of expression LFCs across all perturbed genes (y-axis) is shown. Colored bars: same gene measured in both platforms; grey-scale bars: expression-matched different genes. Error bars: 95% CI from bootstrapping. Stimulation conditions of FACS screens are indicated below x-axis. (**G**) Pearson correlation between LFCs from downsampled and held-out validation data (5 lanes, all donors) for genes significant at 10% FDR. X-axis: fraction of perturbed cells (20–100%, ~55–450 cells). Colors: number of donors (1–4). Error bars: variability across independent downsampling splits and perturbed genes. Aggregated results across 12 perturbed genes are shown (see Suppl. Figure 9 for per-gene results). Grey dotted line: maximum correlation between perturbation effects in independent held-out splits.

To profile tens of millions of cells for this genome-scale, multi-donor, and multi-timepoint study, we leveraged recent advances in probe-based single cell RNA sequencing (scRNA-seq) technology (10x Flex). This technology provides sample collection flexibility by allowing for cell fixation, offers superior sensitivity for low-abundance transcripts [14] and enables “superloading” to profile tens of millions of cells across < 80 lanes – significantly fewer than traditional methods [10].

In total, we collected 12 pools of CRISPRi-perturbed cells from four healthy human donors across 3 stimulation conditions (Suppl. Figure 1A). We recovered a consistent number of mRNA UMIs, genes, and guide UMIs per cell across biological replicates and lanes (Suppl. Figure 1B, mean UMI/cell per condition: Rest = 10,080; Stim8hr = 14,977; Stim48hr = 13,373). After quality control filtering and gRNA assignment, we recovered 33.4M cells with high quality transcriptomes (> 500 genes/cell and < 5% mitochondrial reads), with minimal technical batch effects and expected expression patterns for canonical T cell activation markers across stimulation conditions (Suppl. Figure 2). Of these, 21.99M cells (65.8%) were assigned a single targeting or non-targeting guide, 4.69M (14.0%) had no guide confidently captured, and 6.70M (20.0%) had multiple guides assigned (Figure 1B). Cells with multiple guides assigned had systematically higher numbers of UMIs per cell, indicating these are likely doublets (Suppl. Figure 1C). Conversely, cells with no assigned guide showed systematically lower expression of the puromycin resistance transgene associated with gRNA expressing cassette, suggesting escape from antibiotics selection, rather than failure to detect the gRNA probe (Suppl. Figure 1C). Of the 26,504 guides in the library, 23,573 (88.9%) were detected in at least 50 cells across samples. We recovered on average 575 cells per perturbed gene in each condition (Figure 1C). Outlier perturbations with significantly lower numbers of perturbed cells mostly consisted of essential genes, where we expect knock-down to have an effect on cell survival (Suppl. Figure 1D).

Compared with recently released genome-scale perturb-seq datasets, the median transcripts per cell per context in our data (~10,000) was comparable to other genetic perturbation datasets in cell lines [9,10], but we achieved approximately double the cell coverage per perturbation (Suppl. Figure 1E). In contrast, the Tahoe-100M chemical perturbation dataset [15] captures more cells per perturbation but with nearly an order of magnitude lower read depth (median transcripts per cell per context = 1,780).

### Trans effects of genetic perturbations

We next assessed the effects of genetic perturbations on transcriptome-wide gene expression, starting from on-target knockdown efficiency. Of the 24,174 guides detected across conditions, target gene expression was significantly reduced for 73% of tested guides compared to non-targeting control (NTC) guides (Figure 1D, Suppl. Figure 3A). The subset of guides that failed to achieve significant knockdown predominantly targeted genes with very low baseline expression (Suppl. Figure 3B). Quality control metrics for individual guides are provided in Suppl. Table 4.

We quantified the effect of each genetic perturbation on all measured transcripts using pseudo-bulk differential expression analysis in each condition. For perturbations with sufficient coverage, we tested for differential expression (DE) relative to NTCs, regressing out donor effects and cell count per perturbation (Suppl. Figure 4A, see Methods). We were able to estimate transcriptome-wide effects of perturbation of 11,527 genes in at least one condition (with 96% of them tested in all conditions). Of these, 7,807 (67%) perturbed genes had significant differential expression (FDR < 10%) effects on at least 3 genes in at least one condition. Across conditions, we discovered 2,035,311 significant regulator-to-gene

*trans*-effects. Throughout the manuscript, we use the term “regulator” to indicate a perturbed gene with significant *trans*-effects on at least one measured gene.

Perturbation effects were generally consistent between guides targeting the same gene (median cross-guide correlation: Rest = 0.49, Stim48hr = 0.52, Stim8hr = 0.47), with most discrepancies attributable to differential knockdown efficiency between guides (Suppl. Figure 4D) or to one guide producing substantially larger downstream transcriptional effects (Suppl. Figure 4E). To estimate the prevalence of off-target effects on proximal genes, we examined transcription start sites near guide targeting sequences and found that downregulation was most pronounced within 5 kb (observed in 60% of the 1,782 genes with proximal TSSs, 18% of all tested perturbed genes) and to a lesser extent within 5-10 kb (Suppl. Figure 4C). We flagged these cases as putative off-targets, where transcriptional responses may reflect knockdown of both the intended target and the proximal gene. Notably, some perturbations induced transcriptional responses without significant downregulation of the intended target (Suppl. Figure 5A), potentially reflecting either insufficient sensitivity to detect on-target effects or consistent off-target effects on distal genes. When testing for correlation between *trans*-effect estimates across donors, we observe transcriptome-wide concordance (Suppl. Figure 6), although we observe cases where *trans*-effects manifest only in a subset of donors (see Suppl. Note 1).

To further validate regulator-to-gene *trans-*effects genome-wide, we first tested whether transcriptome-wide effects for a given perturbed gene replicate across datasets. We compared the DE effect estimated by our perturb-seq screens with DE effects measured in published arrayed knockout screens in CD4+ T cells (using the Rest condition, to match the cellular context) [5,16]. We found that differential expression effects were significantly correlated between perturb-seq and independent arrayed screens for matched targets, with correlations significantly higher than for randomly paired perturbations (27 of 32 tested targets showed significant correlation) (Figure 1E). Non-replicating *trans*-effects may reflect noise in differential expression estimates due to limited cell numbers or small effect sizes (for example, IL2RA), or genuine differences between knockdown and knockout responses under distinct culture conditions (for example, NFAT5, TNFAIP3). We next compared genome-wide regulator effects on individual genes with measurements from FACS-based CRISPRi screens targeting four T cell-relevant genes in both resting and stimulated conditions [6,13,17]. Regulator effects were consistent with FACS-based screen estimates (Figure 1F), with significantly stronger correlations between matched stimulation conditions than between mismatched conditions. Overall biological reproducibility was remarkable given differences in perturbation techniques and phenotypic readouts. These results indicate that our dataset captures condition-specific regulatory effects on target gene programs across T cell activation states.

Overall, the median number of trans effects per perturbation with significant knockdown was 2 genes, while the top 5% of perturbations affect > 427 genes (Suppl. Figure 5B, mean = 81.61 genes), reflecting both the sparsity of regulatory networks and the presence of master regulators [18]. The mean number of trans effects can vary drastically between different classes of genes (Suppl. Figure 5C). The number of detected incoming trans-regulatory connections per measured gene was strongly dependent on baseline expression levels, at least partially reflecting power issues in differential expression analysis (Suppl. Figure 5D). After accounting for this expression-dependent detection bias, we found that trans-regulatory effects were most prevalent 8 hours post-stimulation, with immune-relevant gene sets, including cytokines and cytokine receptors, exhibiting more incoming regulatory connections than expected (Suppl. Figure 5E).

We tested whether trans effects identified in CD4+ T cells could be recapitulated in a different cell type, by comparing our results to published genome-wide perturb-seq data from K562 erythroleukemia cells [9]. After accounting for statistical power differences through meta-analysis (see Methods), approximately 8% of trans-effects detected in CD4+ T cells across 3,081 shared perturbations were also observed in K562 cells (Suppl. Figure 7A). For perturbations with measurable trans-effects in K562 cells (n = 1880), the DE effect correlation of the same perturbation in K562 cells or CD4+T cell states was significantly higher than that of random pairs of perturbations (Mean Pearson R = 0.32), but lower than the correlation between donor replicates (Suppl. Figure 7B), indicating partial sharing of regulatory effects across cell types. Perturbed genes showing the highest trans-effect correlation between K562 and CD4+ T cells were enriched for housekeeping cellular processes, including transcription, chromatin remodeling, telomere maintenance, and cell cycle regulation (Suppl. Figure 7C), which are expected to function similarly across cell types. These results demonstrate that even broadly expressed genes can exhibit substantially different downstream regulatory effects across cellular contexts, highlighting the need for comprehensive perturb-seq profiling in multiple cell types to fully map context-dependent regulatory networks. Summary statistics on perturbation effects in each condition are reported in Suppl. Table 5.

To our knowledge, no existing perturb-seq dataset provides comparable depth, perturbation coverage, and replication structure for robustly estimating genome-wide regulator-to-gene effects using methods that account for biological and technical variance [19]. To guide the design of future perturb-seq screens in primary cells, we conducted a power analysis examining how cell number, donor replicates, and read depth affect trans-effect estimation accuracy (Methods). We quantified this by measuring correlations between differential expression estimates in downsampled datasets and either held-out cells (Suppl. Figure 8A-B) or independent arrayed knockout screens (Suppl. Figure 8C-D). Downsampling read depth to approximately 10% (approximately 1,000 UMIs per cell) substantially reduced trans-effect replication, even with >500 cells per perturbation. At higher read depths, increasing cell numbers per perturbation consistently improved differential expression estimates (Figure 1G, Suppl. Figure 8), though estimates stabilized at approximately 200 cells per perturbation. Importantly, for a fixed number of cells per perturbation, increasing the number of donor replicates consistently led to more robust trans-effect estimates (Figure 1G, Suppl. Figure 8), likely by better capturing donor-specific response variation. These findings underscore the importance of cell coverage per perturbation and importantly, replication across biological replicates when performing large-scale perturb-seq studies in human primary cells.

### Identifying regulators of immune cytokines

As a first application of the new data set, we tested whether we could correctly identify the genes that regulate immune cytokines. Production of specific sets of cytokines is a major mechanism by which CD4+ T cells orchestrate immune responses. We and others have dedicated considerable efforts to identify genes that regulate production of key cytokines in human T cells to gain insight into immune regulation [20–22]. Our previous CRISPR screening studies have provided comprehensive insights into regulation of cytokines including IL-2 and IFN-γ [6] using FACS-based CRISPR screens, but this approach required massive sorting efforts to uncover the regulators of each individual cytokine of interest. Cytokine regulators discovered with our perturb-seq data strongly accord with FACS-based screen results despite differences in the experimental methods and readouts (notably mRNA vs. protein levels of cytokines, Figure 1F). The information-rich nature of genome-scale perturb-seq should enable simultaneous identification of regulators for a broad set of immune cytokines without the bottleneck of FACS-sorting large numbers of perturbed cells.

We examined perturbation effects on transcript levels of 30 cytokine genes, including pro-inflammatory cytokines (IFN-γ, TNF), Th2 cytokines (IL-4/5/13), regulatory cytokines (IL-10, TGF-β), chemokines (CCL3/4/5, CXCL8), and others. This analysis identified 1,556 gene perturbations that exerted strong regulatory effects (FDR < 1%) on at least one cytokine in at least one condition (Figure 2A, Suppl. Figure 10). Comparison of the Rest and Stim8hr conditions revealed several key patterns. We identified regulators that controlled a large set of cytokines, including expected TCR signaling components and subunits of the ubiquitous transcriptional coactivator complexes Mediator (MED12, MED24) and SAGA (TAF6L, TADA1, TADA2B, SUPT20H, USP22). Consistent with our previous findings [13], knockdown of Mediator and SAGA components appeared to alter T cell rest and activation and caused stimulation-specific effects on multiple cytokines, including TNF and IL-16 (Figure 2A-B). We also found regulators that controlled more limited subsets of cytokines. For stimulation-responsive cytokines like IL-2 and IL-13, negative regulators (knockdown log2FC > 0) were detected predominantly in the Rest condition, where cytokines exhibit minimal constitutive expression. In contrast, positive regulators (knockdown log2FC < 0) were more readily identified following re-stimulation (e.g., CD3 and LCP2 for IL2; GATA3 for IL13) (Figure 2A, Suppl. Figure 10).

**Figure 2.**
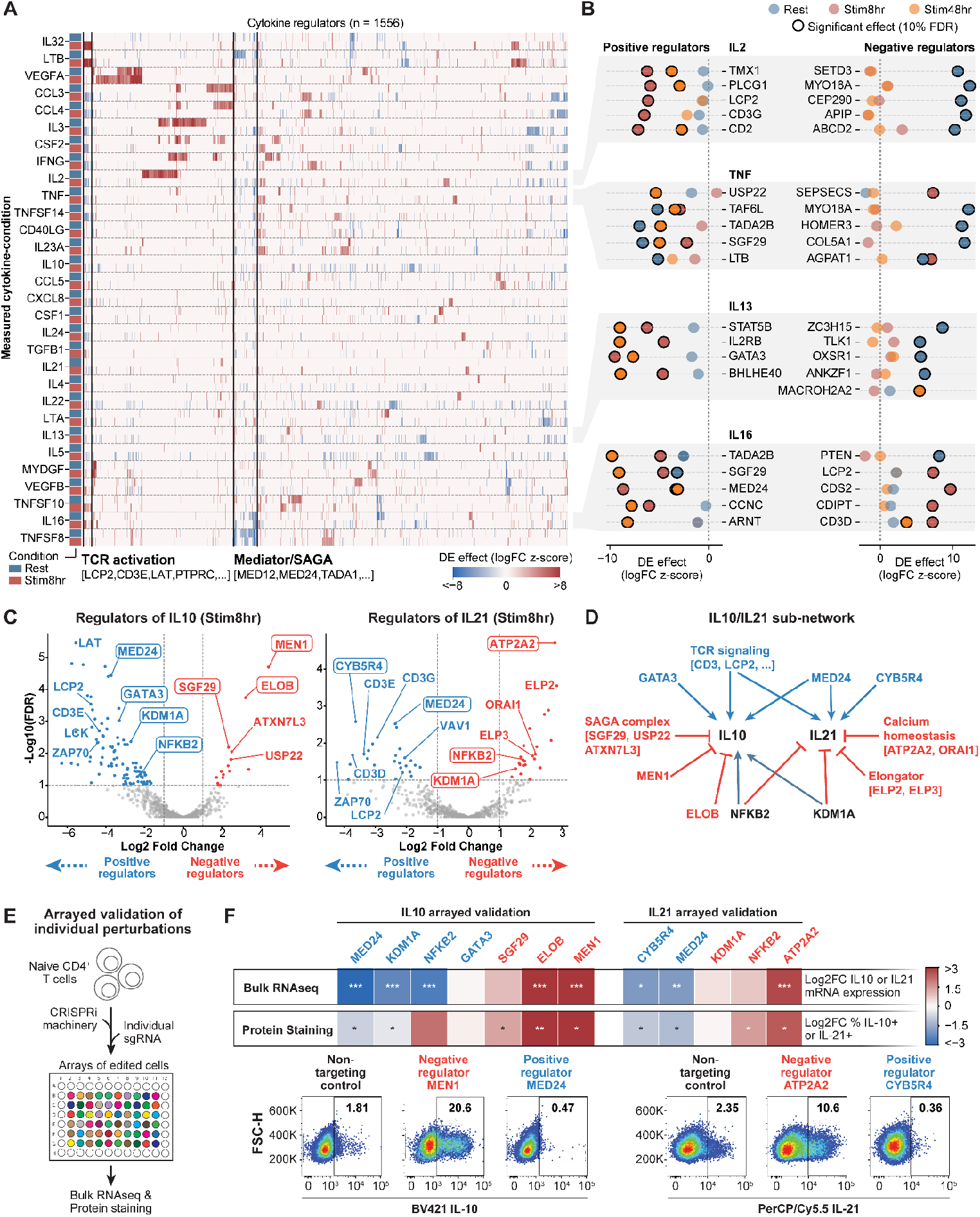
Mapping regulators of cytokines in primary CD4+ T cells. (**A**) Heatmap of DE effects (LFC z-score) of 1556 regulators with at least one significant trans-effect on expression of 30 cytokines (y-axis) in Rest and Stim8hr culture conditions (left color bar). Only significant DE effects (10% FDR) are shown. Axes are sorted by hierarchical clustering on Stim8hr DE effects. (**B**) Scatter plots of top 5 regulators (by z-score across conditions) of selected cytokines (IL16, TNF, IL13, IL2). For each cytokine, positive regulators (left) and negative regulators (right) are shown. Color: culture condition. Circled points: significant DE effects. **(C)** Volcano plots of log2 fold change of IL10/IL21 expression relative to NTCs and −log10 FDR for each regulator that has minimum cross-donor correlation > 0.35 (Suppl. Table 5). Significant regulators (FDR < 10%, |Log2FC| > 1) are colored blue (positive regulators) or red (negative regulators). Regulators validated in (F) are highlighted by white boxes. **(D)** Schematic of inferred IL10/IL21 regulatory sub-network. Connectors denote significant (10% FDR) downregulation (blue arrow) or upregulation (red T-bar) of IL10/IL21 by regulator knockdown. **(E)** Experimental workflow schematic for arrayed validation (also see Methods). **(F) (Top)** Heatmaps summarizing regulator knockdown effects in validation experiments. Regulator names are color-coded the same as (C). * FDR < 5%, ** FDR < 1%, *** FDR < 0.1%. **(Bottom)** Representative flow cytometry plots showing intracellular staining of IL-10 (left) and IL-21 (right) for NTCs and selected regulator knock-downs. Numbers in the gates indicate the percentage of cytokine-positive cells.

We focused on mapping and validating regulators of two cytokine genes, IL10 and IL21, which play pivotal, yet distinct, roles in immune homeostasis. IL10 is a potent anti-inflammatory cytokine gene with pleiotropic functions in suppressing excessive immune responses [23]. Defects in IL10 signaling are genetically linked to early-onset inflammatory bowel disease in both humans and mice [24,25]. Conversely, IL21 is essential for T cell proliferation and serves as a key mediator of cell-dependent B cell differentiation in germinal centers, with signaling defects often manifesting as primary immunodeficiencies [26–29].

Our perturb-seq measurements successfully identified a broad network of regulators for both cytokines, recovering both established and putative drivers of cytokine expression (Figure 2C). For example, MEN1, a component of the MLL1/2 complex known to repress IL10 expression [30], was among the top negative regulators of IL10, validating the sensitivity of the perturb-seq. Beyond shared drivers like core genes of the TCR signaling pathway and Mediator complex subunit MED24, perturb-seq revealed distinct regulatory architectures for each cytokine (Figure 2D). We observed negative regulation of IL10 by members of the SAGA complex (SGF29, ATXN7L3, USP22) and ELOB, a regulatory subunit of the Elongin complex. In parallel, perturb-seq identified oxidoreductase CYB5R4 as a positive regulator of IL21 and regulators of calcium homeostasis (ATP2A2, ORAI1) and the Elongator complex (ELP2, ELP3) as negative regulators of IL21. Notably, we found the transcription factor NFKB2 and the lysine-specific demethylase KDM1A as divergent regulators – acting as positive regulators for IL10 but negative regulators for IL21 (Figure 2C-D). Importantly, other than MEN1 and components of TCR signaling pathway being established players, the remaining regulators identified represent novel findings.

We selected 9 known and candidate regulators nominated by perturb-seq and knocked down each individually in an arrayed format, measuring outcomes via both bulk RNAseq and intracellular protein staining followed by flow cytometry (Figure 2E). The arrayed validation results demonstrated high concordance with the pooled perturb-seq data (Figure 2F, Suppl. Figure 11, Suppl. Figure 12). In transcriptional validation, 11 of the 12 tested regulatory interactions showed directional effects consistent with the screen, 8 of which were statistically significant. This concordance generally extended to the protein level, where flow cytometry confirmed significant shifts in protein production corresponding to the transcriptional changes. A notable exception was observed in the regulation of IL10 by NFKB2, where protein level changes did not mirror the transcriptional phenotype, suggesting additional post-transcriptional mechanisms that regulate cytokine production [31]. Together, these data demonstrate the power of genome-scale perturb-seq to dissect the complex regulatory circuitry of immune cytokines critical for immune homeostasis.

### Functional clustering of T cell regulators

Having shown that we can correctly identify known and novel regulators of cytokines, we next aimed to provide a more comprehensive characterization of CD4+ T cell functional programs. To do this, we clustered perturbations based on their effects on downstream genes, as previous work has shown that genes with similar perturbation profiles frequently have similar functions [9]. We focused on 3341 strong perturbations (defined as a gene knockdown in a specific stimulation condition) from 1860 perturbed regulators (Methods, Figure 3A). Unsupervised clustering based on shared transcriptome effects identified 111 regulator clusters (Suppl. Table 9). We annotated these regulator clusters using gene set enrichment analysis of public databases combined with manual curation (Methods). We identified clusters governing core cellular processes, including cellular signaling, intracellular trafficking, nonsense-mediated decay, cellular metabolism, genome integrity, and transcriptional regulation (Figure 3B, Suppl. Table 9). Many clusters could not be readily annotated based on known biological processes. Regulators in these clusters tended to be expressed in a tissue-specific manner (Suppl. Figure 13), which may explain their under-representation in public biological interaction databases [32].

**Figure 3.**
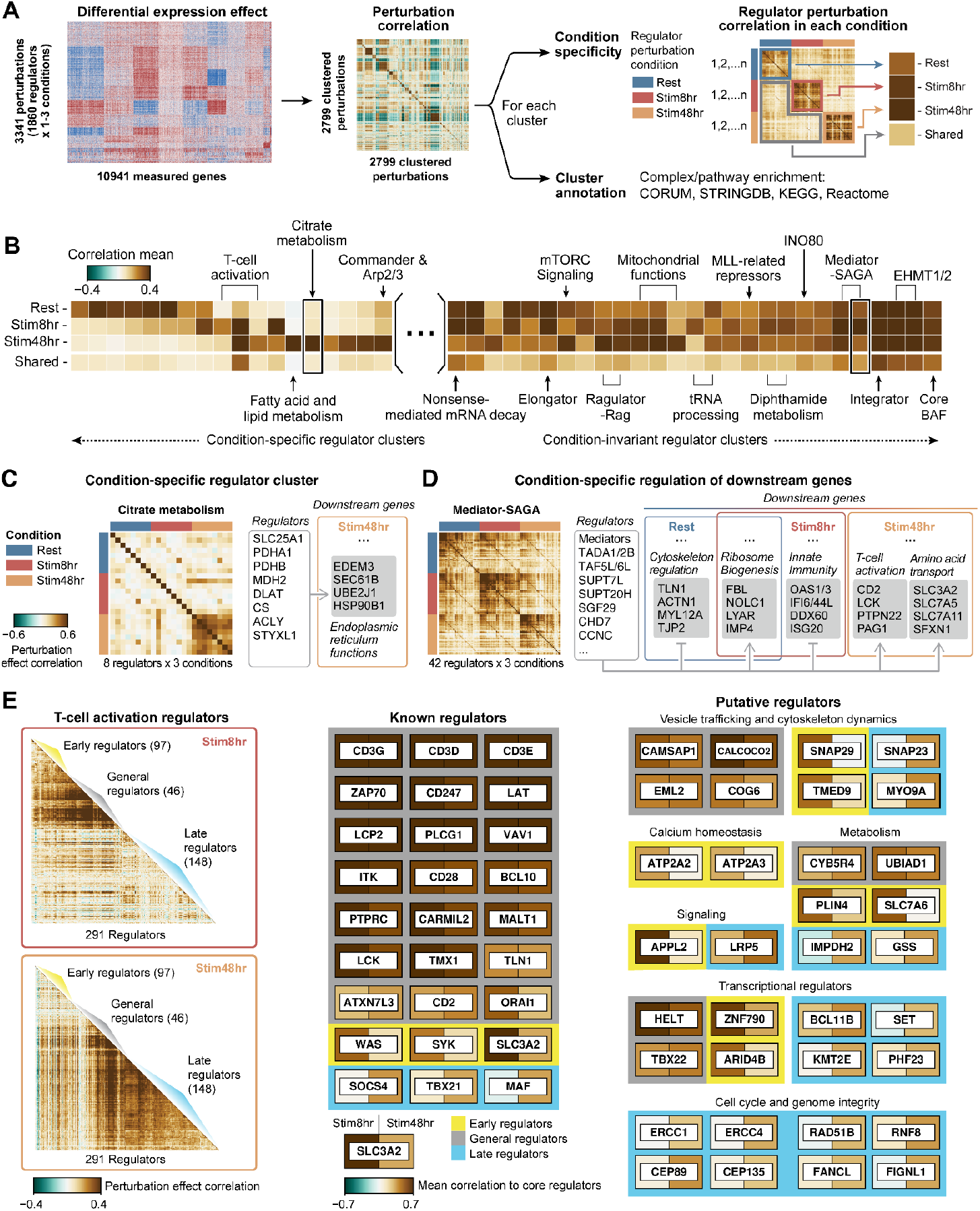
Functional gene programs of CD4+ T cells across conditions. (**A**) Schematic of clustering analysis on 3341 strong perturbations (one perturbation: one perturbed gene in one condition). Each cluster was analyzed for condition specificity and annotated by complex/pathway enrichment. The example correlation matrix shown on the right is from Cluster 28 (mitochondrial functions). (**B**) Condition specificity in a representative set of regulator clusters: heatmap showing mean intra-condition perturbation correlation of cluster regulators in each of the three conditions (“Rest”, “Stim8hr”, “Stim48hr”) and mean inter-condition perturbation correlation (“Shared”). Biological annotations of a subset of clusters are shown. (**C**) An example of condition-specific regulator cluster (Cluster 36). (**D**) An example regulator cluster showing condition-specific regulation of downstream genes (Cluster 10). In both (C) and (D), left heatmaps show the correlation perturbation effects for the same set of clustered regulators across all conditions. Representative downstream genes grouped by gene ontology analysis are shown on the right (Suppl. Figure 15). Connectors denote downregulation (arrow) or upregulation (T-bar) of downstream genes by regulator knockdown. (**E**) (left) Correlation matrix of T cell activation regulators. The rows and columns are ordered by early, general and late regulators. (right) Selected known and putative T cell activation regulators. Tiles are colored by the gene’s mean perturbation correlation with 9 core regulators (CD3D/E/G, CD247, ZAP70, LAT, LCP2, PLCG1, VAV1), in Stim8hr or Stim48hr.

Given that individual gene perturbations often elicit distinct effects depending on the stimulation context (Figure 2A), we assessed whether these regulator clusters exhibited condition specificity. For each cluster, we computed the intra-condition mean regulator perturbation correlation within each of the three conditions (“Rest”, “Stim8hr”, “Stim48hr”), as well as inter-condition mean perturbation correlation (“Shared”) (Figure 3A, top right, Methods). We observed two kinds of condition-specific clusters: 1) clusters of regulators that only exhibited correlated effects under specific conditions, and 2) clusters of regulators that had correlated effects within each of the three conditions, but controlled different sets of downstream target genes in different contexts.

Firstly, about one third of regulator clusters had strong intra-condition perturbation effect correlation in only one or two of the three conditions (Figure 3B, left, Suppl. Table 9). This condition-specific coherence of regulatory effects was not driven by regulator expression levels in the different conditions (Suppl. Figure 14A). These clusters represent regulatory complexes or pathways that only function in certain stimulation conditions. Regulators of citrate metabolism, for example, exhibited correlated regulator perturbation effects exclusively in the Stim48hr condition (Figure 3C), aligning with their established roles in metabolic regulation during immune activation [33]. Notably, we found many clusters without annotated biological functions fell in this category of condition specificity (Suppl. Figure 13).

Secondly, some regulators with correlated effects in multiple conditions regulated distinct sets of genes in different stimulation conditions, as shown by lower inter-condition correlations of perturbation effects (Figure 3B, right, the “shared” row). For instance, the regulators of Mediator-SAGA clusters showed overall high correlated perturbation effects within each of the three conditions, consistent with the regulators acting as a physical complex in all contexts. However, disrupting members of this complex of regulators altered the expression of distinct gene sets depending on the stimulation state of the cell (Figure 3D, Suppl. Figure 15, Suppl. Table 10), consistent with our previous findings [13].

As one noteworthy example of how these condition-specific effects manifest in core immune processes, we examined the three regulator clusters identified by unsupervised clustering that are associated with T cell activation (Figure 3B, 3E). These clusters, active primarily in the Stim8hr and Stim48hr conditions, encompass the majority of canonical activation regulators, including core CD3 receptor complex members (CD3D/E/G, CD247), stimulatory receptors (CD2, CD28, PTPRC), adaptor proteins (LAT, LCP2), and signal transduction enzymes (ZAP70, LCK, ITK, VAV1, PLCG1) [34]. TMX1 was also identified as a critical regulator of T cell activation, consistent with a recent report [35] that identified it via proximity labeling of CD8alpha and demonstrated that it is required for TCR surface expression.

We tested whether we could distinguish these regulators of T cell activation in those with early versus late effects. By leveraging perturbation measurements across the two stimulation time points, we dissected the regulators’ temporal roles. Regulators were considered “early regulators” if they mapped to these clusters at the Stim8h timepoint only, “late regulators” if mapped to these clusters at the Stim48hr timepoint only, or “general regulators” of activation if they mapped to these clusters at both stimulation timepoints (but not at Rest) (Figure 3E, Suppl. Table 9). Early regulators included calcium homeostasis factors (e.g. ATP2A2, ATP2A3), consistent with key roles of calcium signaling following T cell receptor engagement [36]. In contrast, late regulators include factors controlling cell cycle and genome integrity (ERCC1/4, CEP89/135, RNF8, RAD51B, FANCL, FIGNL1), highlighting the importance of these processes in supporting rapid cellular expansion. We also identified other putative early, late or general regulators involved in intracellular vesicle trafficking, cytoskeletal dynamics, central metabolism, cell signaling, and transcriptional regulation (Figure 3E, right). In addition, we observed distinct downstream functional programs that were regulated by early vs. late regulators of activation.

Eight hours post-stimulation, perturbations of early and general regulators induced changes of early activation markers including IL2RA and LAG3 (Suppl. Figure 14). In contrast, at 48 hours post-stimulation, perturbations of late and general regulators induced changes of many metabolic genes, particularly those driving oxidative phosphorylation (Suppl. Figure 14), consistent metabolic switches accompanying T cell activation [37].

Taken together, genome-scale perturb-seq in primary human cells performed in multiple stimulation conditions can identify clusters of regulators that act coherently to shape general and, importantly, context-specific gene expression programs.

### Predicting regulators of human T cell states: Th1 vs Th2 polarization

We next asked whether we can use the new perturb-seq data sets to interpret external data from human cohorts. Over the past decade, international efforts have generated single-cell atlases of cellular states across human tissues in health and disease. We hypothesized that perturb-seq maps in primary cells offer a unique opportunity to identify regulators driving the transcriptional programs observed in these atlases. If successful, perturb-seq could be used to identify causal factors responsible for establishing or maintaining tissue cell states, to elucidate how they emerge, and to guide their programmable control. Furthermore, regulators driving disease-associated or treatment-responsive states could serve as therapeutic targets. Therefore, we sought to test whether perturb-seq can be leveraged to identify regulators and pathways that contribute to T cell state signatures observed in human tissues.

Given the gene expression profile of a T cell state of interest, obtained from bulk or scRNA-seq experiments, we can compute a “state signature” by contrasting the expression profile of the state of interest with a reference T cell population. We hypothesized that knowledge of context-specific gene programs controlled by every potential regulator could be used to nominate a set of regulators that, together, move cells towards a desired cell state. Using differential expression estimates for each perturbed gene in perturb-seq data, we can fit a regression model to reconstruct the state signature as a linear combination of perturbation signatures (Figure 4A, see Methods). With this model, we can nominate putative positive and negative regulators of the cell state signature. Furthermore, we assume that each regulator may only control a subset of genes in the state signature and we can use perturbation effects to partition the state signature into sets of genes controlled by distinct regulators or pathways.

**Figure 4.**
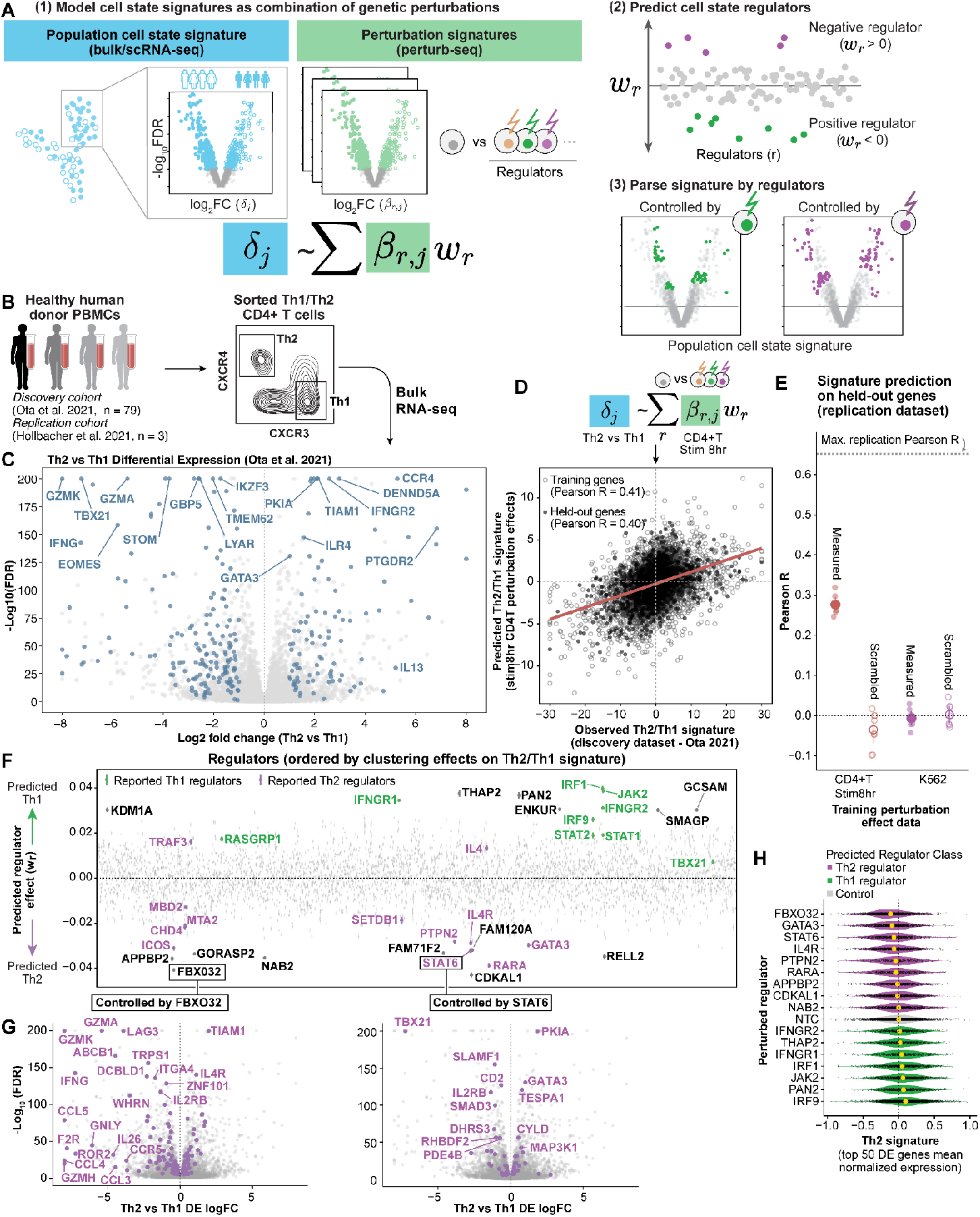
Predicting regulators of CD4+ T cell polarization. **(A)** Model schematic: cell state signatures from observational (bulk or single-cell) RNA-seq data are modeled as linear combinations of perturbation effects to identify regulators and parse their contributions to gene expression programs. **(B)** Schematic illustration of public data used to extract Th2/Th1 polarization signatures with bulk-RNA seq of sorted populations. **(C)** Volcano plot of Th2/Th1 signature (discovery cohort). Log-fold change (x-axis, LFC < 0: high in Th1; LFC >0: high in Th2) vs −log10 FDR (y-axis) of DE analysis (FDR values capped at −log10(FDR) = 200; LFC capped between −8 and 8). Robust genes significantly upregulated in both discovery and replication cohorts are highlighted in blue. Top robust DE genes by FDR are annotated. **(D)** Reconstruction of polarization signatures from perturbation effects: observed Th2/Th1 signature (x-axis) vs predicted signature from the model trained on the discovery cohort (y-axis). Black dots denote genes held-out in model fitting. **(E)** Evaluation of polarization signature prediction on held-out genes in the replication cohort: mean Pearson correlation coefficients (y-axis) across splits (5-fold cross-validation with 3 initializations) for model fit on Stim8hr CD4+T perturbation effects (red, 3994 regulators) and K562 perturbation effects dataset (blue, 2190 regulators). Scrambled controls are shown in light colors. Dotted line indicates maximum achievable correlation (inter-cohort agreement). Error bars: 95% CI across cross-validation splits. **(F)** Predicted regulator effects (w_r_, y-axis) on polarization signature for all regulators in Stim8hr condition (mean and standard error for w_r_ on 15 train-test splits). Regulators are ordered by hierarchical clustering on the perturbation effects on Th2/Th1 signature genes. Known Th1 and Th2 regulators highlighted in green and purple, respectively. Top and bottom regulators without known effects on polarization are annotated in black. **(G)** Th2/Th1 signature genes controlled by predicted regulator pathways: volcano plot of DE genes between Th2 and Th1 cells as in C, highlighting genes significantly regulated by FBXO32 (left) or STAT6 (right) (**H**) Distributions of Th2 signature gene expression (x-axis) in single cells with perturbation of top predicted polarization regulators (y-axis), ordered by mean signature score across cells (yellow dot). Color: predicted polarization effect (purple: Th2-promoting, green: Th1-promoting, gray: non-targeting controls). Signatures of NTC cells are shown only for cells with a subsample of non-targeting guides.

We first tested this approach to identify regulators of Th1 and Th2 polarized helper CD4+ T cells, cell states that each have well-characterized master regulators and play critical roles in immunity and inflammatory diseases [38]. We derived a Th2/Th1 signature from published bulk RNA-seq data of Th1 and Th2 cells FACS-sorted from human donors (Figure 4B) [39,40], and verified the signature’s robustness across independent cohorts (Suppl. Figure 16A). Significantly differentially expressed genes included well-characterized markers, such as *GATA3, CCR4*, and *IL13* upregulated in Th2 cells, and *IFNG, EOMES*, and *TBX21* upregulated in Th1 cells (Figure 4C).

We applied our model to reconstruct the Th2/Th1 signature from perturbation effects in the Stim8hr condition, because T cell polarization occurs upon T cell stimulation (Suppl. Figure 16B; results in different input conditions are discussed in Suppl. Note 2). Perturbation responses accurately predicted the sign and magnitude of held-out genes in the Th2/Th1 signature, both in the training cohort (mean cross-validation R = 0.39, Figure 4D) and an independent validation cohort (mean cross-validation R = 0.27, Figure 4E), demonstrating that predicting regulators from perturbation effects on one set of genes can predict polarization effects on different genes. Reconstruction from CD4+ T cell perturbations significantly outperformed models trained on K562 cell line data (Figure 4E, Suppl. Figure 16C), even when restricting the analysis to well-powered perturbed genes in the K562 data (Suppl. Figure 16D). We fit control models with scrambled perturbation effects for each gene to confirm that performance differences reflected genuine signal rather than baseline expression similarity (Figure 4E). Together, these results indicate that cell-type-specific perturbation responses can reconstruct gene expression signatures of T cell states observed in human populations.

The model nominated both known and novel regulators of Th1 and Th2 polarization. Top regulators promoting a Th1 state included: genes encoding for IFN-gamma receptor encoding genes (IFNGR1, IFNGR2); JAK2 (a signaling kinase downstream of the IFN-gamma receptor) [41] and the IFN-gamma-induced transcription factor IRF1 [42] (Figure 4F, Suppl. Figure 16E). Top Th2 regulators included: IL4R; STAT6 (a transcription factor activated by IL4R signaling) and GATA3 (a master transcription factor induced by STAT6); and the Retinoic Acid receptor (RARA), a transcription factor known to regulate Th2 cytokines [43,44]. The regulators identified by this analysis appear to be specific to Th1 and Th2 signatures as opposed to general activation, as different regulators were nominated when we fit an analogous model to predict a general TCR activation signature (Suppl. Figure 16F-G).

Examination of model weights across a broader set of known regulators confirmed they were generally highly ranked with the expected directional effects (Figure 4F). We found only 2 regulators whose inferred direction was opposite expectations: TRAF3, which is reported as a STAT6 regulator skewing cells toward Th2 fate, but only in specific contexts [45], and IL4. As with other cytokine perturbations, IL4 knockdown showed relatively weak transcriptional effects, likely due to paracrine signaling between cells in the pooled experimental set-up dampening the perturbation response. The model also prioritized several putative regulators without established roles in T cell polarization. Some genes, including KDM1A, SMAGP, and RELL2, have known functions in T cells but no reported effects on Th1/Th2 differentiation. Others, including CDKAL1 [46,47] and PAN2, have been implicated in autoimmune disease pathogenesis through genetic studies; or associated with eosinophil counts, a Th2-dependent phenotype, GCSAM (source: OpenTargets [48]). Notably, FBXO32, a component of the SCF E3 ubiquitin ligase complex, was predicted among the top Th2 regulators. While expressed in T cells, FBXO32’s role in immunity remains unstudied. FBXO32 activates NF-κB signaling during genotoxic stress and inflammation by stabilizing IκBα for polyubiquitination and proteasomal degradation [49], and has been associated with inflammatory responses in myocarditis [50]. Given that NF-κB is essential for GATA3 expression [51,52], FBXO32 knockdown may disrupt Th2 polarization through impaired NF-κB signaling.

The regulators differed in their effects on Th1 and Th2 programs. For example, FBXO32 and STAT6 knockdowns both promoted the Th1 signature while downregulating Th2 genes (Suppl. Figure 17A). In contrast, GATA3 knockdown downregulated Th2 genes but had minimal effect on Th1 signature genes. Moreover, some regulators with similar average effects across signature genes were found to control distinct pathways (Figure 4G). For example, FBXO32 and STAT6 controlled distinct target genes: FBXO32-controlled genes were enriched for chemokines (CCL4, CCL5, CCL3, CCR5; FDR = 0.04) and cytokines (IFNG, IL26, IL15; FDR = 0.07), whereas STAT6 regulated GATA3 and TGF-β pathway components (SMAD3, SMAD2, TGFBR2, LRRC32; FDR = 0.04).

Lastly, we examined the effect of putative regulators on Th1 or Th2 polarization at the single cell level in the perturb-seq data (Figure 4H, Suppl. Figure 17A, see Methods). The population of perturbed CD4+ T cells was not cultured in polarizing conditions, and we observed substantial cell state heterogeneity both among control cells and among cells with each perturbation. Even with heterogeneity of initial cell states and potential heterogeneity of knock-down efficiencies (Suppl. Figure 17B), perturbed cells showed the expected shifts in Th1 and Th2 gene expression compared to non-targeting controls. The subtle magnitude of these shifts likely reflects that full polarization requires the coordinated action of multiple regulators, in concert with external signal input.

Collectively, both pseudobulk and single-cell transcriptome show that regulators nominated by perturb-seq contribute to Th1/Th2 polarization. Together, these analyses illustrate how perturb-seq can identify regulators of cell state signatures and reveal the distinct molecular programs controlled by each regulator.

### Predicting regulators of CD4+ T cell states in population-scale scRNA-seq across age groups

While T cell states like Th1 and Th2 have been extensively characterized through isolation and phenotyping, recent single-cell RNA-seq studies are now revealing immune cell variation across diverse contexts including tissue-specific programs [53,54], disease-associated states [55–57], infection responses [58,59] and population-level variation [60–62]. We therefore wanted to test whether genetic perturbations could recapitulate transcriptional variation in T cells from human cohort studies, enabling us to identify candidate regulators of observed T cell gene signatures.

As a case study, we analyzed age-associated CD4+ T cell variation, where alterations are thought to contribute to chronic inflammation and multi-organ aging phenotypes [63]. We extracted CD4+ T cell expression profiles from 782 donors in the OneK1K cohort [60], holding out cells from 199 donors for validation. We tested for age-associated gene expression changes across CD4+ T cell subsets (naive, central memory, and effector memory) (Figure 5A) while controlling for subset-specific differences to avoid confounding by changes in cell composition associated with aging (Suppl. Figure 19A, see Methods). The age-associated gene signature replicated well in held-out donors and cell type-specific aging signatures from independent cohorts [64,65] (Suppl. Figure 19B), and showed no significant enrichment for genes associated with cytomegalovirus (CMV)-driven variation, a potential confounding factor across individuals of different ages [64] (Suppl. Figure 19C). Age-associated changes include upregulation of proinflammatory factors and downregulation of apoptotic pathways (Suppl. Figure 19D).

**Figure 5.**
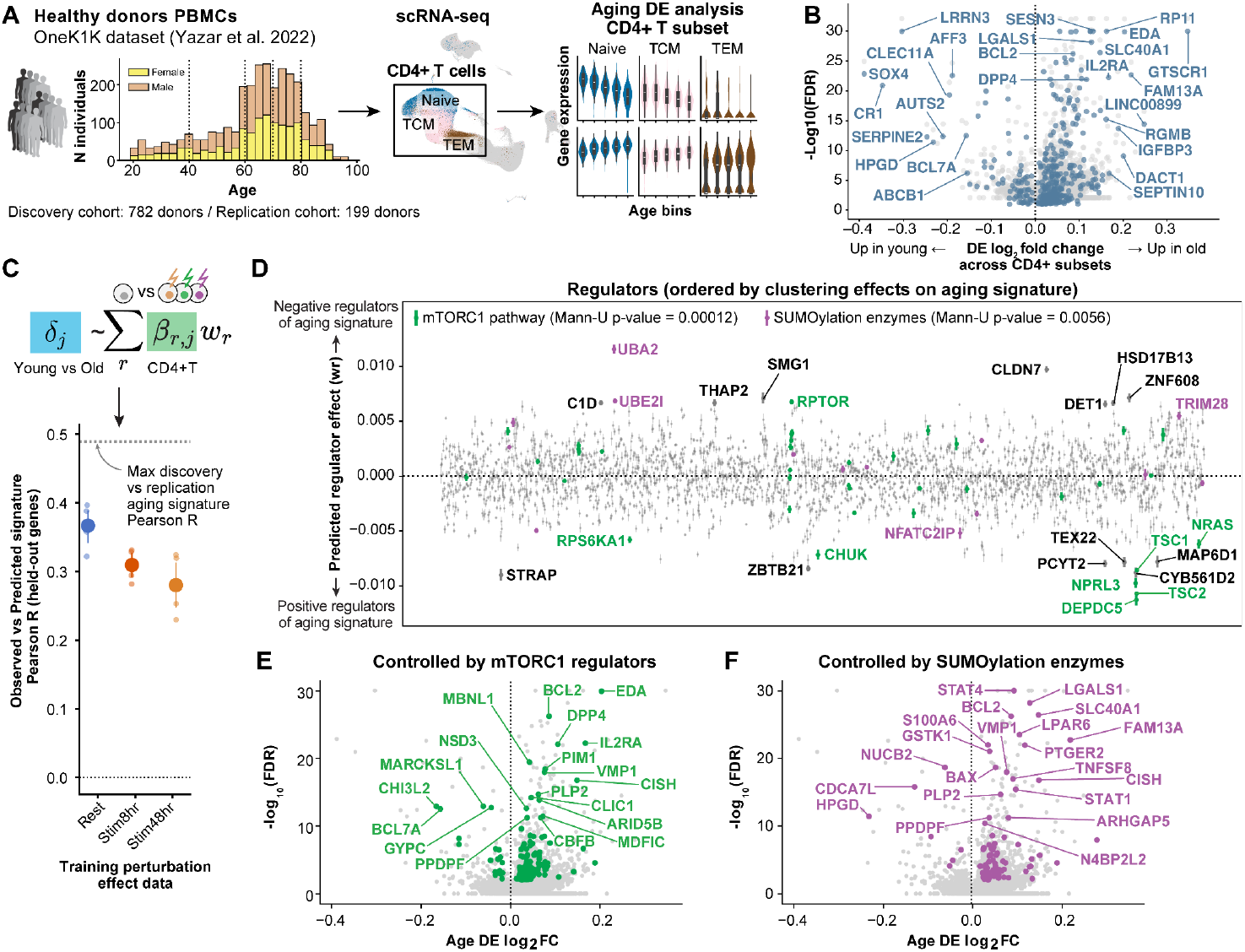
Predicting candidate regulators of CD4+ T cell aging from a population-scale cell atlas. **(A)** Illustration of analysis of OneK1K cohort scRNA-seq data for aging gene signature. The signature was computed by differential expression analysis across CD4+T cell subsets, testing for DE with age across subsets, accounting for subset differences. **(B)** Volcano plot of CD4+T cell aging signature (discovery cohort). Log-fold change (x-axis, LFC < 0: high in young; LFC >0: high in old) vs −log10 FDR (y-axis) of DE analysis (FDR values capped at −log10(FDR) = 30. Robust genes significantly upregulated in both discovery and replication cohorts are highlighted in blue. Top robust DE genes by LFC are annotated. **(C)** Evaluation of aging gene signature prediction on held-out genes: mean Pearson correlation coefficients (y-axis) across splits (5-fold cross-validation) for model fit on CD4+T perturbation effects in different conditions. Dotted line indicates maximum correlation between signatures in discovery and replication cohort. Error bars: 95% CI across cross-validation splits. **(D)** Predicted regulator effects (*w*_*r*_, y-axis) on aging gene signature for all regulators in Rest condition (mean and standard error for on *w*_*r*_ 5 train-test splits). Regulators are ordered by hierarchical clustering on the perturbation effects on aging signature genes. Perturbed regulators involved in mTORC1 signaling pathway (source: KEGG) or protein SUMOylation (source: Gene Ontology) are highlighted. (**E-F**) Aging signature genes controlled by predicted regulator pathways (see Methods): volcano plot of genes differentially expressed with age as in B, but highlighting genes significantly regulated by mTORC1 regulators (in E: including TSC1, TSC2, DEPDC5, NPRL3 and RPTOR) or SUMOylation enzymes (in F: including UBA1 and UBE2I)

We first tested whether genetic perturbation effects could recapitulate the aging signature. Perturbation effects in the Rest condition were predictive of age-associated changes (cross-validation R = 0.366), with significantly better fit than perturbation effects in stimulated CD4+T conditions or in K562 cells (Figure 5C, Suppl. Figure 20A-B). This likely reflects the predominance of unstimulated T cells in PBMC samples. These results demonstrate that genetic perturbations in a limited number of donors can nominate regulators of transcriptional changes observed across individuals.

We then examined which regulators are predicted to serve as positive or negative regulators of age-associated changes (Figure 5D). The first class of genes standing out among the top aging regulators were components of the mTORC1 signaling pathway, notably TSC complex genes (TSC1, TSC2), GATOR complex genes (DEPDC5, NPRL3), and RPTOR (Figure 5D). Hyperactivation of mTOR is considered a hallmark of aging: treatment with the mTOR-inhibitor rapamycin extends longevity in mammals [66] and pharmacological mTOR inhibition has been shown to improve immune function in the elderly [67,68]. In T cells, mTOR serves as a signaling node to activate several downstream effector pathways, including immune receptor signaling, metabolic programs, and migratory activity [69], and aged T cells exhibit increased basal activation of PI3K–AKT–mTOR signaling [63]. In line with this evidence, our model prioritized mTORC1 signaling genes as key regulators of age-associated phenotypes. However, the direction of effect of different mTORC1 components on aging is opposite to what we would expect: mTORC1 inhibiting factors are predicted as positive regulators of aging signature (i.e., knock-down of TSC and GATOR complex genes induces genes that are downregulated in aged CD4+ T cells), while RPTOR is predicted as a negative aging regulator, although the measured transcriptional effects of individual knock-downs have the expected effects on gold standard signatures of mTORC1 activation (Suppl. Figure 21A). We hypothesize that this could be due to negative feedback, compensation or survivor bias in the T cells profiled from aged individuals (Suppl. Note 3). Despite this complexity, the findings underscore mTOR signaling as a critical node of an observed T cell aging signature.

SUMOylation enzymes were also nominated as regulators of the T cell aging signature: these include *UBA2*, which encodes the E1 ubiquitin ligase for protein SUMOylation, and *UBE2I*, which encodes the E2 ligase UBC9. SUMOylation is a reversible post-translational modification in which Small Ubiquitin-like Modifier (SUMO) proteins are covalently attached to target proteins, regulating their localization, stability, and activity. This modification has critical effects on transcriptional regulation [70] and has been increasingly recognized as a regulator of aging processes [71–73]. In T cells, UBC9-mediated SUMOylation plays essential roles in peripheral CD4+ T-cell proliferation and homeostasis, including SUMOylation of PDPK, which regulates glycolysis-dependent T cell function [74,75]. Perturbing either mTOR signaling components or key SUMOylation genes had marked effects on the age-associated T cell signature, but perturbations in these two pathways affected distinct subsets of the overall signature. mTORC1 regulators control several genes associated with resistance to apoptosis that are overexpressed in aged T cells, including *BCL2*, PIM kinases, *ARID5B*, and *SERPINB6*, as well as genes involved in T cell activation and survival such as *DPP4* [76], *IL2RB, KMT2A* [16], *KLF3*, and *TCF7* (Figure 5E, Suppl. Figure 21C). SUMOylation enzymes control expression of STAT genes (*STAT1, STAT4, STAT6*), which are known substrates of SUMOylation [77–79]. These enzymes also regulate several cytoskeletal and structural genes, including *TIMP1, VIM, ITGB1*, and *FLNB* (Figure 5F, Suppl. Figure 21D). Interestingly, the effects of these regulators on these processes appear to be specific to the Rest condition and are not detected post-stimulation (Suppl. Figure 21C-D).

Collectively, we demonstrate that genetic perturbations in a limited number of donors can recapitulate transcriptional changes observed across individuals in population-scale cohorts. Moreover, while perturbing mTORC1 components and SUMOylation enzymes altered the overall age-associated signature, these regulators do not primarily affect the top differentially expressed genes in aging (Suppl. Figure 20C-D), highlighting the value of perturb-seq analysis of signature effects over screening approaches focused on isolated markers to identify major regulators. In summary, this analysis enables prediction of key pathways that shape population-level changes and partition signatures based on genes controlled by specific regulatory pathways.

### Interpreting natural genetic variant effects on immune traits with perturb-seq

As a final application, we sought to use the *ex vivo* perturb-seq data to identify the regulatory pathways through which natural genetic variants influence complex immune traits. Interpreting genetic associations with complex traits remains challenging, in part because many genes detected in genome-wide association studies (GWAS) exert their primary effects on a trait through *trans*-regulation of other genes [80,81]. But perturb-seq can potentially allow us to decipher these regulatory relationships, and to identify the key transcriptional programs that impact phenotypic variation. Indeed, recent studies have shown that CRISPR perturbations of selected genes can be used to identify specific transcriptional programs that are enriched for GWAS hits [3,5,16,82,83]. Since CD4+ T cells are central to many immune-mediated diseases and complex traits [84,85], we hypothesized that the new perturbation data could provide a powerful interpretive tool for immune traits.

We tested this using human lymphocyte counts measured in the UK Biobank [86] as a model T cell-relevant trait. Following recent work from our group [3], we focused on the phenotypic effects of rare coding loss-of-function (LoF) variants, which can be used to directly quantify the magnitude and direction by which dosage of a single gene affects organism-level traits [86]. We reasoned that if a gene is a “core gene” for lymphocyte count (i.e., exerting a direct causal effect on lymphocyte count [80]), then the major regulators of that gene, as identified by perturb-seq, should also show genetic signals in the human LoF data. Specifically, to identify putative core genes, we tested for each gene *j* whether there is a significant correlation between the knockdown effects of regulators on *j* and the LoF effects of those regulators on lymphocyte count in the UK Biobank. We refer to the correlation between LoF effects and knockdown effects for each putative core gene as the *regulator-burden correlation* (Figure 6A).

**Figure 6.**
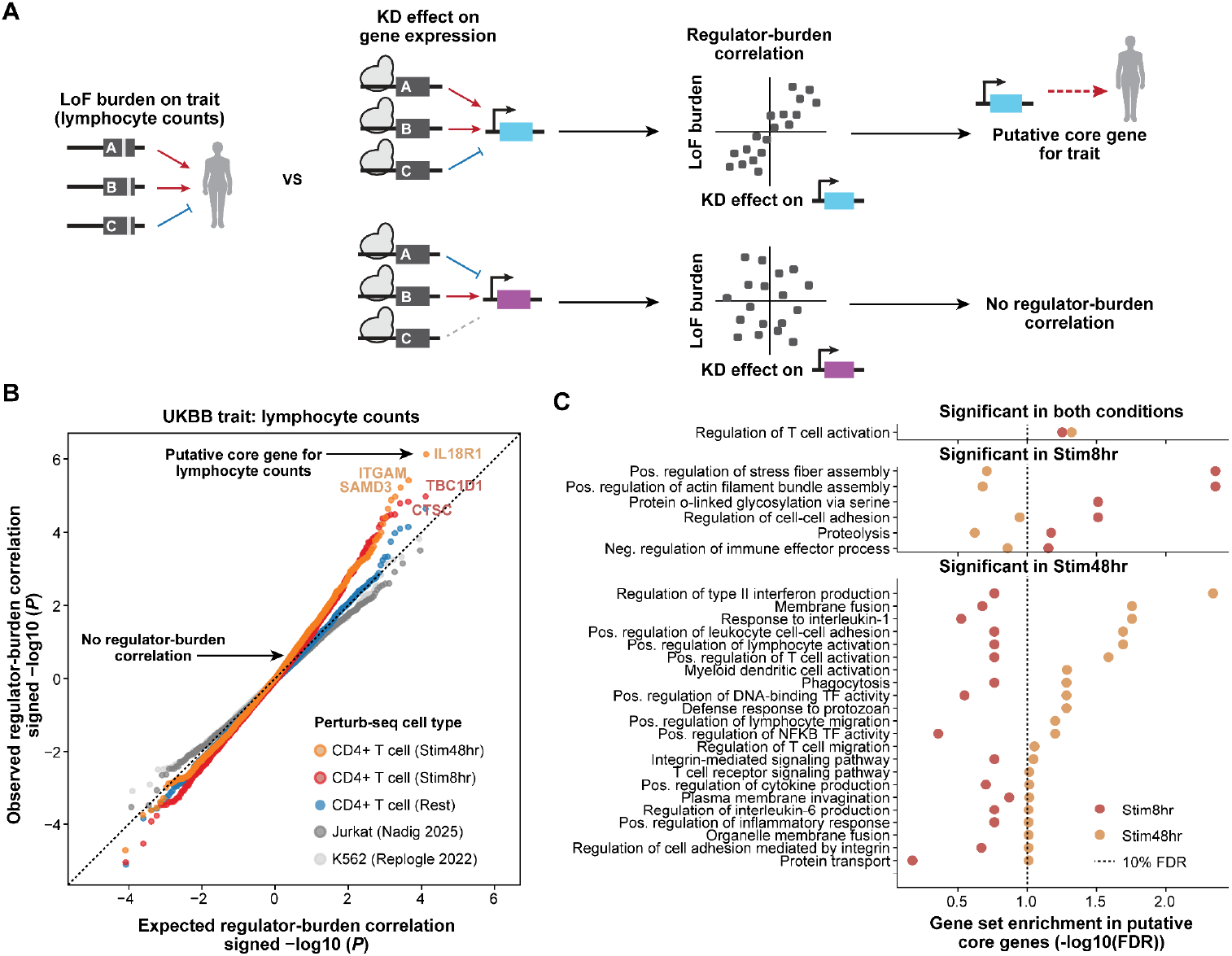
Context-specific regulatory effects in perturb-seq explain genetic association signals on lymphocyte counts. **(A)** Schematic of regulator-burden correlation analysis: we compare directional gene loss-of-function (LoF) burden estimates on complex traits (based on UK biobank [UKBB] annotation of human variants and traits) with regulatory effects on each measured gene in perturb-seq. The correlation between the two estimates is used to nominate putative “core” genes affecting the complex trait. **(B)** Signed Q-Q plot comparing expected vs observed signed p-values for regulator-burden correlations for LoF burden on lymphocyte counts in the UKBB and regulatory effects estimated from perturb-seq data in different cell types/conditions (color). Sign denotes direction of correlation. Each dot represents one measured gene. Perturb-seq data from this study is colored, while perturb-seq data on control cell types is in shades of grey. **(C)** Gene Ontology enrichment of putative core genes positively associated with lymphocyte counts (top 50 by signed −log10 p-value reg-burden correlation) in Stim8hr (red) and Stim48hr (orange) conditions. X-axis: FDR from hypergeometric test. Top: terms enriched in both conditions; middle: Stim8hr only; bottom: Stim48hr only. Missing points indicate no overlap between core genes and gene set.

Analysis of regulator-burden correlations in stimulated T cells indicated an excess of significant correlations at both timepoints. In contrast, we did not find higher than expected correlations in either the rested T cells, K562 cells or in a screen of essential genes in Jurkat cells (an immortalized T cell line) (Figure 6B), suggesting that these other cell states and types are poor proxies for modeling lymphocyte count regulation. The difference in enrichment between perturbations in Jurkats versus primary T cells could also be explained by the larger number of genetic perturbations measured in our perturb-seq data (Suppl. Figure 22A). However, the different correlations in genome-scale perturb-seq in stimulated T cells compared to those in rested T cells or K562 cells indicate that both the number of gene knockdowns and the specific cellular context matter for recapitulating gene effects on complex traits.

We next characterized the putative core genes with high regulator-burden correlations identified in the stimulation timepoints. Among the top putative core genes in the Stim8hr condition was ADAM19, a metalloprotease involved in cell-cell communication. ADAM19 is regulated by high burden genes in the Stim8hr condition (including KLF2, NDFIP2, NSD1, PLCG1, ERGIC1), which do not have a regulatory effect at the later time point (Suppl. Figure 22D). In Stim48hr condition, core genes included multiple subunits of the IL-18 receptor (IL18R1 and IL18RAP), which is known to induce proliferation of lymphocytes, including NK cells and activated T cells [87] (Suppl. Figure 22D). Notably, MYD88, the signaling adapter downstream of the IL-18 receptor, was one of the strongest positive hits from LoF burden analysis on lymphocyte counts. While MYD88 knockdown produced only weak effects in our perturb-seq data (< 3 DE genes in any condition, despite significant knock-down), the regulator-burden correlation analysis nonetheless converged on IL-18 signaling as a pathway driving this trait, highlighting the power of this integrative approach.

Core genes identified in the different conditions were significantly correlated (Pearson R = 0.36), but several top putative core genes were condition-specific (Suppl. Figure 22B). Gene set enrichment analysis on the top genes positively associated with lymphocyte counts showed that in both conditions putative core genes are enriched in T cell activation markers (Figure 6C). However, genes prioritized in different conditions mapped to different biological processes: core genes at the 8 hour timepoint were enriched in actin fiber assembly (including PXN, S100A10, and GPR65) and cell-cell adhesion genes (including ADAM8, ADAM19, and DPP4). Core genes identified at 48 hours post-stimulation were enriched in lymphocyte activation pathways and regulation of cytokine production. These results indicate that measuring perturbation responses in different conditions allows us to detect different aspects of complex trait biology.

Having shown that perturb-seq provides a useful model for studying lymphocyte count, we next characterized genes associated with autoimmune disease risk. Genetic association studies have identified hundreds of loci linked to autoimmune diseases, yet translating these findings into actionable therapeutics requires understanding the causal genes, the cell types in which they exert their effects, and the precise nature of their molecular effects. To address these questions, we investigated whether our clusters of perturbed regulators (Figure 3, Suppl. Table 9) affecting shared downstream gene programs in T cells were enriched for autoimmune disease-associated genes. We aimed to characterize shared regulatory effects and assign putative immunological functions to genes with established genetic associations but unknown roles in T cell biology. We annotated a set of genes associated with 14 autoimmune conditions (hereafter *autoimmune genes*) from Open Targets [48] (Suppl. Figure 23A). We then tested the odds of enrichment of these autoimmune genes in each cluster of regulators and among the top condition-specific genes downstream of these regulators (Figure 7A).

**Figure 7.**
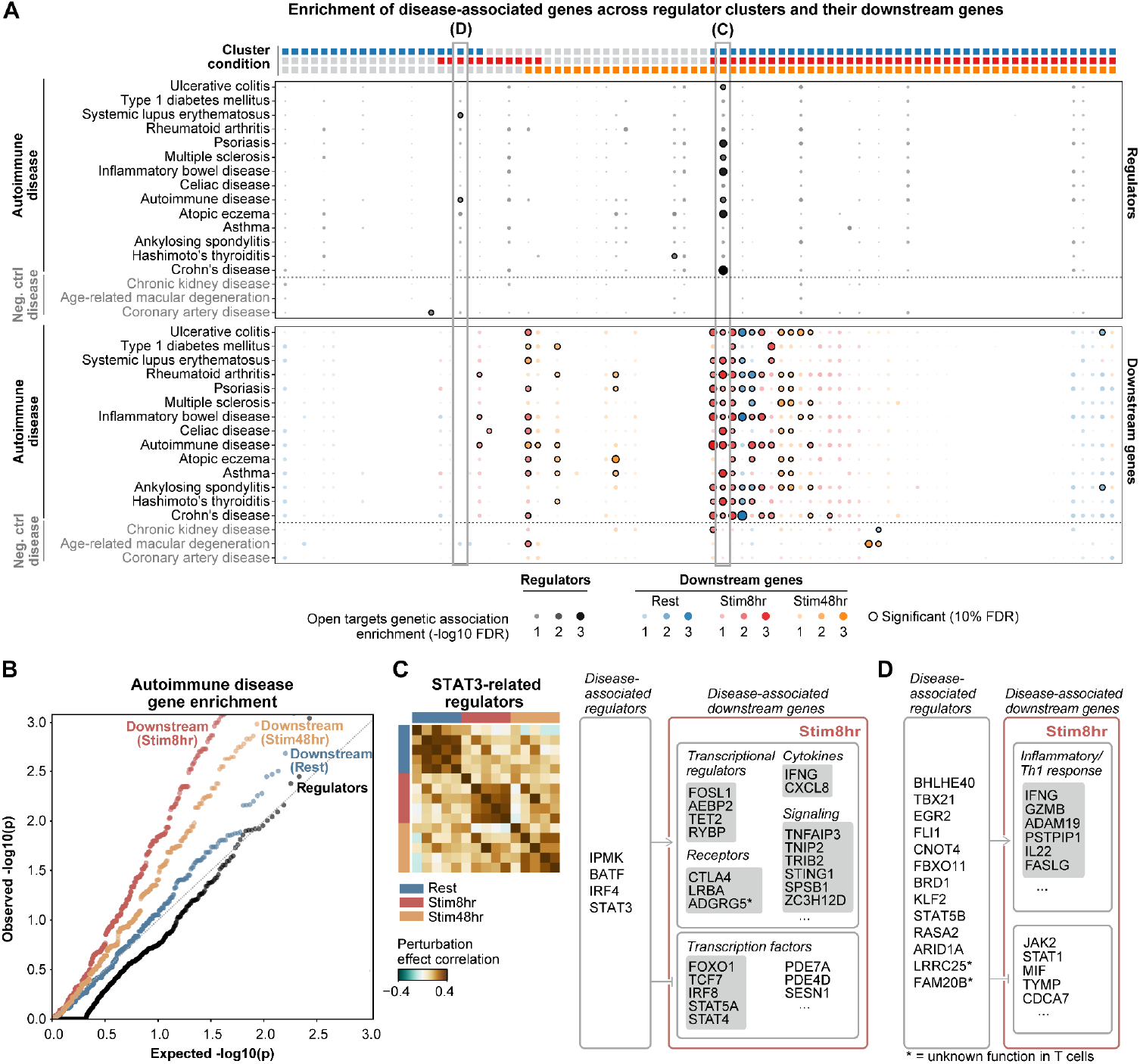
Regulation of autoimmune-associated genes. (**A**) Enrichment of GWAS-evidence genes (Open Targets) for autoimmune and control diseases (y-axis) in regulators (top) and downstream genes (bottom) per cluster (x-axis). Clusters with at least one disease-associated gene are shown. Dot size and color scale are proportional to the −log10(FDR) of the fisher exact test for enrichment in GWAS-evidence genes. Dots with a black outline indicate significant enrichment (10% FDR). Clusters are ordered by condition specificity, as annotated in the top bar. Clusters highlighted in follow-up vignettes are outlined with grey boxes. (**B**) Q-Q plot of p-values for enrichment in disease-associated genes across clusters in regulator genes (black) or downstream genes across conditions (colored). (**C-D**) Illustrations of clustered regulators and downstream genes with genetic evidence for autoimmune conditions from OpenTargets: (C) regulatory relationships in cluster 80 (STAT3-related regulators). Heatmap shows the correlation between perturbation effects of the 5 clustered regulators across conditions (as in Figure 3) (D) illustrates regulatory relationships in cluster 79.

We found significant enrichment of autoimmune genes in 33 of 77 tested clusters, among either regulators or condition-specific downstream genes. By contrast, we observed significantly weaker enrichment for genes associated with three non-immune diseases where CD4+ T cell contribution to pathology is expected to be minimal (Figure 7A, Suppl. Figure 23B). Among condition-specific regulator clusters, autoimmune genes showed significant overlap among clusters that show correlated regulator effects in Stim8hr and/or Stim48hr conditions, but not in clusters whose regulators only show correlated effects in Rest condition (Figure 7A, Suppl. Figure 24). Similarly, within clusters of regulators with correlated effects across conditions but distinct sets of downstream target genes in different conditions, we observed notable enrichment of autoimmune genes among downstream targets regulated in stimulated cells (Figure 7B, Suppl. Figure 24).

One such example is the STAT3 regulatory network. We identified a cluster of regulators enriched for autoimmune disease-associated genes encoding a core network of transcription factors involved in Th1/Th17 cell specification–STAT3, IRF4, BATF [88] and IPMK, which encodes inositol polyphosphate multikinase and was recently shown to control STAT3 phosphorylation via Akt/mTOR signaling in mice [89] (Figure 7C). Perturbations of these regulators had coordinated effects across conditions, but the set of co-regulated downstream genes changed across conditions, with enrichment for autoimmune genes observed only among downstream targets in the Stim8hr condition (Figure 7B, Suppl. Figure 24). This STAT3-related network positively regulated autoimmune disease-associated genes encoding pro-inflammatory cytokines (IFNG, CXCL8), signaling proteins (TNFAIP3, TNIP2, STING1, SPSB1), transcriptional regulators (FOSL1, Polycomb components), and immune receptors (CTLA4, LRBA). The network negatively regulated the transcription factor STAT4 and the phosphodiesterases PDE4A and PDE7A (which regulate intracellular cAMP levels) [90], as well as other key transcription factors linked to T cell regulation and memory (FOXO1, TCF7, IRF8, STAT5A). Several of these autoimmune disease-associated downstream genes showed minimal transcriptome-wide effects when individually perturbed (Suppl. Figure 25A). This could be due to low baseline expression or synergistic mechanisms, for example for the PDE4/7 phosphodiesterases [90]. Among downstream genes co-regulated by the STAT3 network in stimulated cells, we identified genes with autoimmune risk associations but limited functional characterization in T cells (Suppl. Figure 25B). These genes included the putative tumor suppressor ZC3H12D, which has been primarily studied in innate immunity [91], and the GPCR family receptor ADGRG5, which may also function in cAMP signaling [92].

While a major goal of human genetic studies is to discover new biology, prioritizing candidate genes and discovering their mechanisms of action in relevant cell types remain significant bottlenecks. To this end, we examined if correlated gene perturbation effects could link new genes implicated in disease risk to recognizable pathways in human T cells. For example, we identified a Stim8hr-specific cluster of regulators enriched for autoimmune disease-associated genes (FDR < 0.1), controlling expression of Th1 effector genes (IFNG, FASLG [93]) and other genes involved in T cell activation and function (Figure 7C). This cluster included known regulators of Th1 polarization (TBX21, BHLHE40, EGR2, FLI1), MHC-II regulators (CNOT4, FBXO11), and regulators of anti-tumor T cell functions (RASA2 [94], ARID1A [95]). Notably, clustering with these functionally characterized regulators were two autoimmune disease-associated genes with limited functional characterization in T cells: LRRC25 and FAM20B. LRRC25 encodes a membrane protein involved in type 1 IFN signaling [96], primarily described as a myeloid-specific receptor [97]. FAM20B encodes a kinase involved in proteoglycan metabolism [98] and is annotated as a likely causal gene for allergy and systemic lupus erythematosus in Open Targets, but remains largely unstudied in the immune system. FAM20B knock-down induced a substantial transcriptome-wide effect in the Stim8hr condition (Suppl. Figure 25C) that correlated with the effects of knocking down known Th1 regulators (Suppl. Figure 25D). This transcriptome-wide effect was consistent between guides (Pearson R = 0.81) and across donors (mean correlation = 0.50). Our data nominate LRRC25 and FAM20B as unappreciated regulators of T cell activation and Th1 polarization pathways, illustrating how guilt-by-perturbational association can nominate immunological roles for disease-implicated genes with previously unknown function in T cells. In summary, joint analysis of perturb-seq responses and human population genetic studies provides new insights into the functional roles of gene products harboring disease-associated variants, and the regulatory relationships between them.

## Discussion

High-content CRISPR-based perturbation screens promise to map genetic circuits at scale. However, no single study has yet combined genome-scale perturbation libraries across multiple primary cell contexts with deep transcriptional readouts, thus limiting our ability to robustly measure dynamic regulatory maps. We now leverage a CRISPR toolkit developed for primary T cells [5,6,13] and advances in scRNA-seq [14] to generate genome-scale perturb-seq maps of primary human T cells in multiple human donors and multiple stimulation timepoints. Our data serves as a foundational resource for studying regulators of cell state and function in the human immune system. We make the data publicly available to facilitate these efforts and accelerate community-driven discovery.

The cell coverage, single-cell transcript recovery, and biological replication in our dataset enable reliable quantitative estimates of each perturbation’s effect on individual genes throughout the transcriptome (Figure 1). Previous perturb-seq studies have been limited largely to analyzing the gene signature effects of each perturbation due to low cell coverage per perturbation or by sparse read depth of most transcripts [9,15,82,99] (Suppl. Figure 7!). While gene program signatures can provide a useful abstraction [3], here, quantification of gene-level effects enabled us to discover novel cytokine regulators (Figure 2) and query regulators of specific disease-associated genes (Figure 7). Moreover, gene-level summary statistics are readily comparable across datasets, facilitating meta-analysis of perturbation studies and integration with human cohort data (Figure 4, Figure 5). Performing biological replicates of each perturbation in hundreds of cells from multiple human donors was a critical factor enabling this level of resolution in measurements of perturbation effects.

These data bring us closer to the goal of building a complete map of GRNs, which would enable the systematic identification of circuit motifs – such as feed-forward loops and autoregulation – serving as the fundamental building blocks of cellular computation [100]. While such motifs have been extensively characterized in prokaryotes and yeast [101,102], revealing the circuit design principles of mammalian cells has remained challenging due to lack of high-resolution, genome-scale functional maps. Future advancements can integrate dosage-controlled perturbations [103] and dynamic readouts [104] to capture non-linear regulatory relationships and deconvolve direct regulatory interactions from indirect ones. Eventually, we envision these efforts could converge into dynamic and quantitative mathematical models that predict GRN behaviors underlying complex cellular computations.

Measuring responses across stimulation timepoints, we found striking evidence of widespread context-specific effects of gene perturbations. Previous work has shown cell type-specific transcriptional responses to the same genetic perturbation [105,106]. Here, even within closely related cell states, we observe substantial context specificity. Some perturbations exert strong effects in one condition but have no detectable impact in others. In other cases, the same genetic perturbation, or cluster of perturbations, produces significant effects across multiple conditions, but acting on distinct downstream gene programs (Figure 2), consistent with our previous work [13]. Additionally, different cellular contexts are relevant to distinct aspects of human biology. For example, perturbations in stimulated cells reveal the effects of canonical Th1/Th2 regulators (Figure 4), while responses in rested cells capture aging-associated transcriptional signatures (Figure 5). Similarly, when examining regulatory effects of genes with high loss-of-function burden on lymphocyte counts, both stimulation timepoints help to explain the signal through distinct downstream genes (Figure 6). Our observations demonstrate the value of measuring gene perturbation effects in diverse primary cells captured in multiple cellular contexts. We anticipate future studies will capture perturbation effects in primary T cells exposed to distinct modes of cytokine, TCR, costimulatory and metabolic signals *in vitro* and eventually *in vivo* [107].

We demonstrate new opportunities to integrate systematic gene perturbation studies in human primary cells with human genetic studies and atlases of human cells measured in healthy and diseased tissues. Experimentally-derived gene regulatory networks provide a framework to interpret the molecular effects of human genetic variants associated with diverse traits and diseases. In addition, we demonstrate that the regulators and gene programs mapped in our perturb-seq data provide insights into key pathways shaping cell states observed in human samples. Previous studies derived perturbation-based gene signatures and searched for *in vivo* cells expressing these signatures [105,108,109]. Others have proposed methods to identify which experimental perturbations best match the transcriptional profile of a query cell, with metric learning on large cell atlases [110] or, more recently, using a regression approach termed “RNA fingerprinting” that is conceptually similar to the approach we propose here [111]. Our framework differs in two key assumptions: first, that signatures of natural cell states arise from the combined effects of multiple regulators rather than a single best-matching perturbation; and second, that both positive and negative regulators contribute to observed expression changes. This enables us to partition cell state signatures into genes controlled by each predicted regulator or pathway. Modeling differential expression statistics controls for systematic differences between tissue and *ex vivo* cells, making this framework broadly applicable across conditions and cell states. While we employ a relatively simple model here, we view this as a proof-of-concept and a baseline for more sophisticated approaches, for example, models that directly incorporate context-dependent perturbation effects. Ultimately, we envision perturb-seq in primary cells as a critical bridge that will allow us to understand how genetic variation governing health and disease shapes molecular programs that operate in the cells comprising human tissues [112].

We anticipate that our dataset will be particularly valuable for developing machine learning models that predict molecular responses to genetic perturbations, a capability viewed as central to building AI Virtual Cells [113]. True biological replicates provide a rigorous framework for testing out-of-sample predictions and calibrating expectations for reproducibility in human biology. Furthermore, our genome-wide measurements across multiple cell states provide a foundation for developing models that predict perturbation effects across cellular contexts, which is a difficult, but essential task for prediction models. Existing models have had limited success when trained on cell line datasets with systematic biases and low depth [114–117] and struggle to accurately predict *trans*-effects of genetic perturbations across cell types [118,119].

Additionally, screens in primary cells facilitate integration with multi-cell type observational data, which could further guide generalization across contexts. However, our findings on context-specific responses also suggest that models may face substantial hurdles in predicting the effects of perturbations in contexts that were not directly tested. Accurate cross-context prediction may require large-scale data acquired from substantially more cellular contexts than are currently available.

Our study establishes a framework for systematic mapping of gene regulatory networks and genotype-phenotype relationships in primary human cells. Here we analyze the effect of gene knockdowns with CRISPRi, but the approach is readily adaptable for the ever-expanding set of perturbations that are achievable at scale, including new genetic perturbation modalities [6,120–123] and delivery methods to edit more difficult-to-transfect primary cell types [124,125]. Technologies are not only expanding the scope of perturbations, but are also facilitating more comprehensive readouts of perturbation effects: increasing the scale and reducing the per-cell cost of single-cell library preparation [126–128] and sequencing [129], coupling additional cellular measurements to each perturbation [107,130–132], and reading out perturbation effects in their native contexts [107]. As advanced perturbation and measurement technologies converge, we envision this framework extending to new perturbation modalities and cellular contexts, scaling from tens of millions to hundreds of millions, or even billions, of cells. This exponential leap in data generation will necessitate a parallel revolution in scalable computational methods and machine learning infrastructure to efficiently process, store and interpret these massive, high-dimensional regulatory maps.

Over the past two decades, human population genetics have associated genetic variation to human traits. More recently, single-cell genomic technologies have mapped diverse cell states in human health and disease. Now, perturb-seq in human primary cells has potential to map how genetic variation controls cell states, offering new hope that we can systematically link genome sequences to cell programs to human health outcomes. Our analyses showcase how this genome-scale perturb-seq map assembled in human primary CD4+ T cells across stimulation time points and multiple donors can be harnessed for insights into human immunology, immunotherapy design, dynamic gene regulatory control of human cells and human disease genetics.

## Limitations of current study

Despite performing large-scale perturb-seq with 2 gRNAs per target across multiple human blood donors, future work would still be required to fully assess potential off-target guide effects with additional guides per target gene and to differentiate donor-specific biology from experimental batch effects. Our analysis uses pseudobulk aggregation to compare mean expression profiles across cells with the same perturbation. This enhances statistical robustness, but potentially masks heterogeneity of response to perturbation across cells, which could be resolved by future distribution-aware analyses. Our study focuses on a non-polarized culture condition; consequently, the regulatory rewiring driven by polarizing cytokines – central to CD4+ T cell plasticity – remains unmapped. Future work extending this framework to include diverse *in vitro* polarization contexts and complex *in vivo* signals is essential to generate a comprehensive model of context-dependent immune regulation. Although perturb-seq elucidates causal gene regulatory relationships, future experimental and computational efforts will be required to link the transcription effects of each perturbation to specific immune cell functions in order to harness these data for immunology and immunotherapy development.

## Author contributions

R.Z., E.D., A.M. and J.K.P. conceived and designed the study. R.Z., J.Y., J.R.R., performed the perturb-seq experiments. E.D. and R.Z. performed all the data analyses. R.Z., J.R.R. and R.C.G. performed validation experiments. M.O. contributed to the design and interpretation of the analyses, and provided intellectual contributions to all aspects of the study. R.G. contributed to the data analysis on loss-of-function burden statistics. L.K.P. contributed the data analysis on aging T cells in OneK1K. R.Z., E.D., A.M. and J.K.P. wrote the manuscript, with critical input from all authors. J.K.P. and A.M. supervised the study and acquired funding.

## Acknowledgments

We thank T. Gjorgjieva, T. Zeng, S. Alatkar, J. Spence, R. Lopez, S. Dodgson, Z. Steinhart, M. Arce, B. Shy and all members of the Marson lab and Pritchard lab for helpful discussions on the project and for feedback on the manuscript. We additionally thank Z. Steinhart for generously sharing the EF1a-Zim-3-dCas9-P2A-BSD construct. We thank early inputs on 10x Flex from G. Xing, R. Stickels and T. Abay. This project has been made possible in part by grant number CZIF2025-011112 from the Chan Zuckerberg Initiative Foundation DAF, an advised fund of Silicon Valley Community Foundation. We thank B. Marshall and J. Cool from CZI for their support, as well as other grantees of the Billion Cell Project initiative for useful discussions. We thank I. Gadbank, J. Chien, J. Zamanian and all of the Lattice team for supporting data and metadata curation and sharing. In partnership with CZI, 10x Genomics, Ultima Genomics and Psomagen provided reagents and expertise on single-cell RNA sequencing. We also thank P. Lund, P. Smibert from 10x Genomics for early technical support on CRISPR protocol with 10x Flex, and J. Kim, J. Yeoh and D. Cooper for overall 10x related support. We thank T. Clark and the team at Ultima Genomics and J. Stone and the team at Psomagen for providing technical support on sequencing. We thank T. Tolpa for support on figure design. R. Zhu is the Connie and Bob Lurie Fellow of the Damon Runyon Cancer Research Foundation (DRG-2509-23). E. Dann is supported by a Helen Hay Whitney Foundation fellowship and by an EMBO non-stipendiary postdoctoral fellowship. A.M. received funding from the Simons Foundation, Parker Institute for Cancer Immunotherapy, the Weill Cancer Hub, K. Jordan and the CRISPR Cures for Cancer Initiative. This work was supported by the US National Institutes of Health (grants R01HG008140, R01HG014005, P01AI138962, R01DK129364, P01AI155393).

## Competing interest statement

A.M. is a cofounder of Site Tx, Arsenal Biosciences, and Survey Genomics, serves on the boards of directors at Site Tx, and Survey Genomics, is a member of the scientific advisory boards of network.bio, Site Tx, Arsenal Biosciences, Cellanome, Survey Genomics, NewLimit, Amgen, and Tenaya, owns stock in network.bio, Arsenal Biosciences, Site Tx, Cellanome, NewLimit, Survey Genomics, Tenaya and Lightcast and has received fees from network.bio, Site Tx, Arsenal Biosciences, Cellanome, Spotlight Therapeutics, NewLimit, Abbvie, Gilead, Pfizer, 23andMe, PACT Pharma, Juno Therapeutics, Tenaya, Lightcast, Trizell, Vertex, Merck, Amgen, Genentech, GLG, ClearView Healthcare, AlphaSights, Rupert Case Management, Bernstein and ALDA. A.M. is an investor in and informal advisor to Offline Ventures and a client of EPIQ. The Marson laboratory has received research support from Biohub/Chan Zuckerberg Initiative, the Parker Institute for Cancer Immunotherapy, the Emerson Collective, Arc Institute, Juno Therapeutics, Epinomics, Sanofi, GlaxoSmithKline, Gilead and Anthem and reagents from 10x, Ultima, Genscript, Illumina and Cellanome. The remaining authors declare no conflict of interest.

## Methods

### Genome-scale perturb-seq experimental methods

#### Genome-scale gRNA library design and library cloning

We compiled a list of genes to target by taking the union of all expressed genes in human CD4+ T cells and all transcription factors annotated in Lambert et al 2018 [133]. To find expressed genes in human CD4+ T cells across contexts and subsets, we used public gene expression datasets DICE (https://dice-database.org), ENCODE (https://www.encodeproject.org/) and Tabula Sapiens [134]. For each dataset, we selected a set of genes that accounts for >99% of total expression, then took the union of genes from three datasets and transcription factors from Lambert et al 2018, targeting 12,748 genes in total. We then selected two gRNAs per gene cross-referencing two widely-used genome-wide CRISPRi guide RNA databases, hCRISPRiv2 and Dolcetto [135,136], as well as hits from CRISPRi screening studies [6,137–139]. Specifically, for each gene, we selected gRNAs with the following criteria in descending priorities until two gRNAs are found: (1) gRNAs that are hits from existing CRISPRi screening studies in T cells [6] and other cell types [9,137–139]; (2) gRNAs that appear in both hCRISPRiv2 and Dolcetto; (3) gRNAs that are in Dolcetto dataset. We also included 5% of non-targeting gRNAs from the Dolcetto dataset in the library as controls. In total, our genome-scale gRNA library consists of 26,504 gRNAs (Suppl. Table 3).

#### Lentivirus production

The EF1a-Zim-3-dCas9-P2A-BSD [13] plasmid is a generous gift from Zachary Steinhart. Lentiviral particles were produced as previously described [6]. In brief, for a T225 flask, 3.6 × 10^7^ Lenti-X HEK 293T cells were seeded in 45 ml Opti-MEM I Reduced Serum Medium (OPTI-MEM) with GlutaMAX Supplement (Invitrogen, catalog no. 31985088) supplemented with 5% FCS, 1 mM sodium pyruvate (Fisher Scientific), and 1× MEM nonessential amino acids (Fisher Scientific) (cOPTI-MEM). The next morning, cells were transfected with 38 μg transfer plasmid, 28 μg psPAX2 (Addgene 12260) and 14 μg pMD2.G (Addgene 12259) with 167 µl p3000 and 181 µl Lipofectamine 3000 (Fisher Scientific, L3000075). Six hours after transfection, medium was replaced with fresh cOPTI-MEM supplemented with 1x viral boost (Alstem, VB100). Another 18 hours later, virus containing supernatant was collected and spun down at 500g for 5 minutes at 4 °C to pellet cell debris. Lentivirus-containing medium was moved to a new vessel and subsequently concentrated 100-fold using Lenti-X Concentrator (Takara Bio, 631232) per the manufacturer’s instructions. Viral particles were stored at −80 °C until further use.

#### Isolation and culture of naive human CD4^+^ T cells

Primary naive human CD4+ T cells were isolated from PBMC-enriched leukapheresis products (Leukopaks, STEMCELL Technologies, catalog no. #70500) from healthy donors following institutional review board–approved informed written consent (STEMCELL Technologies). The leukapheresis products were washed twice with a 1X volume of EasySep buffer (DPBS, 2% FBS and 1 mM EDTA, pH 8.0) using centrifugation. The washed cells were resuspended at 5 x 10^7^ cells/ml in EasySep buffer and isolated with the EasySep Human Naive CD4+ T Cell Isolation Kit (catalog no. #19555, STEMCELL Technologies), according to the manufacturer’s protocol. CD4+ T cells were cultured in complete X-VIVO 15 (cXVIVO) consisting of X-VIVO 15 (Lonza Bioscience, 04–418Q) supplemented with 5% heat-inactivated fetal calf serum (R&D systems).

#### Genome-scale perturb-seq

After isolation (day 0), 30 million naive human CD4^+^ T cells were seeded at 1 x 10^6^ cells/ml cXVIVO supplemented with 200 IU/ml of recombinant human IL-2 (Amerisource Bergen, catalog no. 10101641). Cells were activated using ImmunoCult human CD3/CD28/CD2 T cell activator

(10990, STEMCELL Technologies) at 25 µl/ml. The next morning, cells were transduced with 2% v/v concentrated ZIM3KRAB-dCas9-BlastR lentivirus. In the afternoon, cells were transduced with 0.1% v/v (multiplicity of infection ~ 0.2) genome-scale gRNA library lentivirus. At day 3, cells were centrifuged and resuspended with above medium supplemented with 200 IU/mL IL-2. Blasticidin and puromycin were added to 10 µg/mL and 2 µg/mL final concentrations, respectively. At day 5, cells were counted and diluted to 5 x 10^5^ cells/mL with cXVIVO supplemented with 200 IU/mL IL-2 and 10 µg/mL blasticidin. Recombinant human IL-7 (R&D Systems, catalog no. 207-IL) was also added to a final concentration of 5 ng/mL. At day 7, cells were counted and diluted to 7.5 x 10^5^ cells/mL with cXVIVO supplemented with 200 IU/mL IL-2, 5 ng/mL IL-7 and 10 µg/mL blasticidin. At day 8, cells were centrifuged and resuspended at 1 x 10^6^ cells/mL with cXVIVO supplemented with 200 IU/mL IL-2 and 5 ng/mL IL-7. At day 10, cells were centrifuged and resuspended at 5 x 10^5^ cells/mL and 1 x 10^6^ cells/mL with cXVIVO supplemented with 200 IU/mL IL-2 and 5 ng/mL IL-7. At day 12 morning, cells were centrifuged and resuspended at 1 x 10^6^ cells/ml with cXVIVO supplemented with 200 IU/mL IL-2 and split into three populations. The first population (Rest) was kept at a rested state without restimulation and harvested in the afternoon after 8 hours. The second population (Stim8hr) was restimulated by adding 12.5 µL/ml ImmunoCult human CD3/CD28/CD2 T cell activator and harvested in the afternoon after 8 hours. The third population (Stim48hr) was restimulated by adding 12.5 µL/ml ImmunoCult human CD3/CD28/CD2 T cell activator and harvested after 48 hours. Cells were harvested, fixed and stored using GEM-X Flex Sample Preparation v2 Kit (10x genomics, catalog no. 1000781) per the manufacturer’s instructions.

#### Single-cell library construction

Stored cells were processed to scRNA-seq and gRNA sequencing libraries using GEM-X Flex Gene Expression Human n-plex kit (10x genomics, catalog no. 1000829) for 16-plex probes and GEM-X gel beads and GEM-X Flex Gene Expression Human kits (catalog no. 1000792) for additional GEM-X gel beads, following the manufacturer’s protocol. Protocols were modified to enable CRISPR gRNA detection according to 10x’s technical note “CG000814, CRISPR Screening with GEM-X Flex Gene Expression”. Spike-in probes to detect gRNA expression were ordered from IDT as oPool. To assess the off-target gRNA detection by off-target probe binding, we also ordered 768 gRNA probes that detect gRNA that are not in our genome-scale library. Less than 0.2% of cells had an assignment of “guides” corresponding to one of out-of-library gRNA probes, demonstrating the low off-target detection of gRNA by probe-based methods. Probes to detect puromycin resistant (PuroR) gene expression were also ordered from IDT and were spiked in following 10x’s technical note “CG000621, Custom Probe Design for Visium Spatial Gene Expression and Chromium Single Cell Gene Expression Flex”. Briefly, 12 samples were processed in three batches.

- Batch 1: Rest - D1, Rest - D2, Stim8hr - D1, Stim8hr - D2.
- Batch 2: Rest - D3, Rest - D4, Stim8hr - D3, Stim8hr - D4.
- Batch 3: Stim48hr - D1, Stim48hr - D2, Stim48hr - D3, Stim48hr - D4.

Of the three batches, the second batch and the third batch were processed on the same days (see Suppl. Table 1). For each batch, we thawed, counted and put 8 million cells from each of the four biological samples into the hybridization step (32 million cells in total). The 8 million cells from each biological sample (e.g. Rest - D1) was resuspended with 640µL Hyb Mix and evenly distributed into 4 hybridization reaction tubes. Then we added WTA probes for gene expression, spike-in probes for gRNA expression and PuroR expression. In summary, we had 16 hybridization reaction tubes, with 4 hybridization reaction tubes per biological sample. Each hybridization reaction tube consisted of 2 million cells, 160µL Hyb Mix, 40µL WTA probes, 20nM CRISPR LHS probe, 2nM/probe CRISPR RHS probes, 2nM/probe PuroR LHS probes, 2nM/probe PuroR RHS probes with total volume slightly more than 200µL. We further split each ~200µL hybridization reaction into 4 aliquots, with each aliquot tube having ~50µL hybridization reaction before putting into thermocycler for ~19 hours overnight incubation (including a ~1.5hr 60C-42C ramp-down phase specific for CRISPR). The next day, we performed the post-hybridization wash protocol. Then we loaded:

1. Batch 1: 1.161M cells per lane for 23 lanes (~2.5x overloading)
2. Batch 2: 0.926M cells per lane for 24 lanes (~2x overloading)
3. Batch 3: 0.8M cells per lane for 24 lanes (~1.7x overloading)

We proceeded with library construction according to the manufacturer’s protocol (10x Genomics).

#### Sequencing

The final 10x libraries were converted into Ultima compatible libraries in 7 PCR cycles using conversion primers provided by Ultima Genomics. Converted libraries were purified with SPRISelect beads (Beckman Coulter) and quantified using Qubit 1X dsDNA High Sensitivity Assay kit (Thermo Fisher). Converted libraries were sent to Psomagen to sequence on Ultima sequencers. Each 10x lane was sequenced on one Ultima wafer with ~10B sequencing reads. The sequencing reads were processed by Psomagen into single cell gene expression count matrices using CellRanger on 10x Cloud services with following configuration setting (chemistry,auto; no-secondary,false; create-bam,false; Filter-probes,true) and our custom probe set. The custom probe set consists of Chromium Human Transcriptome Probe Set v1.1.0 plus two spike-in probes targeting the puromycin resistance gene associated with gRNA expression cassette. Spike-in probes are these two lines in the custom_probe_set.csv: CUSTOM001_PuroR,TCCGCGACCCACACCTTGCCGATGTCGAGCCCGACGCGCGTGAGGA AGAG,CUSTOM001_PuroR|PuroR|8aab555,TRUE,unspliced,PuroR CUSTOM001_PuroR,AACCACGCGGGCTCCTTGGGCCGGTGCGGCGCCAGGAGGCCTTCC ATCTG,CUSTOM001_PuroR|PuroR|8aab556,TRUE,unspliced,PuroR

#### Plasmids

ZIM3KRAB-dCas9-BlastR (pZR316) plasmid is a kind gift from Zachary Steinhart. lentiGuide-Puro-BlpI (PJY390, Addgene #251159) was cloned by Gibson assembly with lentiGuide-Puro plasmid (Addgene #52963) as the backbone. Backbones for single gRNA expression for arrayed experiments were designed by switching Puromycin resistance genes with GFP (pRZ109) or mCherry (pRZ111) and were cloned by GenScript. All gRNAs for validation experiments were cloned into pRZ109 or pRZ111 by GenScript. Sequences for all backbones used in this study and individual gRNA sequences used in validation experiments (for Figure 2) can be found in Suppl. Table 2.

Our genome-scale CRISPRi library (to be available on Addgene) was cloned into PJY390 by Golden Gate Cloning as described previously [136]. The oligonucleotide pool was obtained from Agilent and amplified with KAPA HiFi HotStart (Roche 07958935001). The PCR product was purified by MinElute PCR Purification Kit (Qiagen 28004) and ligated into lentiGuide-Puro-BlpI (PJY390) with the New England Biolabs NEB Golden Gate Assembly Kit (BsmBI-HF v2) (New England Biolabs, E1602L) per the manufacturer’s instructions. The ligation product was transformed into Endura electrocompetent cells (LGC, 60242-2) per the manufacturer’s instructions. Cells were expanded at 30 C for 22 h and plasmid was extracted using ZymoPURE II Plasmid Maxiprep Kit (Zymo, D4203). For library quality control, the guide RNA distribution was confirmed by NGS sequencing at coverage of >1,000× (raw fastq available through study GEO). The plasmid pool was then used for lentivirus production as described under ‘Lentivirus production’.

### Perturb-seq data preprocessing and quality control

After CellRanger analysis was performed, we processed in parallel mRNA and gRNA UMI count matrices for each sample and lane. We initially filtered out cells with >20% mitochondrial gene expression and < 200 detected genes.

We used the Poisson-Gaussian mixture model to assign guides to cells, following the model in Replogle et al. 2022, using the implementation in the *crispat* package [140] (https://github.com/velten-group/crispat). This models the distribution of UMIs across cells as a mixture of 2 distributions, one for background guide probe detection and one for true signal from cells where the guide is expressed. In our pilots, this approach outperformed using a fixed threshold to assign guides to cells excluding background (in terms of number of cells with multiple guides assigned and identification of guides with frequent non-specific binding). We fit a separate model for each guide in each processed sample, assuming there might be differences in mean UMI counts per guide across samples.

We then identified cells with no assigned gRNA, one assigned gRNA or multiple assigned gRNAs and the UMI counts of the most abundant guide in each cell.

After guide assignment, we excluded cells with low quality transcriptomes (>5% mitochondrial reads, <500 genes), ambiguous gRNA assignments (“multi_sgRNA”), or no detectable gRNA.

When reporting normalized expression values, raw counts were normalized per cell and log-transformed using default parameters implemented in *scanpy* [141,142]. To assess batch effects between lanes, we analyzed non-targeting control (NTC) cells across experimental conditions, subsampling 10 sequencing lanes per sample from all experimental batches, resulting in 395,030 cells. We performed dimensionality reduction with scVI, minimizing differences between donors with the following parameters: 30 latent dimensions, negative binomial gene likelihood, 2 hidden layers with 128 units, 0.2 dropout rate. This revealed no substantial batch effects between 10x lanes or experimental batches, with most transcriptional variance explained by culture condition and proliferation status (Suppl. Figure 2A-E). We observed expected expression patterns for canonical T cell activation markers across stimulation timepoints, although expression levels varied between donors, especially post-stimulation (Suppl. Figure 2F).

For a preliminary estimate of guide efficiency (Suppl. Figure 3), we calculated the expression of the targeted gene in cells with each guide and compared it to expression in NTC cells with a one-sided T-test, using the Benjamini-Hochberg correction to control the false discovery rate. The knockdown was considered significant if FDR < 10% and t-statistic < 0. Guides for which no significant knock-down was detected in any condition (with at least 10 cells and mean log-normalized expression in NTCs > 0.01) were considered inefficient and excluded from differential expression analysis (n=869). Summary statistics on target expression per condition are reported in Suppl. Table 4.

### Analysis of trans effects

#### Differential expression analysis

To estimate the average effect of each perturbed gene in each culture condition, we aggregated data from all cells within the same biological sample (donor and condition) harboring the same guide, by summing mRNA counts. We refer to each experimental unit of donor-condition-guide as a “pseudobulk” sample. We discard pseudobulks with data from less than 5 cells, and where the total mRNA count was much lower compared to other pseudobulk in the same conditions (< 0.5% percentile). Only perturbed genes represented by at least 3 pseudobulks across guides and donors were included in differential expression testing (N perturbed genes per condition: Rest = 11,289; Stim8hr = 11,417; Stim48hr = 11,333). For downstream gene selection in each condition, we first removed expression outliers (mean counts > 10,000 or < 1, and percentage of expressing pseudobulks > 99.9% or < 1%). We then retained the top 10,000 highly variable genes for differential expression analysis using the Scanpy function *scanpy*.*pp*.*highly_variable_genes* (flavor = ‘seurat_v3’) on pseudobulk expression profiles.

We performed differential expression (DE) analysis with the DESeq2 method [143], using the python implementation [144,145]. Briefly, the mRNA counts of gene *j* in pseudobulk *p* were modeled by a negative-binomial generalized linear model:

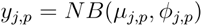

The expected count value *μ*_*j,p*_ is given by the following log-linear model:

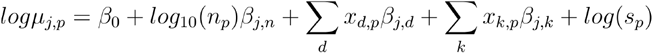

- *β*_0_ denotes the intercept
- *n*_*p*_ denotes the number of aggregated cells in pseudobulk *p*
- *x*_*d,p*_ denotes the assignment of pseudobulk *p* to donor *d*
- *x*_*k,p*_ denotes the assignment of pseudobulk *p* to perturbed gene *k*
- *β*_*j,n*_, *β*_*j,d*_and *β*_*j,k*_denote the effect of number of cells, donor and perturbation, respectively, on the mean expression of gene *j*
- *s*_*p*_denotes the library size (total UMI counts per pseudobulk)

For each perturbed gene, we estimated the log2 fold-change in gene expression and *β*_*j,k*_test for differential expression using the Wald test, contrasting expression estimates for each perturbed gene against non-targeting control pseudobulks. For each culture condition, we fit DE models independently, to obtain context-specific perturbation estimates 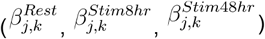. To account for differences in noise between perturbations with different number of recovered cells across replicates, unless otherwise specified we use the z-score of the DE log-fold change 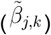 as normalized perturbation effect estimates, where:

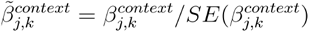

The standard error of the log-fold change is obtained from DESeq2 outputs. We define 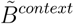 as the *K* x *J* matrix of DE effect estimates in each cellular context.

To improve computational scalability across thousands of perturbed genes, we randomly partitioned perturbed genes into groups of 50 and fit one GLM for each group along with non-targeting controls (NTCs) in each condition in parallel. Pilot experiments with alternative partitioning schemes confirmed that DE estimates for individual perturbed genes were consistent regardless of which genes were included in the model, and that covariate effect estimates remained stable (data not shown). We further verified robustness by comparing mean expression estimates for each condition across partitions.

To assess the statistical calibration of the DE analysis (Suppl. Figure 4B), we performed a null model validation comparing cells with NTC guides. We select amongst guides with cell counts between the 5th and 95th percentile of the NTC distribution to ensure comparable sample sizes to true targeting guides. For each of five independent random splits, we randomly assigned 2 NTC guides to a simulated target group, selecting guides with cell counts between the 5th and 95th percentile of the NTC distribution to ensure comparable coverage to that of the true targeting guides. The remaining NTC guides are used as controls. We performed DE analysis on each split across all three culture conditions (Rest, Stim8hr, Stim48hr) using the same statistical model and design formula applied to targeting perturbations (Suppl. Figure 4B).

We detected significant on-target knock-down in 73% of tested condition-target pairs. Beyond low baseline expression, perturbations without detectable on-target knockdown also showed significantly fewer cells per perturbation (Kolmogorov-Smirnov test, p = 6.57e-12). To estimate off-target prevalence on proximal genes, we annotated each gRNA’s distance to the nearest non-target TSS and assessed DE effects on that gene (Suppl. Table 3). Putative off-targets were flagged where we detected significant knock-down of the nearest TSS gene (DE z-score < −1, FDR < 10%) (Suppl. Table 5).

### Guide robustness analysis

To assess the reproducibility of perturbation effects across independent gRNAs targeting the same gene, we performed guide-level differential expression analysis. For each culture condition, we selected guides meeting the following criteria: (a) minimum of 3 biological replicates per guide; (b) both guides targeting a gene must be testable; (c) strong perturbation effect in the primary DE analysis (>75 perturbed cells and >30 significantly differentially expressed genes at 10% FDR). This filtering yielded 2,040 guides in Rest, 2,350 guides in Stim8hr, and 2,118 guides in Stim48hr conditions. For each guide, we performed differential expression analysis using the same DESeq2-based framework as the gene-level analysis, treating each guide as an independent perturbation.

To quantify reproducibility, we calculated Pearson correlations between log fold changes of guide pairs targeting the same gene. Correlations were computed using only genes showing significant differential expression (10% FDR) in at least one of the two guides, restricted to genes with > 2 total DE genes to ensure sufficient statistical power. Where computed, cross-guide correlation statistics are reported in Suppl. Table 5.

### Validation of *trans*-effects with complementary screens

To validate perturb-seq measurements, we first compared them with transcriptome-wide effects of genetic knock-outs from published arrayed KO screens with bulk RNA-seq measurements [5,16]. DESeq2 statistics (comparing expression levels in perturbed cells against non-targeting controls) were either downloaded from Supplementary tables or obtained from re-analysis of count matrices, replicating the differential analysis design used by the original authors. We compared arrayed screen results for 28 perturbed genes tested in both perturb-seq and arrayed screen datasets, where the perturbation had a strong effect in the arrayed screens (at least 100 significant DE genes at FDR 10%). This analysis was performed using the results from the Rest condition in the perturb-seq screen, to match experimental conditions of the arrayed screens.

We computed the Pearson correlation between DE log-fold changes estimated from perturb-seq and arrayed screen RNA-seq for the measured genes that were significantly DE in the arrayed screen (FDR 10%). As a negative control, for each perturbed gene we also computed the DE effect correlation with perturbation of a randomly picked perturbed gene in the set of 28 tested genes. For each comparison, we performed 100 bootstrap iterations, each randomly sampling 70% of shared measured genes. We estimated the theoretical maximum achievable correlation (correlation ceiling) by accounting for measurement uncertainty in the log fold change estimates of each test. For each test, we estimate the error variance 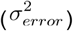 and true variance 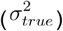 of log-fold changes as:

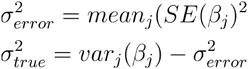

The reliability of log-fold change measurement is calculated as the proportion of observed variance that is due to true signal rather than measurement error:

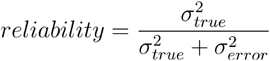

The correlation ceiling was then calculated as the geometric mean of reliabilities for the two guides being compared.

To validate regulator predictions for individual measured genes, we collected a set of published CRISPRi FACS-based screens [6,13,17]. For each FACS-based screen and target gene, we computed correlations between FACS log fold changes (MAGeCK statistics) and perturb-seq z-scores across shared perturbations with FDR < 0.05 in FACS. To control for expression-level confounding, we selected control genes with similar baseline expression (we ranked all genes by mean expression, and picked genes within ±20 expression ranks of the target gene). For each comparison, we performed 100 bootstrap iterations, each randomly sampling 70% of shared perturbations. We then calculated Pearson correlations between effects in FACS-based screen and perturb-seq z-scores for both matched comparisons (target gene in both assays) and mismatched comparisons (FACS target vs. expression-matched controls in perturb-seq).

### Cross-donor robustness analysis

To quantify the reproducibility of perturbation effects across donors, we performed differential expression analysis on donor subsets of the pseudobulk data. This analysis was restricted to condition-target tests meeting the following criteria in the full dataset: >75 perturbed cells and >30 significantly differentially expressed genes at 10% FDR. For each target-condition pair, we conducted 6 separate DE analyses using the method described above, each time subsetting the expression data to include only two donors, representing all possible pairwise combinations of the 4 donors. We then calculated the cross-donor correlation by comparing tests performed on disjoint donor pairs (e.g., donors 1+2 vs. donors 3+4) for the same perturbed gene, using the following procedure:

1. Identify significantly DE genes in each test using a lenient FDR threshold (20%)
2. Take the union of DE genes from both tests being compared
3. Calculate the Pearson correlation of log-fold changes between the two tests for genes in this union
4. Compute the theoretical maximum correlation using the reliability estimate, as described above

For each target-condition pair, there are 3 possible comparisons between disjoint donor pairs. The reported mean and minimum cross-donor correlation represents the average of these 3 correlations (Suppl. Figure 6A-B, Suppl. Table 5).

### Comparison with K562 genome-wide perturb-seq

Gene expression count matrices and cell-level metadata for K562 genome-wide perturb-seq data [9] (K562-GW) were downloaded from the *scPerturb* database [146]. We pseudobulked expression profiles for cells with the same guide in the same batch and performed differential expression analysis on the pseudobulked profiles as described above for our dataset, testing the effect of perturbation compared to non-targeting controls for 3081 perturbed genes that met two criteria: (1) Perturbed genes with significant *trans*-effects on at least one gene other than the target gene in at least one culture condition in CD4+T perturb-seq; (2) perturbed genes with sufficient coverage in K562-GW: at least 3 pseudobulk replicates aggregating data from at least 3 cells, with at least 10 mRNA counts per gene.

To compare DE analysis results accounting for differences in statistical power between the datasets, we processed DE summary statistics for K562 cells and CD4+T conditions using Multivariate Adaptive Shrinkage (MASH [147]), which fits a mixture of multivariate normal distributions to model the joint distribution of effect sizes for all tests (perturbed gene-measured gene) across all 4 cell contexts (K562, CD4+ T Rest, CD4+ T Stim8hr, CD4+ T Stim48hr). The use of MASH posterior estimates is akin to the bivariate analysis proposed by Nadig et al. [106], although we fit a single model for all cell types on a subset of all tests, following the recommended eQTL analysis workflow (https://stephenslab.github.io/mashr/articles/eQTL_outline.html). We then considered 5% local false sign rate (LFSR) as a threshold for significant *trans*-effects across datasets, and consider a *trans*-effect “shared” between cell types if the regulator-to-gene test passed the LFSR threshold in both CD4+T and K562 cells (Suppl. Figure 7A).

We compare effect size estimates between K562 and CD4+T results computing the Pearson correlation between MASH posterior estimates of perturbation effects in K562 and each CD4+T condition (Suppl. Figure 7B). As a negative control, we correlate each the effect of each perturbation to the effect of 3 random perturbations in the other cell types. Results of comparisons of effect sizes are shared in Suppl. Table 6.

### Power analysis

We tested the robustness of perturbation effect estimates to downsampling the number of reads, cells and replicates (donors) profiled per perturbed genes. This analysis was performed in Rest condition and on effects of perturbation of 12 perturbed genes with significant downstream effects replicated by arrayed KO RNA-seq screens (see section *Validation of trans-effects with complementary screens*).

To make downsampled versions of the perturb-seq data, we split 10x lanes into disjoint groups of 5 lanes (with 3 independent splits). For each downsampling experiment, we randomly designated one group of 5 lanes as the validation set, ensuring no overlap with training data (pseudobulking by donor-guide). Then we sequentially pool data from 5, 10, 15 and 20 lanes, simulating an increase in number of cells profiled. Finally, from each group of lanes, we aggregate data from all combinations of 1, 2, 3 or 4 donors, simulating an increase in the number of replicates profiled. To downsample the number of UMIs, we use the method described in [7] on pseudobulk expression profiles. For example, a vector of gene expression counts for 4 genes in a sample with values [3,0,1,6] is converted into the vector [1,1,1, 3,4,4,4,4,4,4] (i.e. from counts to individual instances of each gene), on which sampling is performed with equal probability without replacement. We downsample to 10% of UMIs to match the depth of perturbation datasets profiled with split-pool scRNA-seq protocols [15] and to 50% of UMIs to simulate the depth of perturbation datasets profiled with capture-based 10x Genomics scRNA-seq [9]. We then test for differential expression as described above for the sampled perturbed genes in subsampled pseudobulks with downsampled number of cells, reads or donors.

We evaluate the robustness to downsampling by comparing the DE effects (raw log-Fold changes) for perturbation of each target in each subsample with 2 validation datasets: (a) DE effect estimates in arrayed KO RNA-seq screens from Freimer et al. [5] (Pearson correlation of log-Fold Changes for significant genes at 10% FDR); and (b) DE effect estimates on all donors in the held-out validation set of 5 lanes (Pearson correlation of log-Fold Changes for significant genes at 10% FDR).

### Arrayed validation of IL10/IL21 regulation

At day 0, primary naive human CD4^+^ T cells were seeded at 1 x 10^6^ cells/ml in cXVIVO and 200 IU/ml IL-2 at 150µL/well in 96-well flat bottom plates. Cells were activated using ImmunoCult human CD3/CD28/CD2 T cell activator at 25 µl/ml. The next morning, cells were transduced with 2% v/v concentrated ZIM3KRAB-dCas9-BlastR lentivirus. In the afternoon, cells in each well were transduced with a distinct, individual single gRNA expressing lentivirus (Suppl. Table 2) at 2% v/v. On day 3, cells were split to ~5 x 10^5^ cells/mL and Blasticidin was added at 10 µg/mL final concentration. At day 5, cells were centrifuged and resuspended with cXVIVO supplemented with 200 IU/mL IL-2, 5 ng/mL IL-7 and 10 µg/mL blasticidin at ~5 x 10^5^ cells/mL. At day 7, cells were counted and diluted to 1 x 10^6^ cells/mL with cXVIVO supplemented with 200 IU/mL IL-2, 5 ng/mL IL-7 and 10 µg/mL blasticidin. At day 8, cells were centrifuged and resuspended at 1 x 10^6^ cells/mL with cXVIVO supplemented with 200 IU/mL IL-2 and 5 ng/mL IL-7. On day 10 morning, cells were split into two pools. In one pool, cells were centrifuged and resuspended at 2 x 10^6^ cells/ml with cXVIVO supplemented with 200 IU/mL IL-2 and 1 µL/mL Activation Cocktail with Brefeldin A (BioLegend). After 6 hours, cells were stained for IL-10 (Brilliant Violet 421 anti-human IL-10 Antibody, catalogue number 501422, Biolegend) and IL-21 (PerCP/Cyanine5.5 anti-human IL-21 Antibody, catalogue number 513012, Biolegend) protein expression following the BD Cytofix/Cytoperm kit instructions (BD Biosciences, catalog no. 554714). In another pool, cells were centrifuged and resuspended at 2 x 10^6^ cells/ml with cXVIVO supplemented with 200 IU/mL IL-2 and 12.5 µL/mL ImmunoCult human CD3/CD28/CD2 T cell activator. After 8 hours, cells were centrifuged and resuspended in DNA/RNA Shield (Zymo Research) and sent for bulk RNAseq by Plasmidsaurus.

Count matrices were generated by Plasmidsaurus as follows: quality of the fastq files was assessed using FastQC v0.12.1. Reads were then quality filtered using fastp v0.24.0 with poly-X tail trimming, 3’ quality-based tail trimming, a minimum Phred quality score of 15, and a minimum length requirement of 50 bp. Quality-filtered reads were aligned to the reference genome using STAR aligner v2.7.11 with non-canonical splice junction removal and output of unmapped reads, followed by coordinate sorting using samtools v1.22.1. PCR and optical duplicates were removed using UMI-based deduplication with UMIcollapse v1.1.0. Alignment quality metrics, strand specificity, and read distribution across genomic features were assessed using RSeQC v5.0.4 and Qualimap v2.3, with results aggregated into a comprehensive quality control report using MultiQC v1.32. Gene-level expression quantification was performed using featureCounts (subread package v2.1.1) with strand-specific counting, multi-mapping read fractional assignment, exons and three prime UTR as the feature identifiers, and grouped by gene_id. Raw fastq and count matrices are available via GEO (see “Data Availability“).

From count matrices, we performed differential expression analysis using DESeq2 [143] as previously described. We fit a negative binomial generalized linear model where the expected mRNA count was modeled as a function of donor identity and sample perturbation target gene (design: `~donor+target_gene)`.

The non-targeting control samples of this experiment were intentionally designed to be transduced with a mixture of two non-targeting control gRNAs (NTC425 and NTC720, Suppl. Table 2). This is to avoid systematic perturbation effect biases from small chance of using a non-targeting control gRNA that induces off-target effects.

### Functional clustering of perturbations

#### Clustering workflow

To define regulator clusters, we performed clustering on the 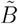 matrix of differential expression z-scores, representing the effects of perturbations on measured genes, where each perturbation corresponds to a perturbed gene in a cellular context (culture condition). We first filtered for high-quality perturbations (each perturbation is a perturbed gene x culture condition) with strong effects, requiring >75 differentially expressed genes (DEGs) and >50 cells per perturbation. This yielded 3341 perturbations across three conditions (Rest: 1088; Stim8hr: 1136; Stim48hr: 1117). We also trimmed the feature space to 10941 highly variable genes (from an initial 13959) by requiring a sum of absolute z-scores >= 1,200 (~20 percentile) across the selected perturbations. To preclude strong on-target knockdown effects from driving cluster assignment, on-target z-scores were masked to zero.

We used Leiden clustering on the first principal components of the filtered 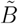 matrix, using the methods implemented in scanpy (scanpy.tl.leiden). We observed that the resulting cluster assignments showed some sensitivity to hyperparameter selection (PCA dimensions, k-neighbors, and leiden resolution). To ensure stability and robustness against hyperparameter selection and sampling bias, we implemented a consensus clustering approach. We generated a grid ensemble of 27 clustering parameters, varying PCA dimensions (50, 100, 150), k-nearest neighbors (k=15, 31, 63), and Leiden clustering resolution (r=2, 3, 4) across 400 randomly bootstrapped subsampled datasets. Specifically, for each of the 27 hyperparameter combinations, we then performed clustering with the hyperparameter grid on 200 perturbation-subsampled and 200 measured gene-subsampled datasets (70% random sampling rate), resulting in 10800 clustering outcomes. After collecting all the clustering results, we computed a pairwise co-occurrence matrix representing the frequency with which perturbations were co-clustered across all runs and converted this to a distance matrix (1 - consensus frequency). Final clusters were identified using HDBSCAN on this precomputed distance matrix (using the implementation in scikit-learn, min_cluster_size=4, min_samples=1, cluster_selection_method=‘eom’), resulting in 111 regulator clusters.

#### Annotation of clusters

We annotated the possible functions of clusters by performing gene set enrichment analysis on clustered regulators on public biological complexes, pathways, and process datasets including CORUM, STRINGDB, KEGG, and Reactome. To focus on more specific complexes/pathways, we only used terms in these databases with less than 200 genes. For each database, we then performed a hypergeometric test to detect complex/pathway enrichment among unique regulator genes in each cluster compared to all regulators included in the clustering analysis (genes with >75 total DE genes and >50 cells per target). In cases where no overlap existed between the tested gene sets and complex/pathway genes, we assigned a p-value of 1. We controlled for multiple testing using the Benjamini-Hochberg method to estimate false discovery rates. Based on FDR ranking, we reported the best-matching protein complex name (CORUM), the best functional description (STRINGDB) or the top associated pathways (KEGG and Reactome), enrichment significance (hypergeometric test p-value/FDR), the fraction of cluster genes overlapping with the term, and the specific overlapping genes. We assigned annotations to clusters showing strong enrichment (FDR < 0.05). For clusters without strong enrichment in any of these datasets, we used a combination of Gene Ontology analysis (https://geneontology.org/), LLM lookup, and manual literature search.

#### Assessing condition specificity of clusters

To assess the condition specificity of regulator clusters (Figure 3A-B), we analyzed the correlation of perturbation effects between distinct regulators. For a given cluster containing a set of regulator genes *G*, we computed the mean pairwise correlation for each condition *c* ∈ {Rest,Stim8hr,Stim48hr}:

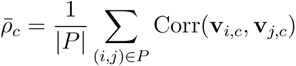

where ***v***_*i,c*_is the perturbation effect vector (z-score) for gene *i* in condition *c*, and *P* = {(*i,j*) *i,j* | *i,j* ∈ *G,i* < *j*} represents all unique pairs of distinct regulators. We similarly computed a “Shared” mean correlation, 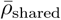, using pairs of perturbations drawn from different conditions.

We classified cluster specificity (e.g. used in Figure 7A, Suppl. Figure 13) using the coefficient of variation (*CV*) of the intra-condition means 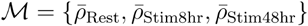. Clusters with *CV* < 0.5 were classified as “across_condition.” Clusters with high variability (*CV* ≥0.5) were further sub-classified based on condition enrichment. A condition *c* was considered enriched if it satisfied two criteria:

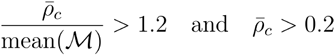

Clusters were labeled according to their enriched conditions (e.g., “Rest”, “Rest_Stim8hr”). High-variability clusters meeting no enrichment criteria were retained as “across_condition”.

Cluster annotations and condition-specificity statistics are provided in Suppl. Table 9.

#### Curation, ranking and annotation of downstream genes

To characterize the downstream target genes associated with each regulator cluster, we performed a rank aggregation analysis on the perturbation effects. Because we frequently observe the same set of regulators genes in a cluster regulate distinct downstream genes in different conditions, the curation and ranking of downstream genes are stratified by conditions. For each cluster identified by HDBSCAN, we first get the differential gene expression results of regulators on all measured genes in one of the three conditions. Then we defined the set of potential downstream genes as the union of all genes significantly differentially expressed (10% FDR) by at least one regulator within that cluster in the corresponding condition.

For each potential downstream gene, we calculated three key metrics to quantify the strength and consistency of the *trans*-effect:

1. Upstream regulator count: the number of regulators that significantly affected the gene (10% FDR) in the corresponding condition.
2. Sign Coherence: to assess the consistency of directionality, we calculated the sign coherence as the sum of the signs (+/-1) of the z-scores for all significant regulators, normalized by the upstream regulator count. A score of 1 or −1 indicates perfect directional agreement among regulators.
3. Aggregate Rank Score: to prioritize genes with the strongest aggregate response, we utilized a rank-sum approach. For each downstream gene, we first converted the perturbation z-scores into ranks within each individual regulator perturbation in the corresponding condition. We then summed these ranks across all regulator perturbations in that corresponding condition.This yielded a single cumulative rank metric for each downstream gene, where the lowest cumulative sums correspond to the strongest negative z-score regulation (i.e. regulator perturbations strongly downregulate the downstream gene) and the highest cumulative sums correspond to the strongest z-score regulation (i.e. regulator perturbations strongly upregulate the downstream gene).

These downstream genes of regulator clusters are reported in Suppl. Table 10

To annotate biological processes of top downstream genes of selected clusters (Figure 3C, Figure 3D, Suppl. Figure 15), we performed Gene Ontology enrichment analysis using the gseapy Python package (https://github.com/zqfang/GSEApy), using the GO Biological Process 2025 libraries. For each cluster, enrichment was performed independently for its downstream genes in one of the three conditions and stratified by regulation direction (positive or negative). To prioritize the strongest co-regulated downstream genes, we selected downstream genes based on their Aggregate Rank Score in the positive or negative direction. Because the total numbers of potential downstream genes (significantly differentially expressed by at least one regulator for that cluster) for each cluster are different, the input gene list for enrichment was defined as the top 100 genes or the top 3% genes, whichever is larger. The background gene set for statistical testing consisted of the complete pool of downstream genes identified within the specific cluster and condition being analyzed. Significant GO terms were identified using an FDR threshold of <0.05. To generate a concise and non-redundant list of biological terms, we removed terms that have more than 10% overlap with any more significant term.

#### Tissue-specific gene expression analysis

To characterize the tissue-specific expression patterns of genes in regulator clusters (Suppl. Figure 13), we utilized bulk RNA-seq data from the Human Protein Atlas (HPA) version 25.0 (based on Ensembl version 109) [148,149]. We obtained the consensus transcript expression levels dataset (https://www.proteinatlas.org/about/download, “RNA expression consensus”), which summarizes gene expression across 51 human tissues. These consensus normalized expression (nTPM) values represent the maximum nTPM value reported for each gene between the HPA and Genotype-Tissue Expression (GTEx) datasets. For organ systems containing multiple sub-tissues (e.g., brain regions, lymphoid tissues, and intestine), the maximum nTPM value across all sub-tissues was used to represent the tissue type. For each gene, to quantify the tissue specificity of gene expression, we performed two analyses.

For overall tissue specificity of gene expression, we calculated the Tau (τ) index as described by Yanai et al., [150]. The Tau index provides a scalar value ranging from 0 (broadly expressed) to 1 (highly tissue-specific). It is calculated as:

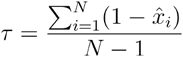

Where *N* represents the number of tissues (51), and 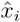 is the expression value of the gene in tissue *i*, normalized by the maximal expression value across all tissues:

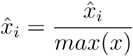

Prior to calculation, expression values were log1p-transformed. Genes exhibiting a Tau index greater than 0.5 were classified as having high tissue specificity.

To determine specific tissue enrichment for each gene, we calculated the log2 fold change of gene’s expression (nTPM) in a specific tissue relative to its median expression across all 51 tissues. A gene was considered enriched in a specific tissue if the log2 fold change > 2.5.

### Cell state signature analysis

#### Model overview

To infer the regulators of a cell state of interest in observational data, we used a regularized regression approach that integrates differential gene expression signatures with context-specific perturbation effect estimates (Figure 4A). The model takes as input (a) differential expression estimates of genes in the cell state (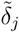 for gene *j*, representing the z-score of the log-Fold change in differential expression), compared to a reference cell state in the observational dataset; (b) Perturbation effect estimates for *K* candidate regulators across genes in the perturb-seq data (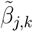, z-score of the log-Fold change representing the effect of regulator *k* on gene *j*). We train each model with 5-fold cross validation, holding-out from the full set of measured genes a test set to use for model evaluation.

To reduce the dimensionality of the perturbation effect matrix and mitigate multicollinearity among regulators, we applied truncated singular value decomposition (SVD) to the perturbation matrix 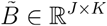. The truncated SVD approximates the original matrix as:

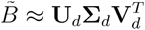

where ***U***_*d*_∈ ℝ ^*j* × *D*^ contains the first *D* left singular vectors, **∑**_*d*_∈ ℝ ^*D* × *D*^ is a diagonal matrix of the *D* largest singular values, and ***v***_*d*_∈ ℝ ^*K* × *D*^ contains the first *D* right singular vectors. The matrix 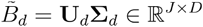 represents the effect of *D* eigen-perturbations on *J* genes. This decomposition was performed using the TruncatedSVD implementation in scikit-learn [151]. Unless otherwise specified, we used *D* = 60 for applications in this study, after selection with a small grid search on reconstruction of Th2/Th1 polarization signatures.

We fit an elastic net regression model to predict the differential expression signature 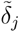 from the reduced-dimensional perturbation effects 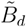. The elastic net model solves the following optimization problem:

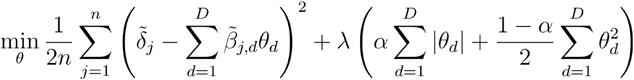

where *θ* = (*θ*_1_, …, *θ*_*D*_) are the regression coefficients for the *D* eigen-perturbations, *λ* is the overall regularization strength, and *α* is the ratio that balances L1 and L2 penalties. Hyperparameters *λ* and *α* were optimized using 4-fold cross-validation on the training set. The model was implemented using scikit-learn’s ElasticNet class [151].

To evaluate model performance on held-out genes, we first projected the test set perturbation effects onto the eigen-perturbation space learned from the training set:

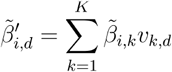

where 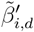 *s the effect of eigen-perturbation d on test gene* 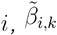 is the original perturbation effect of regulator *k* on gene *i, v*_*k,d*_is the (*k,d*) element of *V*_*D*_, and *K* is the total number of regulators. We then applied the fitted elastic net model to obtain the reconstructed 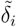 from 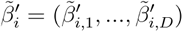.

To obtain interpretable coefficients at the level of individual regulators (*w*_*k*_), we back-transformed the eigen-perturbation coefficients (*θ*_*d*_) through the right singular vector matrix:

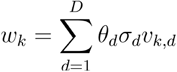

where *σ*_*d*_is the *d*-th singular value and *v*_*k,d*_is the (*k,d*) element of **V**_*D*_. These regulator-level coefficients represent the estimated contribution of each regulator’s perturbation signature to the observed cell state differential expression signature. We used mean coefficients across training folds for downstream analyses (Figure 4F, Figure 5D). The model is implemented as a stand-alone python class (https://github.com/emdann/pert2state_model).

For visualization of regulator-level coefficients (Figure 4F, Figure 5D), we implemented a modified version of the “Long Island City plots” introduced by Grabski et al. [111], where in each plot, the x-axis consists of the set of K regulators included in the model fit, ordered by hierarchical clustering of the DE effects of regulators on significantly DE genes in the cell state of interest.

We consider signature genes as “controlled by a set of *m* regulators” if they meet the following conditions: (a) significant DE from knock-down of least one of the regulators (10% FDR) and (b) the sign of mean DE z-score across all regulators in the set needs to be consistent with the sign of predicted regulator effect (*w*_*k*_) and of the cell state signature (*δ*_*i*_):

- If *w*_*k*_> 0: a gene upregulated by the regulators (*mean* (*z*_*j*_) > 0) should be upregulated in the target cell state (*δ*_*i*_> 0)
- If *w*_*k*_< 0: a gene upregulated by the regulators (*mean* (*z*_*j*_) > 0) should be downregulated in the target cell state (*δ*_*i*_> 0)

#### Th1/Th2 polarization bulk RNA-seq data processing and analysis

Bulk RNA-seq counts for Th1 and Th2 sorted cells were downloaded from the NBDC Human Database for the Discovery cohort (Accession: E-GEAD-397)[39] and from GEO for the Replication cohort (Accession: GSE149090)[40].

For each cohort, differential expression analysis was performed using DESeq2 [143] as previously described. We fit a negative binomial generalized linear model where the expected mRNA count was modeled as a function of T cell subset (Th1 or Th2), the first principal components capturing technical variation and donor identity (where replication was present). The model designs were: `~ cell_subset + PC1 + PC2`for the discovery cohort and `~ cell_subset + donor + PC1`for the replication cohort. We estimated log2 fold-changes in gene expression and tested for differential expression between Th2 and Th1 samples using the Wald test. FDR < 1% after Benjamini-Hochberg correction was used as threshold for significance. Full DESeq2 results are provided in Suppl. Table 11. The z-score of the differential expression log-fold change was used as the input signature for the model 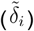. As a negative control, we used a TCR activation signature derived from DESeq2 results comparing stimulated and resting Teff cells from [13] (Suppl. Figure 16F).

To identify regulators of Th2 and Th1 polarization from perturb-seq effects, we considered regulators with significant differential expression effects on at least 10 measured genes in any condition. We fitted the differential expression signature reconstruction model (as described above) to reconstruct the effect of Th2/Th1 polarization, using 5-fold cross-validation on 8947 measured genes with 3 independent splits, evaluating model fit on a validation set of held-out genes in each of the 15 splits. The knock-down effect of each perturbed gene on itself was masked before training. To evaluate model fit, we computed the Pearson correlation between the reconstructed Th2/Th1 signature and the true input signature for both training set and test set genes in the Discovery cohort (samples used for training) and Replication cohort (samples not used during training). For comparisons with scrambled controls (Figure 4E, Suppl. Figure 16D), we scrambled the differential expression z-scores for each gene across regulators, preserving the mean z-score for each measured gene. This approach accounts for the tendency of highly expressed genes to show larger z-score effects. For comparison with perturbation effects in K562 cells (Figure 4, Suppl. Figure 16D), we used differential expression estimates obtained as described above (see section *Comparison with Replogle et al*) and fitted the regression model using only K = 2190 common perturbed genes with sufficient power in both the Stim8hr CD4+ T cell and K562 datasets. Regulator-level regression coefficients (*w*_*k*_) for models fitted on CD4+ T cell conditions to reconstruct Th2/Th1 and activation signatures are provided in Suppl. Table 12.

To estimate expression of Th2 and Th1 signature genes in single cells from perturb-seq data (Figure 4G, Suppl. Figure 17), we used a modified version of the *sc*.*tl*.*score_genes* function implemented in scanpy [141] to account for donor-specific differences in expression profiles. Briefly, we obtained normalized single-cell expression profiles for cells with guides targeting the regulators of interest and a subset of non-targeting control guides. We defined signature genes as those differentially expressed at 1% FDR in the Discovery dataset, selecting the bottom 50 genes (Th1 signature) and top 50 genes (Th2 signature) ranked by z-score of differential expression log-fold change. For each gene set (Th1 and Th2 signatures), we retained genes expressed in the dataset (mean log-normalized expression > 0.1) and standardized expression values using the mean and standard deviation calculated from NTC cells matched by donor and condition. We computed the mean standardized expression across all genes in each signature as the final score.

#### Aging scRNA-seq data processing and analysis

Single-cell RNA-seq data from the OneK1K cohort were downloaded from the CellxGene collections (collection ID: dde06e0f-ab3b-46be-96a2-a8082383c4a1) [60]. Quality control filtering was applied to remove low-quality cells based on the following criteria: cells with >8% mitochondrial reads, or cells with values ±3 median absolute deviations (MADs) from the median percentage of mitochondrial reads, or ±5 MADs from the median for total counts, number of detected genes, or percentage of counts in the top 20 genes. We subsequently subset the dataset to retain only cells annotated by the original authors as CD4+ T cells, including the subset labels “CD4 Naive”, “CD4 EM”, and “CD4 CM” (Figure 5A).

For differential expression analysis, we retained genes that were measured across all experimental batches (pools) and selected the top 10,000 highly variable genes using the scanpy function *sc*.*pp*.*highly_variable_genes* (parameters: *min_mean=0*.*1, max_mean=10, n_top_genes=10000*). Gene expression data were aggregated to the pseudo-bulk level by summing mRNA counts from all cells belonging to the same donor and cell subset annotation. We partitioned the dataset into discovery (n = 782 donors) and replication (n = 199 donors) cohorts, by splitting donors while stratifying by age bins (<40, 40-60, 60-70, 70-80, >80 years). Principal component analysis of pseudo-bulked gene expression profiles revealed that the first principal components captured technical variation between experimental batches (each donor was processed in a single batch) and donor outliers in cell type composition. Consequently, we included the first 5 principal components computed on the “CD4 Naive” subset as covariates in the differential expression model to account for these confounding factors. For each cohort, differential expression analysis was performed using DESeq2 [143] as previously described. We fit a negative binomial generalized linear model where the expected mRNA count was modeled as a function of CD4+ T cell subset (Naive, EM, CM), the first 5 principal components capturing technical variation, the number of aggregated cells, donor sex, and age bin. We estimated log2 fold-changes in gene expression and tested for differential expression across age using the Wald test, with the age bin modelled as a continuous covariate. This model design was chosen to identify genes that exhibit age-associated expression changes across all CD4+ T cell subsets, while controlling for expression differences between subsets that might reflect age-related shifts in the composition of the T cell compartment (Suppl. Figure 19A). FDR < 1% after Benjamini-Hochberg correction was used as threshold for significance. Full DESeq2 results are provided in Suppl. Table 13. We evaluated the robustness of the resulting differential expression estimates by comparing results between the discovery and replication cohorts, and by benchmarking against published estimates from independent cohorts that tested for age-associated differential expression in CD4+ T cells [64,65] (Suppl. Figure 19B). Outlier genes with inconsistent differential expression effects across cohorts were predominantly red blood cell genes, including those related to hemoglobin metabolism (e.g. HBB), likely reflecting contamination from ambient RNA during single-cell RNA-seq processing. The scaled z-score of the differential expression log-fold change with age was used as the input signature for the model. We applied centering to account for a slight bias in DE estimates, leading to logFCs being skewed towards positive values (mean logFC = 0.02), likely due to imbalance in sample numbers between age bins.

To identify regulators of the CD4+T aging signature from perturbation effects, we considered regulators with significant differential expression effects on at least 10 measured genes in the condition of interest. We fitted the differential expression signature reconstruction model (as described above) to reconstruct the signature of CD4+T cell aging, using 5-fold cross-validation on 6014 measured genes. The knock-down effect of each perturbed gene on itself was masked before training. To evaluate model fit, we computed the Pearson correlation between the reconstructed aging signature and the true input signature for both training set and test set genes in the Discovery cohort (samples used for training) and Replication cohort (samples not used during training). Regulator-level regression coefficients (*w*_*k*_) for models fitted on CD4+ T cell conditions to reconstruct aging signatures are provided in Suppl. Table 14.

### Integration with human population loss-of-function (LoF) burden data

We used LoF burden test summary statistics from the UKB with 454,787 participants, as previously reported [3]. Gene effect sizes were de-noised using GeneBayes [152] and previously described [3]. Replicating the analysis in [3], to identify putative core genes that directly influence lymphocyte counts, we ran, for each core gene candidate *j*, a linear regression to test the association between knockdown effects of *K* regulators on the gene (log-Fold Change, *β*_*j,k*_) and the LoF burden on lymphocyte counts (*γ*_*k*_). To avoid the confounding effects of selective constraint due to the large correlation between LoF burden test effect sizes and gene-level effects on fitness (*S*_het_) [153], we fit the following regression model:

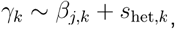

where *S*_het_ is included as a covariate. We masked the effects of knock-down gene on itself, as it does not reflect a trans-regulatory effect. We obtained regulator-burden correlations using perturbation effects estimated from all three CD4+ T cell culture conditions in our genome-wide perturb-seq, as well as cell lines K562 [9] and Jurkat [106] (using DE estimates from [3]). For Stim8hr and Stim48hr conditions, we evaluated the putative core genes positively associated with lymphocyte counts using Gene Ontology enrichment analysis. We obtained genes with top 50 regulator-burden correlations by − log_10_*P*, and ran gene set enrichment analysis using the *gseapy* Python package (https://github.com/zqfang/GSEApy), using all tested genes in each condition as background.

### Analysis of effects of autoimmunity GWAS genes

We queried the OpenTargets Platform API (v4) to retrieve genes with genetic association with autoimmune diseases (rheumatoid arthritis, systemic lupus erythematosus, inflammatory bowel disease, multiple sclerosis, type 1 diabetes, psoriasis, ankylosing spondylitis, asthma, Hashimoto’s thyroiditis, celiac disease, and atopic eczema) and non-autoimmune control diseases (coronary artery disease, macular degeneration, and chronic kidney disease). We included genes with a minimum genetic evidence score of 0.1 (https://platform-docs.opentargets.org/evidence) derived from GWAS associations, gene burden studies, and somatic mutation data, which includes evidence from ClinVar (Suppl. Figure 23A).

To test for enrichment of disease-associated genes among the regulators in each cluster and/or their downstream targets, we excluded clusters containing fewer than five unique regulators. We then performed Fisher’s exact tests to assess enrichment of disease-associated genes in two gene sets for each condition: (a) cluster regulators and (b) condition-specific downstream target genes. For regulator enrichment, we tested whether disease-associated genes were overrepresented among the regulators in each cluster compared to all regulators included in the clustering analysis (genes with >75 total DE genes and >50 cells per target). For downstream gene enrichment, we first selected the top 50 genes with the lowest z-score rank for positive regulation and the top 50 genes with the lowest z-score rank for negative regulation within each cluster and condition (see section “Curation, ranking and annotation of downstream genes”). We then tested whether disease-associated genes were overrepresented among the downstream targets of each cluster compared to all selected downstream genes across all clusters in that condition. Downstream genes were only considered in conditions where regulators exhibited coordinated effects. In cases where no overlap existed between the tested gene sets and disease-associated genes, we assigned a p-value of 1. We controlled for multiple testing using the Benjamini-Hochberg method to estimate false discovery rates. The complete results and tested gene sets are provided in Suppl. Table 15.

## Data availability

Cell-level count matrices, pseudobulk-level count matrices and differential expression estimates are available at https://virtualcellmodels.cziscience.com/dataset/genome-scale-tcell-perturb-seq. Raw sequencing data and cellranger outputs will be made available through SRA/GEO (accession: SRP643211 / GSE314342). Supplementary tables and additional metadata are available via our code repository (https://github.com/emdann/GWT_perturbseq_analysis_2025).

## Code availability

All processing and analysis code is available at https://github.com/emdann/GWT_perturbseq_analysis_2025. The model to predict regulators of observed cell states is implemented as a stand-alone python class (https://github.com/emdann/pert2state_model).

## Notes

https://virtualcellmodels.cziscience.com/dataset/genome-scale-tcell-perturb-seq

https://github.com/emdann/GWT_perturbseq_analysis_2025

https://github.com/emdann/pert2state_model

