## Supplementary materials for "Genome-scale perturb-seq in primary human CD4+ T cells maps context-specific regulators of T cell programs and human immune traits"

#### Supplementary notes

##### Supplementary Note 1. Donor-to-donor robustness of DE estimates

We investigated the robustness of perturbation effect estimates across donors. For perturbations (condition-perturbed gene) showing strong effects in the test in all donors (defined as >75 perturbed cells and >30 significantly differentially expressed genes at 10% FDR), we performed differential expression (DE) analysis on all possible donor pairs (yielding 6 tests per condition-target). To quantify reproducibility, we then compared tests from disjoint donor pairs by calculating the Pearson correlation between DE log-fold changes across the union of DE genes identified in each pair (termed "cross-donor correlation"). On average, perturbation effects showed positive correlation across donors (Suppl. Figure 6A, mean cross-donor correlation = 0.38, n = 4,621 target-condition pairs). Cross-donor correlations for the same target were generally consistent across conditions, though some targets exhibited high consistency in one condition but not others (Suppl. Figure 6B). We interpret these cases as perturbations whose effects manifest in a condition-specific manner. We further examined cases with inconsistent perturbation effects across donors. Some instances of low cross-donor correlation could be attributed to noise in the DE estimates (Suppl. Figure 6C). Additionally, we found that inconsistent effects typically arose when one donor pair showed substantially stronger overall signal than another, whereas cases where disjoint donor sets yielded entirely different sets of DE genes were rare (Suppl. Figure 6D). Notably, we observed several instances where DE tests yielded significant effects only upon inclusion of a specific donor. While some cases likely represent cell culture artifacts, others suggest that donor-to-donor variability may cause perturbation effects to manifest in a donor-specific manner. A notable example involves known Th17 regulators (SMAD3 [154,155], ICOS [156], RASGRP1 [157], IL12RB2 [158]): knockdowns of these genes showed significant downstream effects only when including data from donor 2 (Suppl. Figure 6E). However, the transcriptome-wide perturbation effects were significantly correlated across perturbations, and the affected downstream genes were consistent with established roles in Th17 differentiation (e.g., significant upregulation of *RORC*). These findings suggest that donor-specific differences in baseline cell states can cause known regulatory effects to manifest exclusively in certain donors, highlighting that increasing donor diversity may enhance discovery potential.

#### Supplementary Note 2. Polarization prediction with perturbation effects in Rest condition

We compared models trained to predict Th1/Th2 polarization signatures using perturbation effects measured under different culture conditions. The model trained on effects in resting cells showed significantly better performance than the model trained on effects in stimulated cells (Suppl. Figure 18A). However, these two models prioritized distinct sets of regulators. While perturbations of certain regulators were equally predictive across both models (e.g., GATA3, RARA, IRF1), the resting cell model prioritized several regulators of T cell activation and effector function (e.g., LAT2, MED24, CD69) (Suppl. Figure 18B).

We hypothesize that this difference reflects the incorporation of activation state variation into the Th1/Th2 signature itself. Across cohorts, multiple activation-associated genes, including granzyme genes, LYAR, CCL4, and GBP5, were significantly overexpressed in Th1 cells compared to Th2 cells (Suppl. Figure 16F). In stimulated conditions, all cells are already activated at baseline, which may diminish the effect of perturbations on activation-related genes and reduce their predictive value for the Th1/Th2 signature. Conversely, the effects of certain master polarization regulators may manifest only in stimulated conditions where relevant cytokine signaling is active. For instance, IL4R knockdown effects may require the presence of IL4-secreting cells to be detected.

#### Supplementary Note 3. Direction of mTORC1 effects on age-related CD4+ T cell changes

In our analysis to predict regulators of aging signatures in CD4+ T cells, we identified mTORC1 pathway components as key regulators. However, the predicted direction of effect on the aging signature was opposite to expectations based on existing literature: mTORC1 inhibitors were predicted as positive regulators of the aging signature (i.e., knock-down of TSC and GATOR complex genes upregulates genes that are downregulated in aged CD4+ T cells), while RPTOR was predicted as a negative aging regulator. This finding warrants further investigation given that hyperactivation of mTOR is considered a hallmark of aging: treatment with the mTOR inhibitor rapamycin can extend longevity in mammals [66], and pharmacological mTOR inhibition has shown potential to improve immune function in the elderly [67,68]. In T cells, mTOR serves as a signaling node to activate several downstream effector pathways, including immune receptor signaling, metabolic programs, and migratory activity [69]. Aged T cells exhibit increased basal activation of PI3K–AKT–mTOR signaling [63].

To validate our perturbation response estimates, we examined the effects of inhibitors and activators on gold-standard mTORC1 activation signature genes, which showed consistent patterns (Suppl. Figure 21A). Additionally, transcriptome-wide comparison of perturbation responses across different mTOR pathway components revealed expected correlation patterns, with repressors and activators of mTORC1 showing significantly correlated and anti-correlated responses, respectively (Suppl. Figure 21B). These results provide confidence in the robustness of our perturbation response estimates.

When examining expression signatures of CD4+ T cells in aged individuals in the OneK1K cohort, we detected upregulation of both positive regulators of mTORC1 (e.g., RAGulator components LAMTOR4, LAMTOR5, RRAGA, activating V-ATPases responding to amino acids, FLCN; Suppl. Figure 19D) and negative regulators (STRADA components, CAB39, DDIT4), as well as genes inhibited by mTORC1 signaling (e.g., ULK2, ULK3). Although aged T cells in this dataset do not exhibit strong evidence for mTORC1 hyperactivation, their transcriptional signature is significantly explained by perturbation effects of key mTORC1 regulators. This pattern suggests that the observed expression profile in T cells from older individuals may reflect a late or adaptive response to prior mTORC1 activation rather than ongoing pathway hyperactivity. Supporting this interpretation, we observed upregulation of multiple known mTORC1 inhibitors in aged CD4+T cells, including PIM kinases, which directly inhibit the TSC complex [159,160], and *SESN3*, an established mTORC1 inhibitor [161].

We propose three non-mutually exclusive hypotheses to explain this pattern (Suppl. Figure 21):

1. **Negative feedback via PI3K-Akt:** Negative feedback loops are well-characterized in mTORC1 signaling. mTORC1 activation leads to phosphorylation of ribosomal protein S6 kinase (S6K), which functions in ribosomal regulation but also acts as an inhibitor of PI3K/Akt. Reduced PI3K/Akt signaling subsequently inhibits mTORC1 via the TSC complex, establishing a self-regulating negative feedback loop. Studies have demonstrated that mTOR inhibition can paradoxically increase mTOR signaling by disrupting the PI3K/Akt feedback loop [162–164], while knockout of mTOR inhibitors conversely leads to PI3K activation in mice [165]. This feedback mechanism is considered one reason for rapamycin's disappointing clinical efficacy, prompting second-generation mTOR therapies that target multiple pathway nodes to circumvent feedback activation [69,166].

2. **Other compensatory pathways:** Hyperactivation of mTORC1 may trigger compensatory upregulation of additional pathway inhibitors beyond the canonical feedback mechanisms.

3. **Survival bias:** T cells with sustained high mTORC1 activity may undergo cell death, enriching the surviving population for cells that maintained lower mTORC1 activation levels despite aging-associated pressures. This interpretation is particularly relevant given the age of this cohort, with the oldest age bin covering individuals between 80 and 100 years old. Individuals reaching these advanced ages may possess more resilient cells, representing a cohort with advantageous cellular phenotypes.

### Supplementary figures

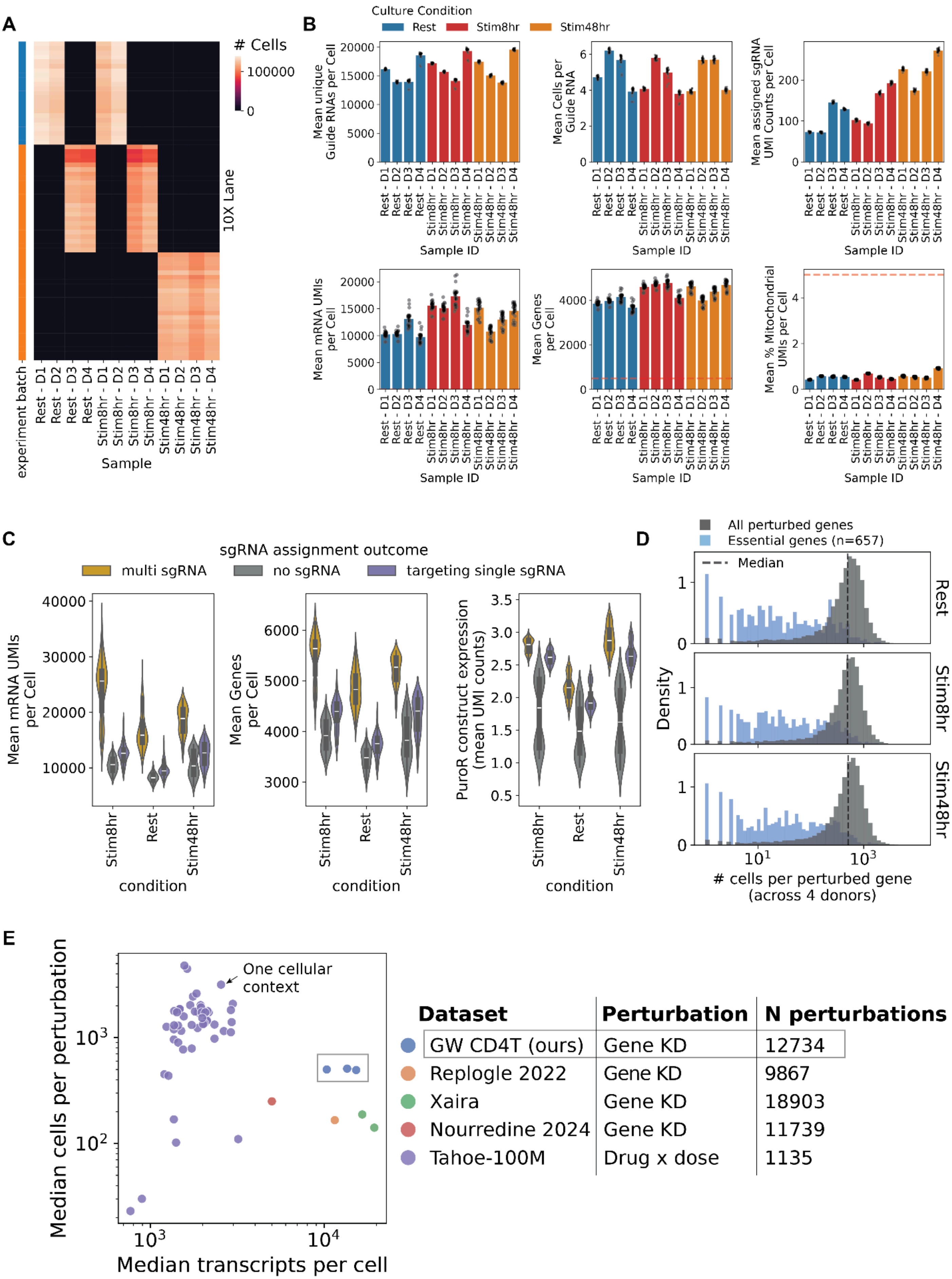
A
### Cells
100000
0
10X Lane
experiment batch
Rest - D1
Rest - D2
Rest - D3
Rest - D4
Stim8hr - D1
Stim8hr - D2
Stim8hr - D3
Stim8hr - D4
Stim48hr - D1
Stim48hr - D2
Stim48hr - D3
Stim48hr - D4
Sample
B
Culture Condition
Rest
Stim8hr
Stim48hr
Mean unique Guide RNAs per Cell
Mean Cells per Guide RNA
Mean assigned sgRNA UMI Counts per Cell
Mean mRNA UMIs per Cell
Mean Genes per Cell
Mean % Mitochondrial UMIs per Cell
Sample ID
C
sgRNA assignment outcome
multi sgRNA
no sgRNA
targeting single sgRNA
Mean mRNA UMIs per Cell
Mean Genes per Cell
PuroR construct expression (mean UMI counts)
Stim8hr
Rest
Stim48hr
condition
D
All perturbed genes
Essential genes (n=657)
Median
Density
Rest
Stim8hr
Stim48hr
### cells per perturbed gene (across 4 donors)
E
One cellular context
Median cells per perturbation
Median transcripts per cell
Dataset | Perturbation | N perturbations
GW CD4T (ours) | Gene KD | 12734
Replogle 2022 | Gene KD | 9867
Xaira | Gene KD | 18903
Nourredine 2024 | Gene KD | 11739
Tahoe-100M | Drug x dose | 1135

***Supplementary Figure 1*. *Study design and quality control (QC) metrics.*** *(**A**) Design of cell culture and scRNA-seq experimental batches: for each biological sample (x-axis) we show the number of cells recovered from each 10x lane. The row annotation indicates which lanes were processed in each experiment batch. Samples profiled in the same lane were multiplexed. (**B**) Summary of QC metrics (y-axis) for each biological sample (x-axis). Each point represents a 10x lane, barplots show the mean across lanes. Barplots show from left to right: mean number of scRNA-seq UMI counts per cell, mean number of genes captured per cell, mean percentage of mitochondrial reads per cell (lower is better), and mean number of UMIs from assigned guide RNA in each cell (excluding UMI counts for guides below the signal-to-noise threshold). Dotted lines indicate threshold values that were used to filter cells with low quality transcriptomes during preprocessing. (**C**) Violinplots comparing quality control statistics across cells with different guide assignments (color) in each condition (x-axis). PuroR: puromycin resistance transgene (**D**) Cell coverage per perturbed gene across four donors, shown separately for each experimental condition, comparing the number of cells recovered per gene for all perturbed genes (grey) and perturbed essential genes (blue, n=657, reference: [https://github.com/macarthur-lab/gene_lists/blob/master/lists/CEGv2_subset_universe.tsv](https://github.com/macarthur-lab/gene_lists/blob/master/lists/CEGv2_subset_universe.tsv)). Dashed lines indicate median coverage in all genes. (**E**) Comparison of summary statistics between this dataset (GW CD4T) and other released large scale perturb-seq datasets (color). Statistics were collected from supplementary tables in the original studies [9,10,15,99]. Each point represents a cell line/context. x-axis: median number of transcripts per cell; y-axis: median number of cells per perturbation. The type and total number of perturbations is annotated in the legend.*

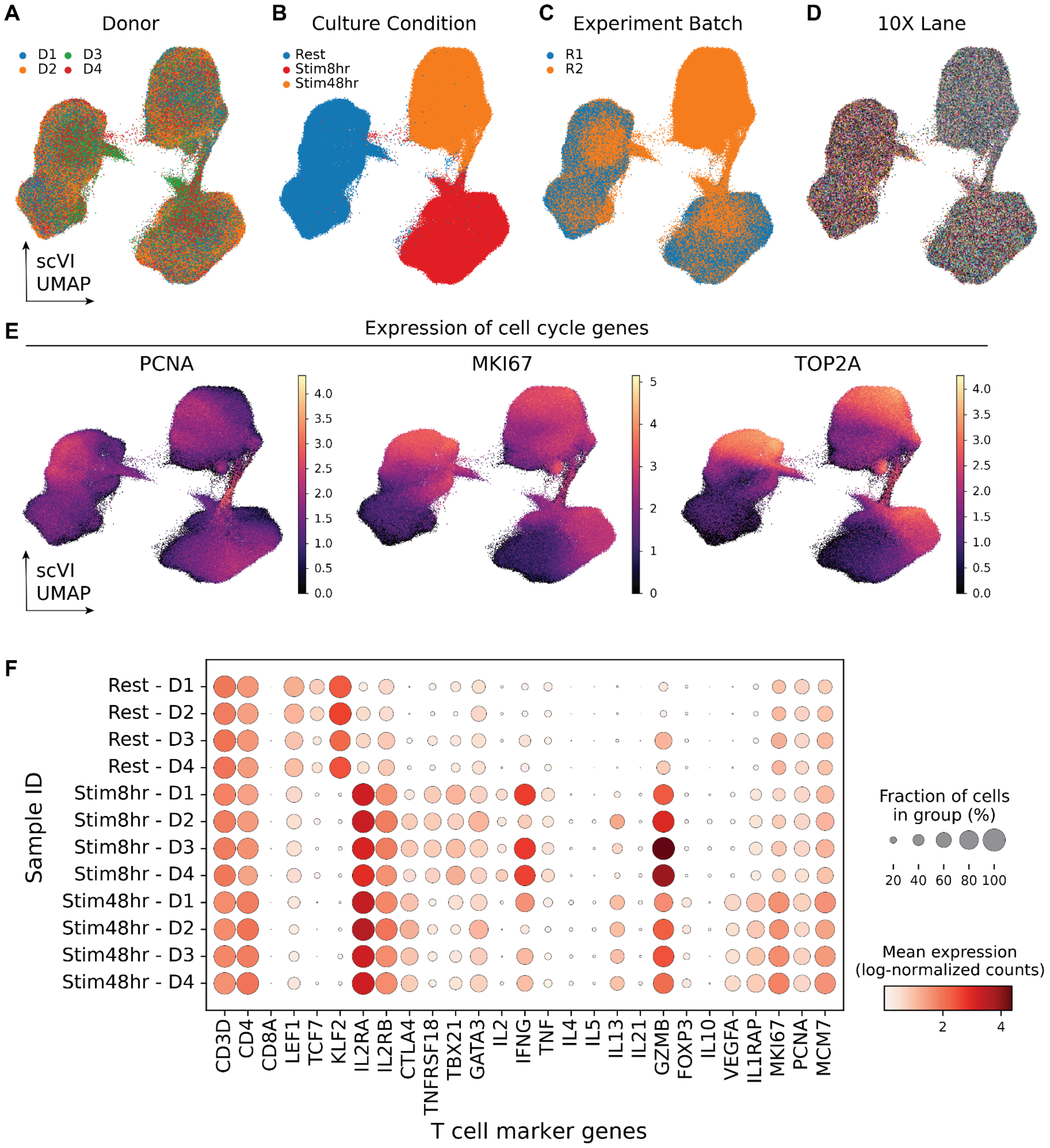


***Supplementary Figure 2**. **Cell state variation in non-targeting control (NTC) CD4+ T cells.** (**A-E**) UMAP visualization of scVI embeddings of a subset of NTC cells (n=395030 cells) in the CD4+ T perturb-seq dataset (see Methods). Points are colored by the donor (A), culture condition (B), experiment batch (C), sequencing lane (D), and log-normalized expression of cell cycle marker genes (E). (**F**) Dotplot showing mean expression (log-normalized counts, dot color) and fraction of NTC cells expressing (dot size) canonical CD4+ T cell markers (x-axis) in different biological samples (y-axis).*

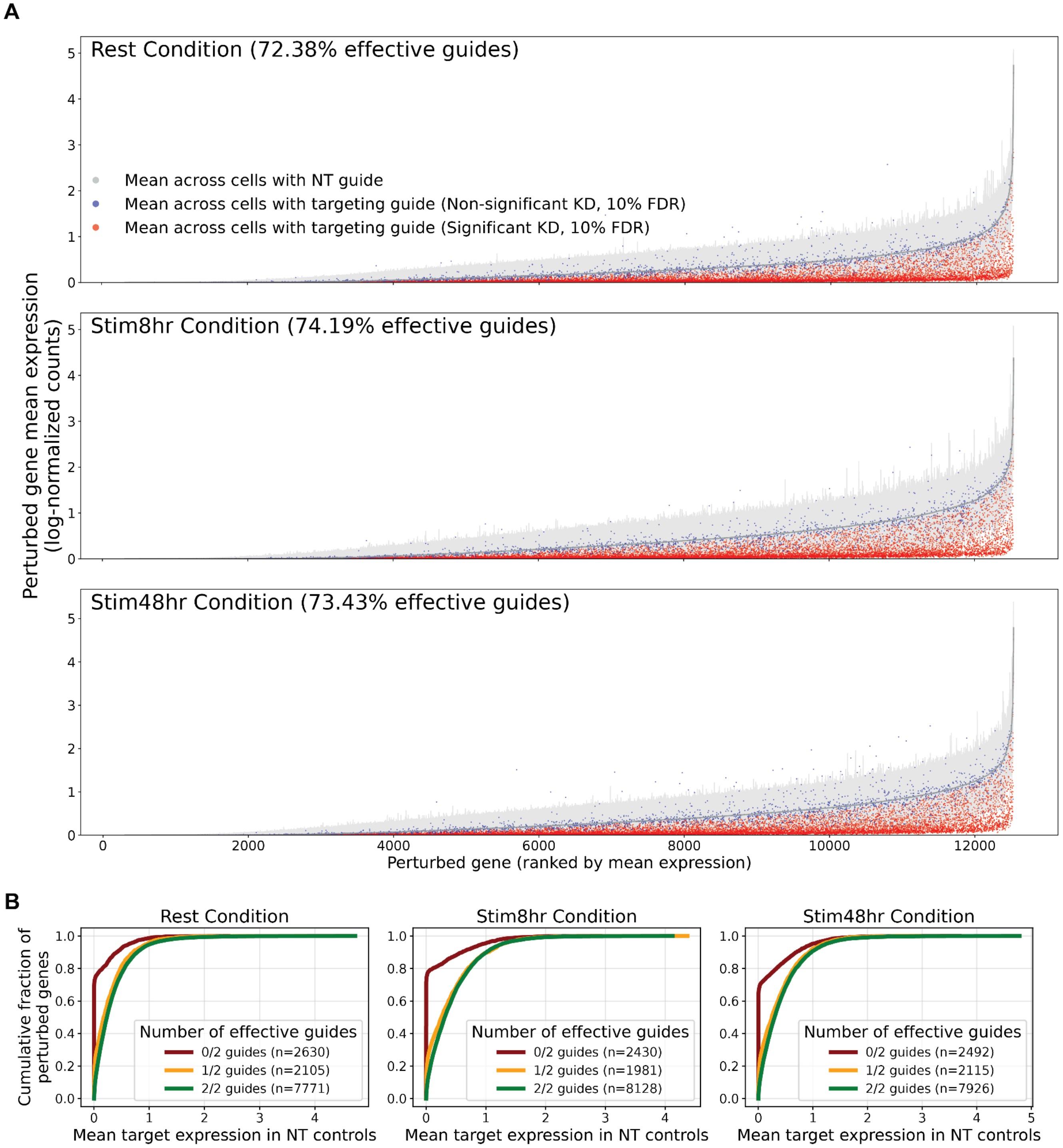


***Supplementary Figure 3. Knockdown efficiency estimates.*** *(**A**) Perturbed gene mean expression (log-normalized counts, y-axis) ranked by expression level of the perturbed gene (x-axis) in non-targeting (NT) control cells (gray) and cells with targeting guide (red = significant knockdown, blue = non-significant knockdown, see methods). Each point is a guide. Results for each condition are shown. (**B**) Cumulative distribution of perturbed gene expression in NT controls, stratified by the number of effective guides for the perturbed gene (0/2 guides in brown, 1/2 guides in yellow, 2/2 guides in green) across conditions.*

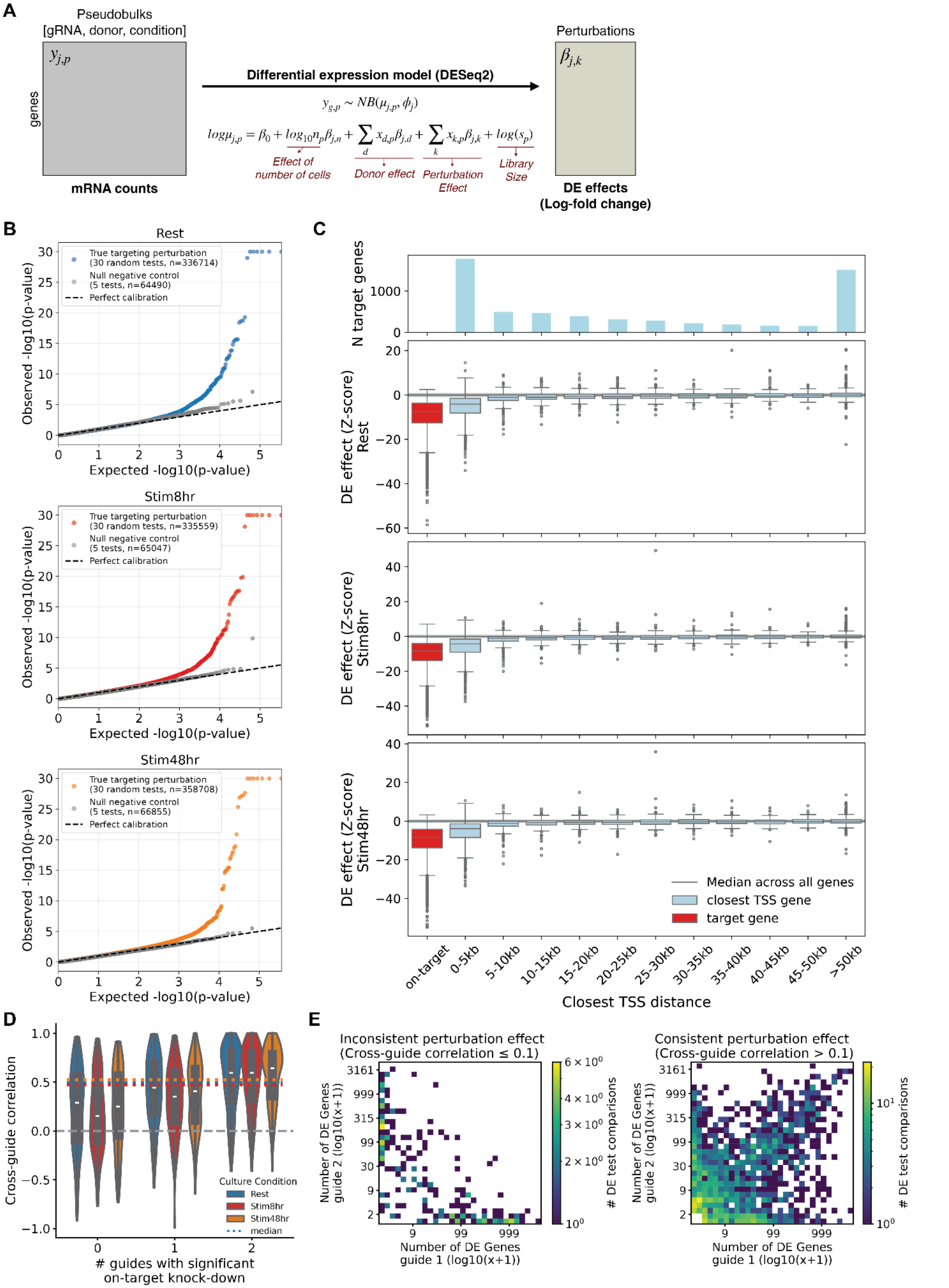


$$y_{g,p} \sim NB(\mu_{j,p}, \phi_j)$$

$$log\mu_{j,p} = \beta_0 + log_{10}n_p\beta_{j,n} + \sum_d x_{d,p}\beta_{j,d} + \sum_k x_{k,p}\beta_{j,k} + log(s_p)$$

***Supplementary Figure 4*. Quality control of perturbation effects quantified by differential expression analysis.** *(**A**) Schematic of differential expression analysis design. (**B**) Quantile-quantile plots comparing observed versus expected -log10(p-values) for null and true differential expression tests in Rest, Stim8hr, and Stim48hr conditions. Blue points represent true targeting perturbations (n = 336,714 to 358,708 tests), gray points represent null negative control tests (n = 64,490 to 66,055 tests), and the dashed line indicates perfect calibration (expected uniform distribution under the null hypothesis). (**C**) Differential expression effects (z-score, y-axis) of perturbations on target gene (in red) compared to gene with the closest transcription start site (light blue), within increasing distance bins (x-axis). DE effect estimates for each condition are shown in different rows. The top barplot shows the number of target genes with the closest gene within each bin. The line and shaded area denote the median and interquartile range of DE effect across all genes. (**D**) Violin plots showing the Pearson correlation of differential expression effects between guide pairs targeting the same gene, stratified by the number of guides (0, 1, or 2) achieving significant on-target knockdown. Correlations are shown separately for Rest (n = 952 perturbed genes), Stim8hr (n = 1123 perturbed genes), and Stim48hr (n = 977 perturbed genes) conditions. Dashed horizontal lines indicate median correlations per condition. (**E**) 2D histogram comparing the number of differentially expressed (DE) genes detected by two independent guides targeting the same gene. Left: guide pairs with inconsistent effects (cross-guide correlation ≤ 0.1). Right: guide pairs with consistent effects (cross-guide correlation > 0.1). Color scale indicates the number of DE test comparisons in each bin.*

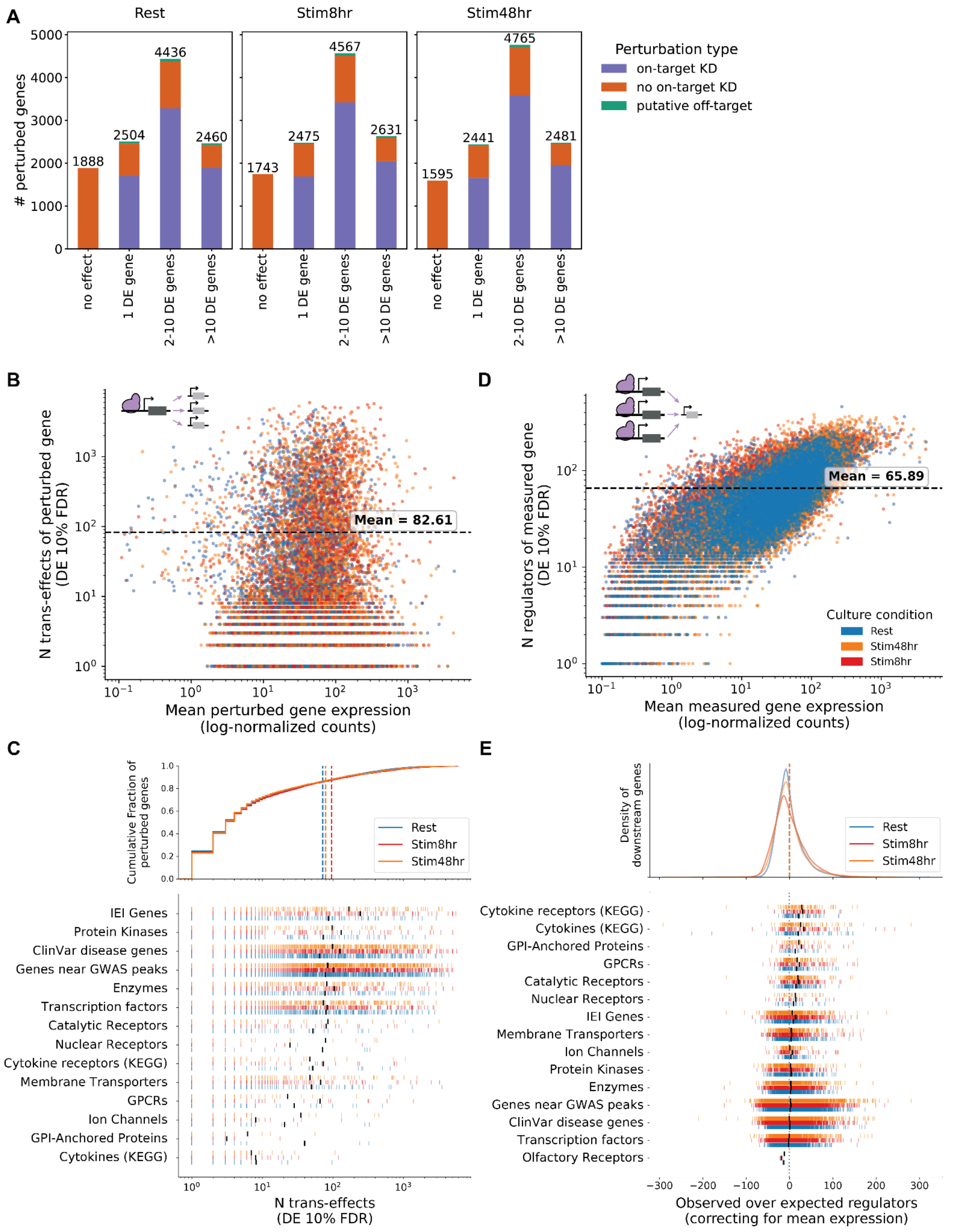
A
Rest
Stim8hr
Stim48hr
### perturbed genes
Perturbation type
on-target KD
no on-target KD
putative off-target
no effect
1 DE gene
2-10 DE genes
>10 DE genes
1888
2504
4436
2460
1743
2475
4567
2631
1595
2441
4765
2481
B
N trans-effects of perturbed gene (DE 10% FDR)
Mean = 82.61
Mean perturbed gene expression (log-normalized counts)
D
N regulators of measured gene (DE 10% FDR)
Mean = 65.89
Culture condition
Rest
Stim48hr
Stim8hr
Mean measured gene expression (log-normalized counts)
C
Cumulative Fraction of perturbed genes
Rest
Stim8hr
Stim48hr
IEI Genes
Protein Kinases
ClinVar disease genes
Genes near GWAS peaks
Enzymes
Transcription factors
Catalytic Receptors
Nuclear Receptors
Cytokine receptors (KEGG)
Membrane Transporters
GPCRs
Ion Channels
GPI-Anchored Proteins
Cytokines (KEGG)
N trans-effects (DE 10% FDR)
E
Density of downstream genes
Rest
Stim8hr
Stim48hr
Cytokine receptors (KEGG)
Cytokines (KEGG)
GPI-Anchored Proteins
GPCRs
Catalytic Receptors
Nuclear Receptors
IEI Genes
Membrane Transporters
Ion Channels
Protein Kinases
Enzymes
Genes near GWAS peaks
ClinVar disease genes
Transcription factors
Olfactory Receptors
Observed over expected regulators (correcting for mean expression)

***Supplementary Figure 5*. Trans-effects of genetic perturbations.** *(**A**) Summary of number of significant DE genes (10% FDR) for each perturbed gene. Perturbed genes are grouped according to the number of significant downstream DE effects. The colors indicate whether we detected significant on-target downregulation for a perturbed gene (purple), no on-target KD (orange), or no on-target KD but significant KD in a gene within 10 kb of the target (green). Summaries for each culture condition are shown. (**B**) Scatterplot of mean perturbed gene expression (log-normalized counts, x-axis) against the number of trans-regulatory effects per perturbed gene (y-axis). Points are colored by culture condition. The dashed horizontal line marks the mean number of trans-effects across all perturbed genes. (**C**) Distribution of trans-regulatory effects across functional categories of perturbed genes. The upper panel shows the cumulative fraction of genes as a function of the number of trans-effects (DE at 10% FDR) across three culture conditions. Vertical dashed lines indicate mean values for each condition. The lower panel displays the number of trans-effects for individual genes within functionally defined categories, with each vertical tick representing a single gene colored by culture condition. The black line denotes the mean for each category in each condition. (**D**) Scatterplot of mean measured gene expression (log-normalized counts, x-axis) against the number of incoming trans-regulatory effects per measured gene (y-axis). Points are colored by culture condition. The dashed horizontal line marks the mean number of incoming trans-effects across all measured genes. (**E**) Distribution of incoming trans effects across functional categories of measured genes. The upper panel shows the distribution of observed over expected regulators per measured gene across three culture conditions. The expected number of regulators is determined based on the mean expression level of each measured gene, by fitting a Poisson generalized linear model, and taking the residuals to estimate the observed vs expected number of regulators. The lower panel displays the number of trans-effects per measured gene within functionally defined categories, with each vertical tick representing a single gene colored by culture condition. The black line denotes the mean for each category in each condition. (IEI = Inborn Errors of Immunity)*

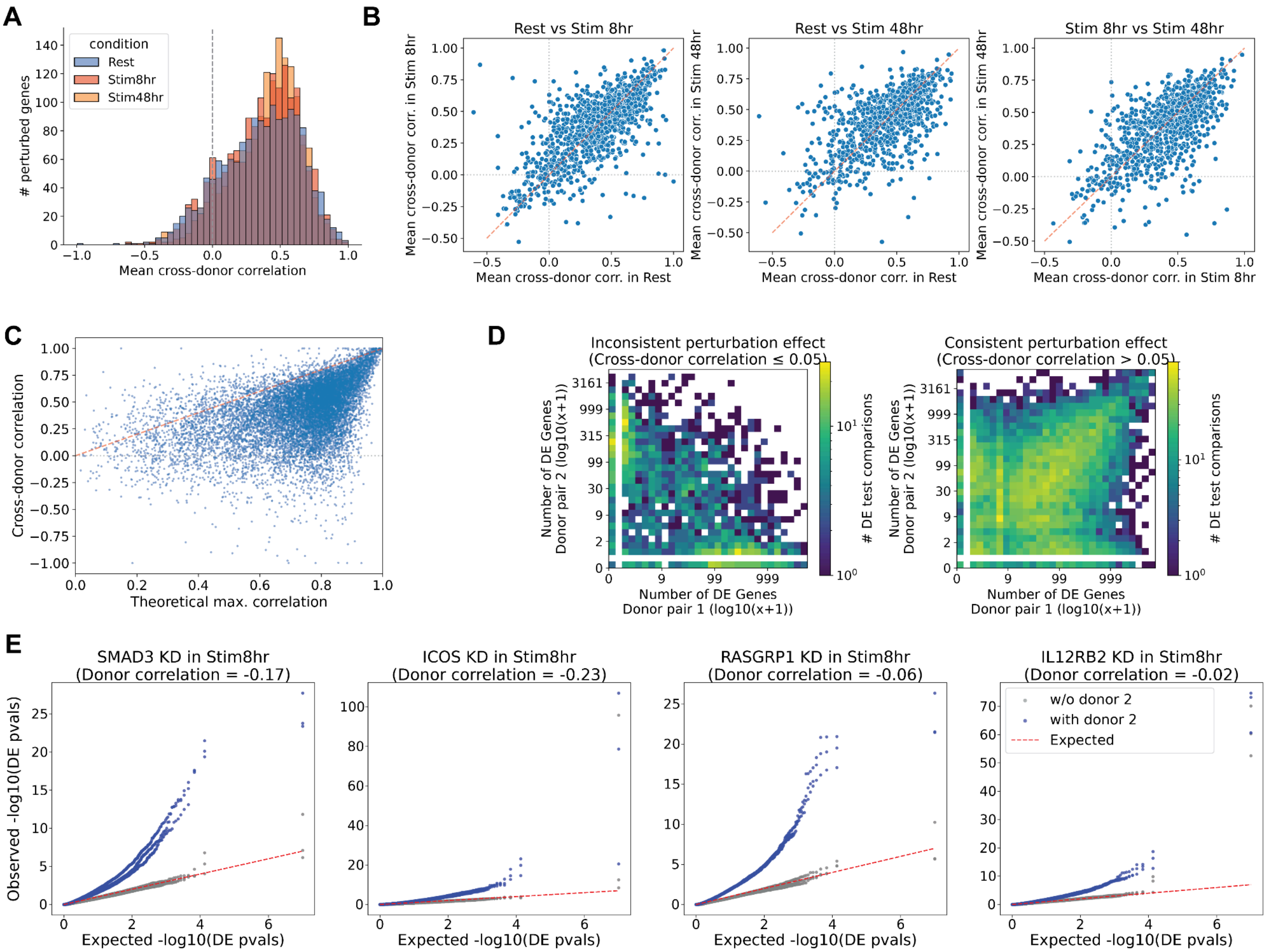


***Supplementary Figure 6**. **Robustness of perturbation effects across donors.** (**A**) Histograms of mean cross-donor correlation for 4621 target-condition pairs. For each target-condition, the mean is calculated across 3 comparisons between non-overlapping pairs of donors. Distributions for comparisons in each culture condition are shown in different colors. (**B**) Comparisons of mean cross-donor correlations for the same targets in different conditions. (**C**) Scatterplot showing the theoretical maximum correlation given noise in DE estimates (x-axis, see Methods) against the measured cross-donor correlation between non-overlapping pairs of donors (13863 comparisons on 4621 target-condition pairs). (**D**) 2D histograms comparing the number of significant DE genes (10% FDR) in the DE test on the first donor pair (x-axis) and second donor pair (y-axis). Results are shown for target-condition pairs with inconsistent effects (mean cross-donor correlation ≤ 0.05, left plot) and consistent effects (mean cross-donor correlation > 0.05, right plot). (**E**) Q-Q plots of DE test p-values across genes for tests over different pairs of donors for perturbation of 4 known Th17 regulators (SMAD3, ICOS, RASGRP1, IL12RB2). Dots in blue denote tests including data from donor 2.*

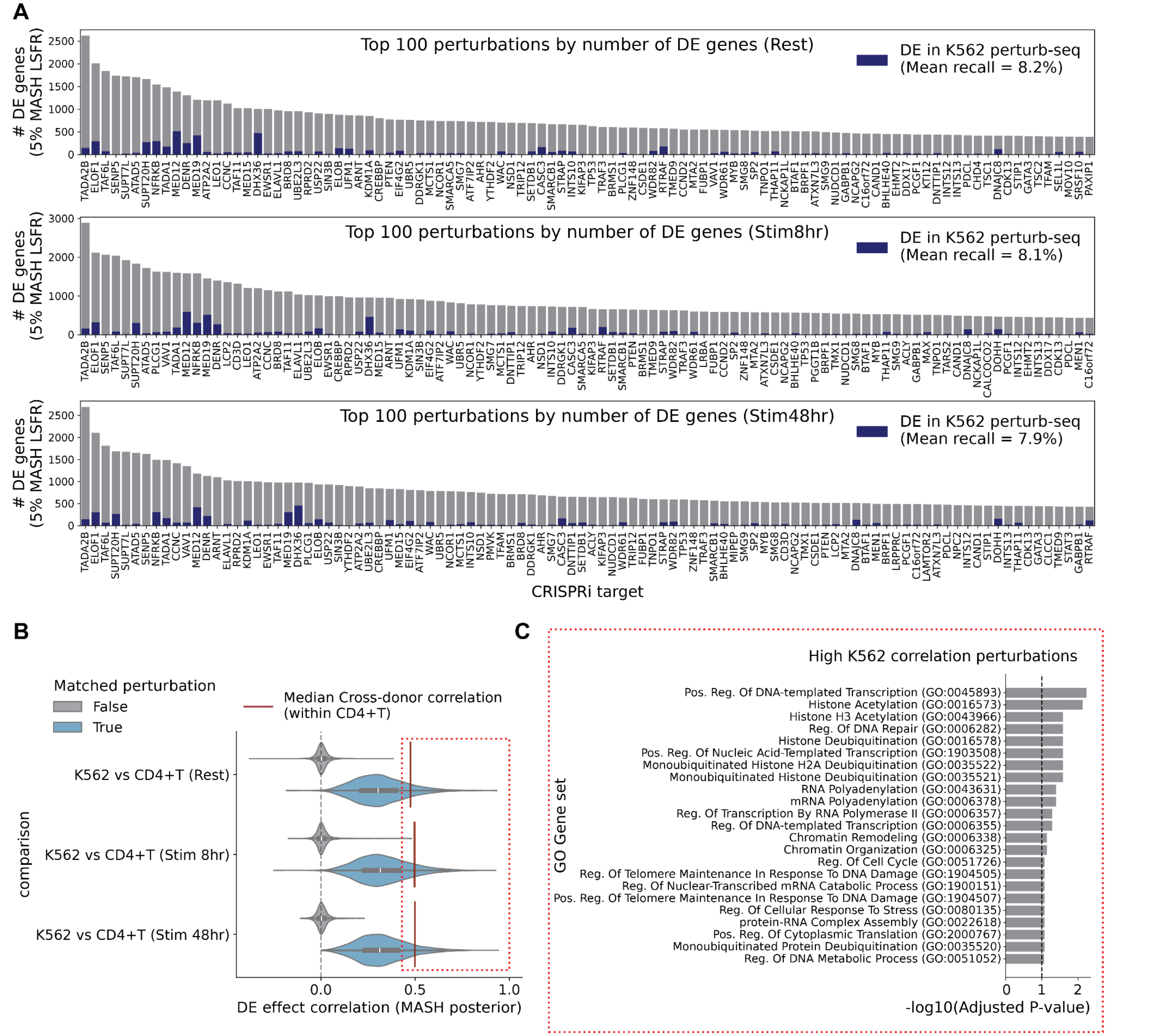


***Supplementary Figure 7. Comparison with trans effects in K562 genome-wide screen.*** *(**A**) Barplots of number of trans effects in CD4+T screen for top 100 perturbations by number of DE genes for each condition. Blue bars indicate trans-effects also identified in K562. DE results were jointly modeled using Multivariate Adaptive Shrinkage (MASH) to account for power differences (see Methods). Mean recall in K562 is reported for 2670 perturbed genes with sufficient coverage across datasets. (**B**) Distribution of correlations between DE effects (MASH posterior) measured in K562 vs CD4+T perturb-seq. Light blue: same perturbed gene across datasets; gray: random gene pairs. Correlations are shown for 1880 perturbed genes with significant trans-effects in at least 3 genes in the K562 dataset. DE effects on the target genes were masked for this analysis. The dark red line denotes the median cross-donor correlation in the CD4+T perturb-seq data (see* Suppl. Figure 6*) (**C**) Gene Ontology biological process terms significantly enriched among perturbed genes with highly correlated perturbation effects between K562 and CD4+T cells (top 20 percentiles of correlation in B). Gene set enrichment analysis was performed using the EnrichR method [167] implemented in the gseapy python package. Horizontal bars represent the negative log10 of adjusted p-values for each GO term; dashed line indicates FDR < 10%.*

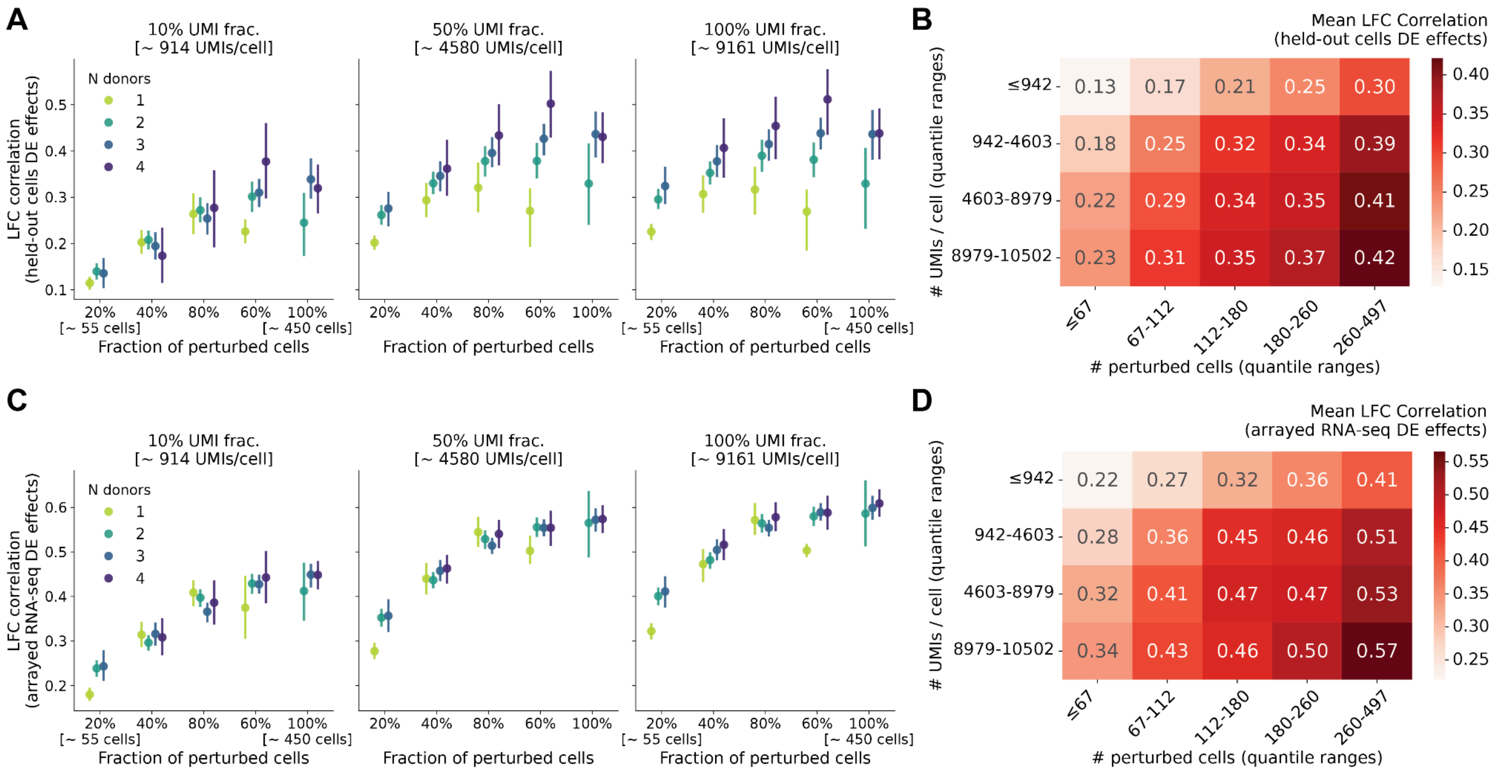


***Supplementary Figure 8*****.** ***Power analysis for perturb-seq dataset****. (****A****) Pearson correlation between log-fold changes (LFC) estimated from downsampled data and held-out validation data (5 lanes, all donors), computed for genes significant at 10% FDR in the held-out validation data. Each panel shows results at different sequencing depths: 10%, 50%, and 100% of UMIs (corresponding to approximately 914, 4,580, and 9,161 UMIs per cell, respectively). The x-axis indicates the fraction of perturbed cells included (20–100%, corresponding to approximately 55–450 cells). Colors indicate the number of donors (1–4). Error bars represent standard error across independent downsampling splits. Approximate cell counts at 20% and 100% fractions are indicated below the x-axis. The mean results across DE tests on 12 perturbed genes are shown (see* Suppl. Figure 9 *for individual perturbed genes). (****B****) Heatmap summarizing mean LFC correlation with held-out validation data across quantile ranges of perturbed cell counts (x-axis) and UMIs per cell (y-axis). Values indicate mean Pearson correlation coefficients, computed for genes significant at 10% FDR in the held-out validation data. (****C****) As in (A), showing Pearson correlation between LFC estimates from downsampled data and independent arrayed RNA-seq knockout screen data (Freimer et al. 2022, see Methods), computed for genes significant at 10% FDR in the arrayed screen data. (****D****) As in (B), showing mean LFC correlation with arrayed RNA-seq knockout screen data, computed for genes significant at 10% FDR in the arrayed screen data.*

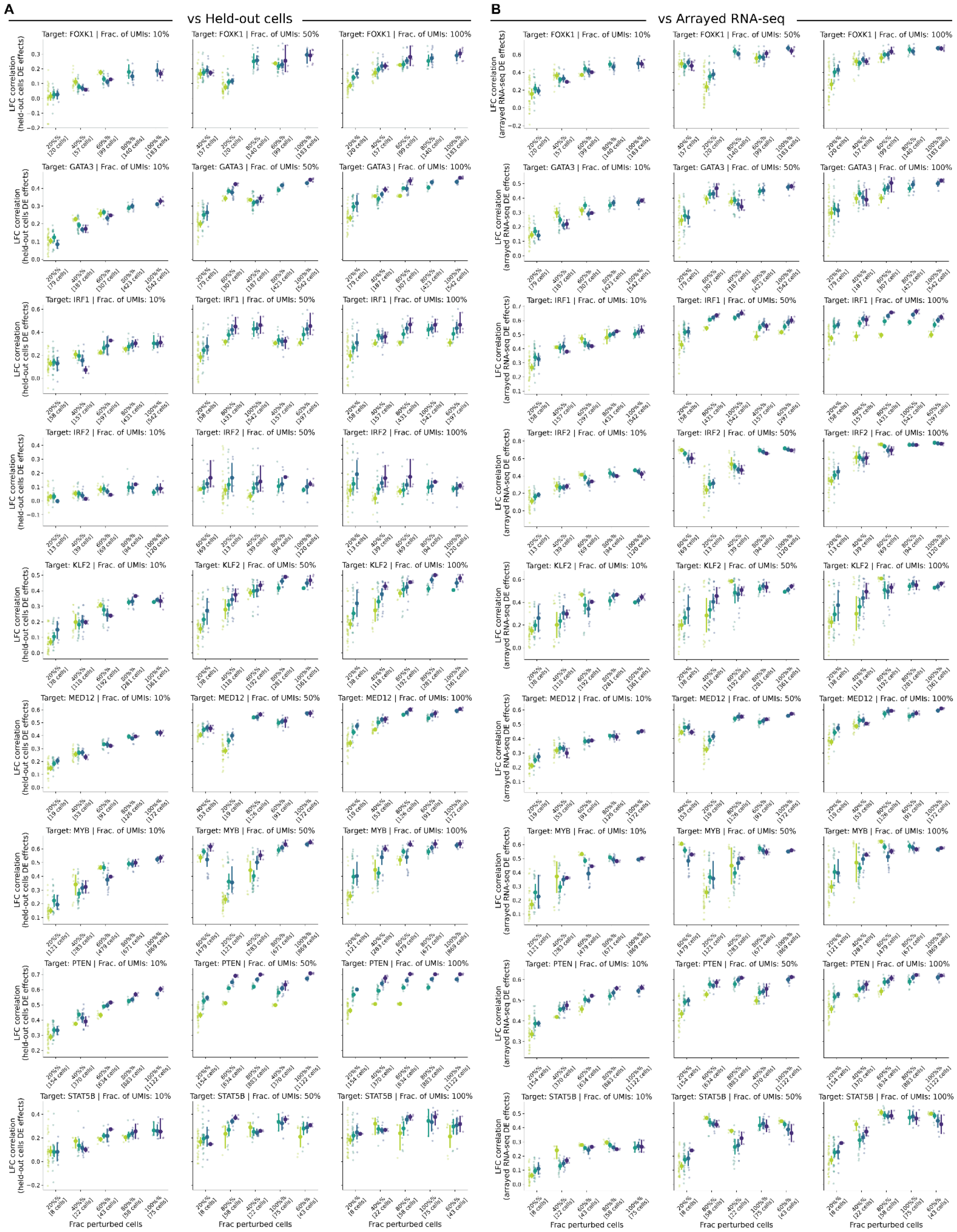


***Supplementary Figure 9. Power analysis results for each perturbed gene*** *(see* Suppl. Figure 8*)*

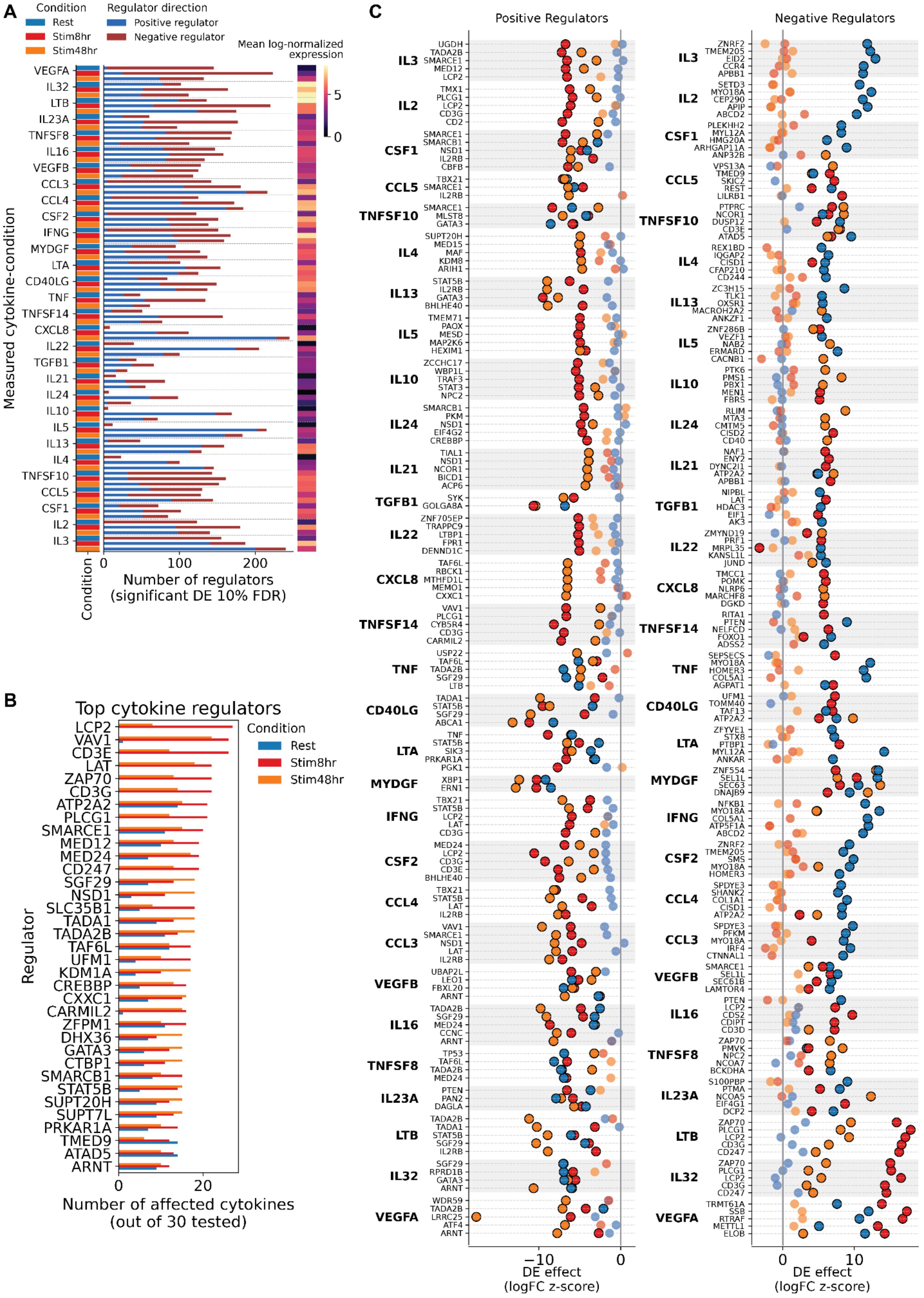
A
Condition
Rest
Stim8hr
Stim48hr
Regulator direction
Positive regulator
Negative regulator
Mean log-normalized expression
Measured cytokine-condition
VEGFA
IL32
LTB
IL23A
TNFSF8
IL16
VEGFB
CCL3
CCL4
CSF2
IFNG
MYDGF
LTA
CD40LG
TNF
TNFSF14
CXCL8
IL22
TGFB1
IL21
IL24
IL10
IL5
IL13
IL4
TNFSF10
CCL5
CSF1
IL2
IL3
Condition
Number of regulators (significant DE 10% FDR)
B
Top cytokine regulators
Regulator
LCP2
VAV1
CD3E
LAT
ZAP70
CD3G
ATP2A2
PLCG1
SMARCE1
MED12
MED24
CD247
SGF29
NSD1
SLC35B1
TADA1
TADA2B
TAF6L
UFM1
KDM1A
CREBBP
CXXC1
CARMIL2
ZFPM1
DHX36
GATA3
CTBP1
SMARCB1
STAT5B
SUPT20H
SUPT7L
PRKAR1A
TMED9
ATAD5
ARNT
Condition
Rest
Stim8hr
Stim48hr
Number of affected cytokines (out of 30 tested)
C
Positive Regulators
Negative Regulators
DE effect (logFC z-score)
DE effect (logFC z-score)

***Supplementary Figure 10. Overview of trans-effects on cytokines.*** *(**A**) Barplot showing the number of positive and negative regulators (x-axis, at 10% FDR) for each of 30 cytokines (y-axis) in each culture condition (left color bar). The mean expression of each cytokine in each condition is shown in the colorbar to the right (log-normalized expression). (**B**) Barplot of number of affected cytokines for top cytokine regulators, for each culture condition (color). Regulators are sorted by maximum number of affected cytokines. (**C**) Scatterplot showing the DE effect (logFC z-score, x-axis) of top 5 positive (left) and negative regulators (right) for each of the measured cytokines (cytokines ranked by z-score across conditions).*

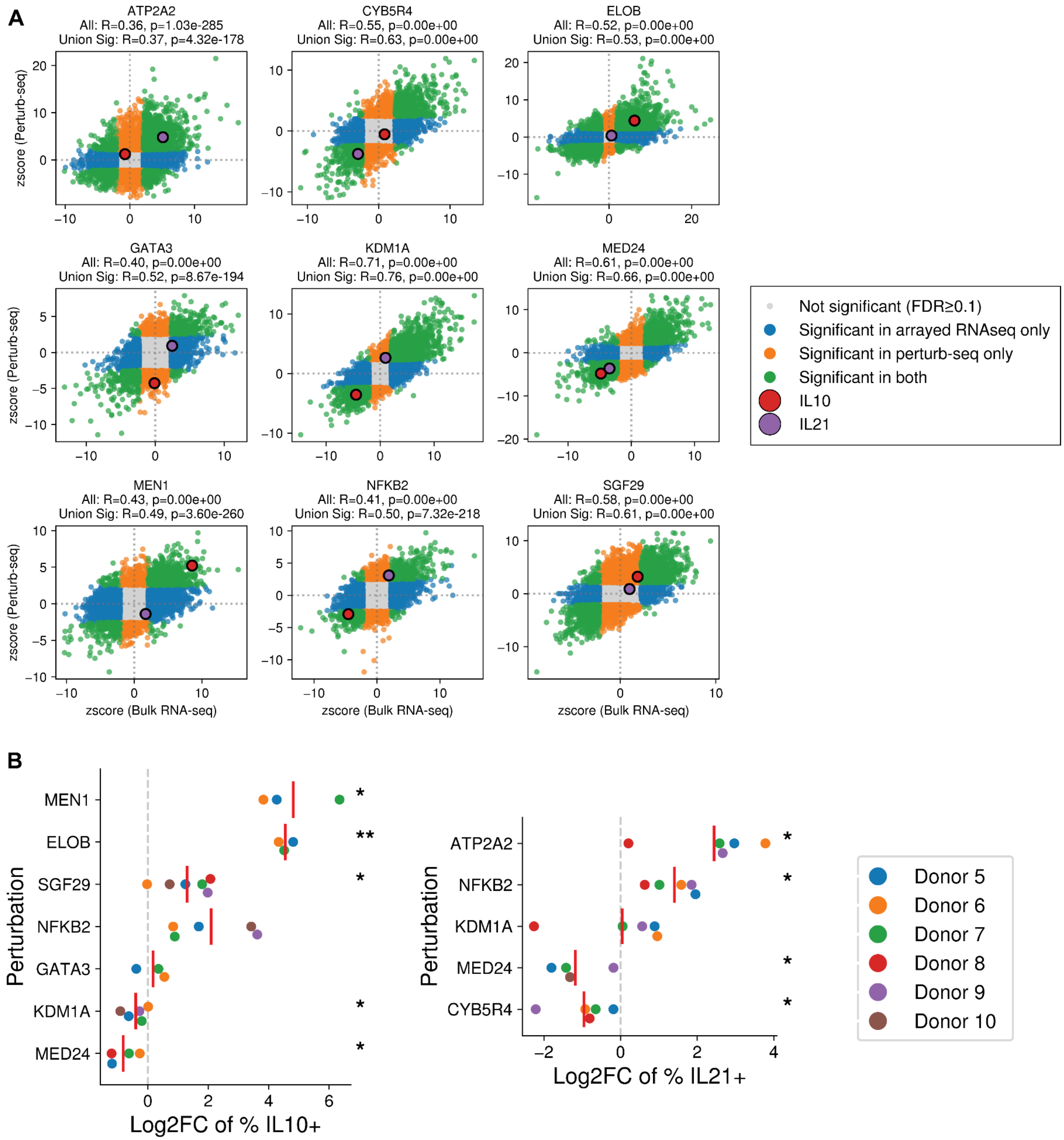


***Supplementary Figure 11. Arrayed validation of IL10/IL21 regulation.*** *(**A**) Scatter plots comparing the differential expression z-scores from arrayed bulk RNA-seq (x-axis) versus perturb-seq (y-axis) for the 9 selected regulators. Each point represents a measured gene. The z-scores for the on-target knockdown were masked (set to 0). Pearson correlation coefficient and corresponding p-values are shown for all shared genes ("All") and for the union of significantly differentially expressed genes between the two datasets ("Union Sig"). (**B**) Flow cytometry validation of regulatory effects on cytokine production. Swarmplots show the Log2 Fold Change (Log2FC) in the frequency of IL-10+ (left) or IL-21+ (right) cells following knockdown of the indicated regulators, normalized to non-targeting controls. Each dot represents cells from a unique human donor. Red vertical lines indicate the mean across donors. Statistical significance was determined using a one-sample t-test followed by Benjamini-Hochberg FDR correction (*FDR < 0.05, **FDR < 0.01). Flow cytometry plots used for quantification are in Suppl. Figure 12.*

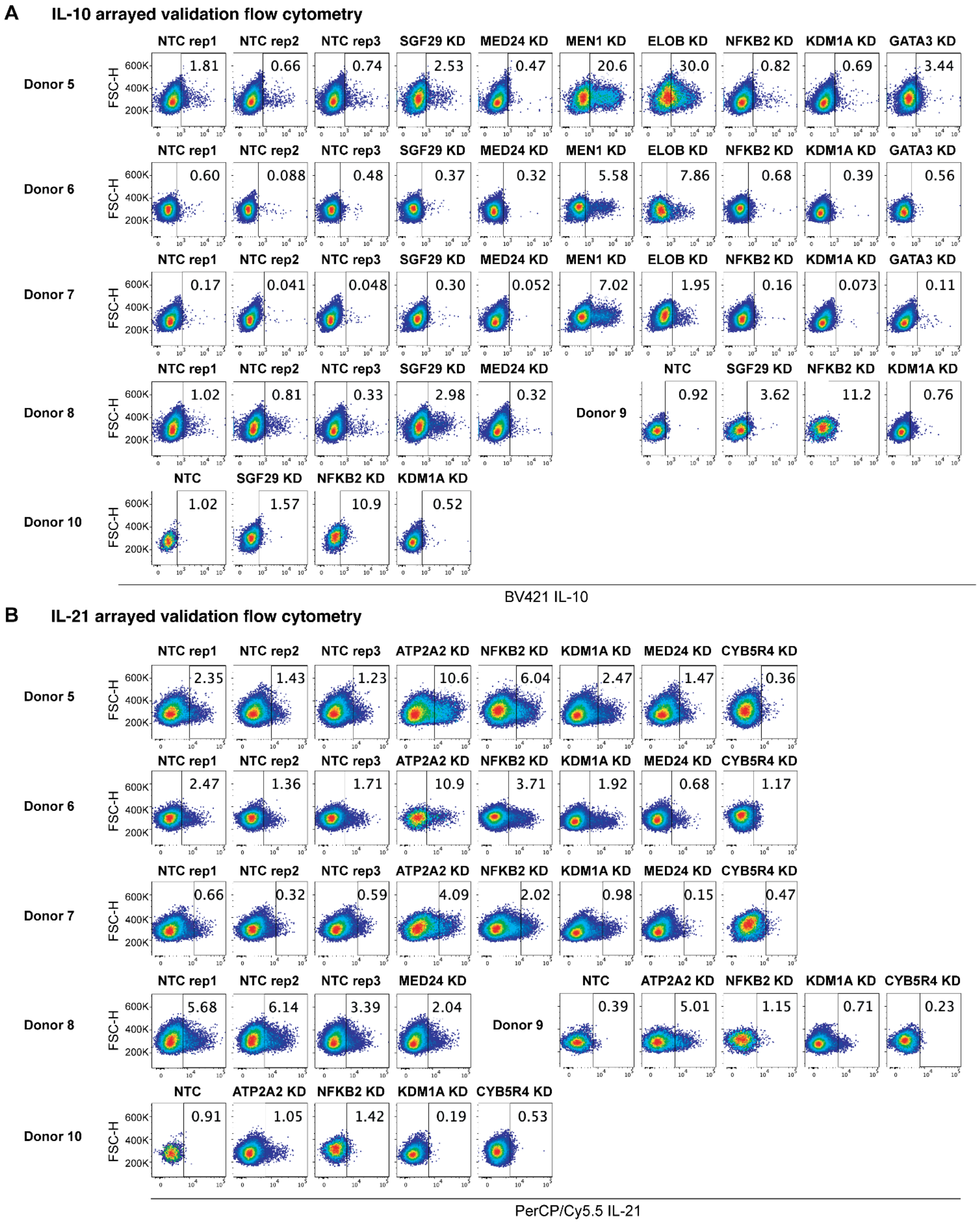


***Supplementary Figure 12*. *IL10/IL21 arrayed validation flow cytometry plots.*** *Some perturbations were not measured for donor 8-10 due to insufficient cell numbers for both flow cytometry and bulk RNAseq. NTC = non-targeting control. NTC, NTC1, NTC2 wells are transduced with a mixture of two non-targeting control gRNAs (NTC425 and NTC720) expressed from pRZ111 (mCherry marker), and NTC3 are transduced with a mixture of two non-targeting control gRNAs (NTC425 and NTC720) expressed from pRZ109 (GFP marker) (also see Methods* *“Arrayed validation of IL10/IL21 regulation”* *and Suppl. Table 2).*

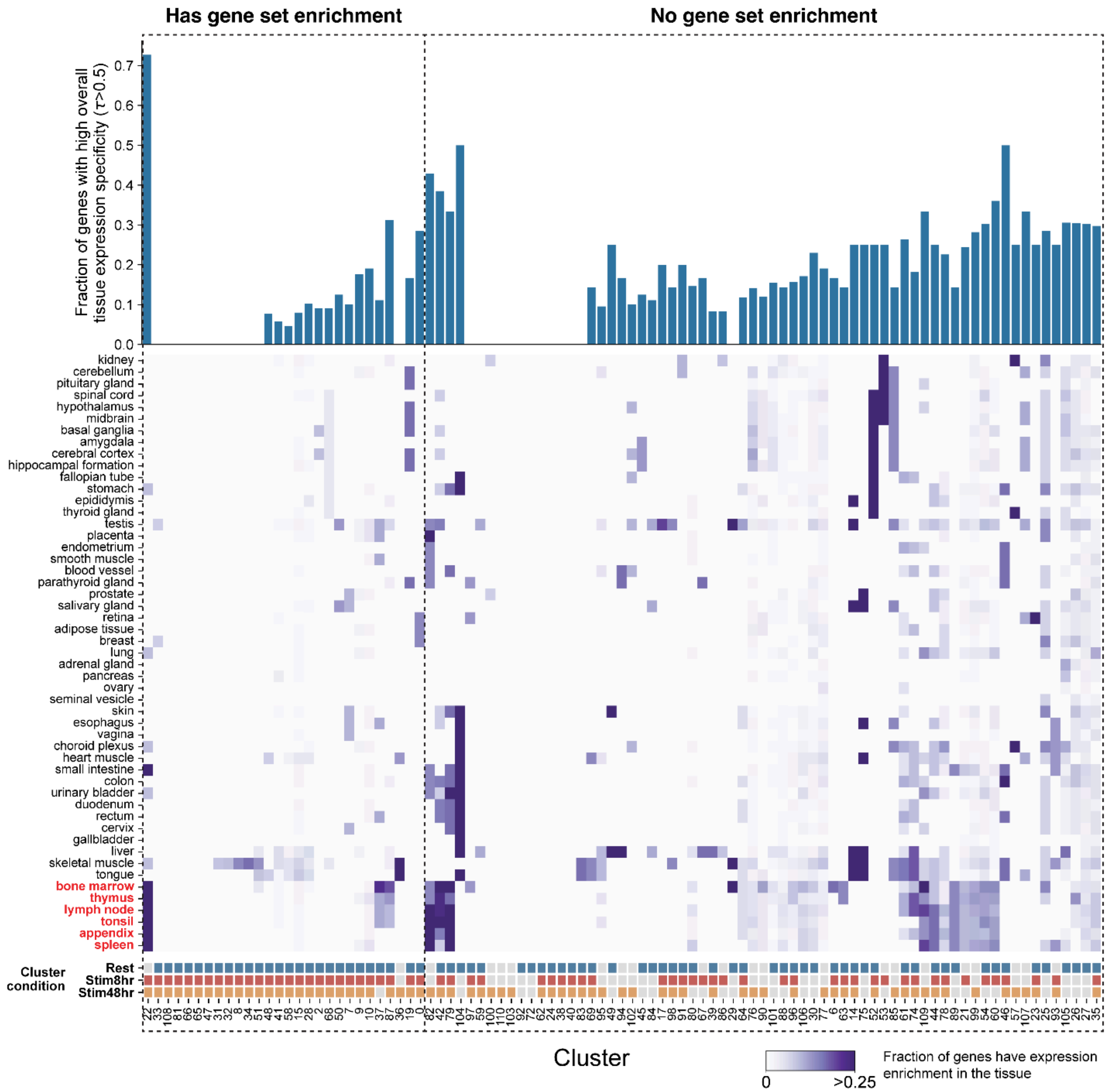


***Supplementary Figure 13. Tissue expression specificity and condition specificity of regulator clusters with or without gene set enrichment.*** *Regulator clusters with at least 4 regulator genes are shown here. They are separated into two groups based on whether they show significant gene set enrichment (left) or no detectable enrichment (right). "Has gene set enrichment" means regulators in those clusters have gene set enrichment with 5% FDR and 15% cluster regulator overlaps in at least one complex/pathway term from the CORUM, STRINGDB, KEGG, and Reactome databases (also see Methods "Annotation of clusters".* ***(Top)*** *For each cluster, the fraction of genes exhibiting high overall tissue specificity, quantified by Tau (τ) index.* ***(Middle)*** *Heatmap summarizing tissue-level expression enrichment for each cluster. Each block is the fraction of regulators within a given regulator cluster (column) that are enriched in a specific tissue (row) (see Methods "Tissue-specific gene expression analysis" for calculation of τ index and tissue expression enrichment). Tissue names labeled in red are lymphoid tissues or related to T cell development.* ***(Bottom)*** *Condition specificity of regulator clusters (see "Assessing condition specificity of clusters" section in Methods for classification of condition specificity).*

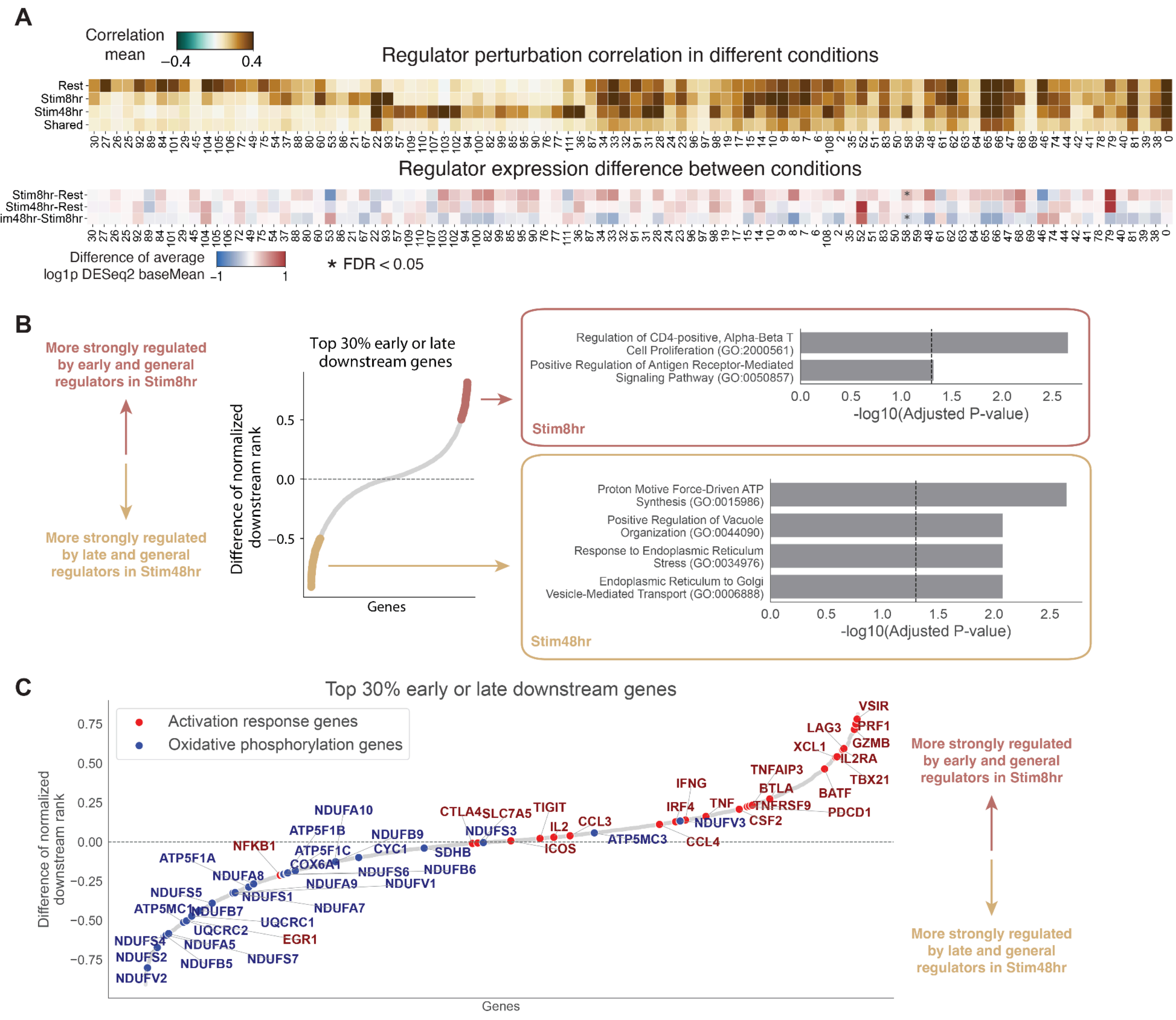


***Supplementary Figure 14. Condition specificity of regulator clusters. (A) (Top)*** *Heatmap for mean intra-condition correlation ("Rest", "Stim8hr", "Stim48hr") and inter-condition correlation ("Shared") for each cluster. Only clusters with at least 4 unique regulator genes are shown.* ***(Bottom)*** *Heatmap of the regulator expression differences between conditions for the same set of regulator clusters. For each regulator cluster (columns), the aggregate change in expression of its regulators was compared between two conditions. Expression values were derived from DESeq2 baseMean counts and log-transformed. The heatmap displays the difference in mean expression change for regulators in a specific cluster compared to the global background expression difference distribution of all regulators. Statistical significance was assessed using Welch's t-test with Benjamini-Hochberg FDR correction. Colors indicate the magnitude of the relative difference. * FDR<0.05.* ***(B) (Left)*** *Downstream genes were analyzed for early regulators and general regulators at Stim8hr ("early") and late regulators and general regulators at Stim48hr ("late"). For each condition, genes were ordered using the rank aggregation approach for ranking downstream genes of regulator clusters (Methods "Curation, ranking and annotation of downstream genes"). The ranks of downstream genes were normalized to a 0–1 scale (1: the most downregulated gene in rank by regulator knockdown). The scatter plot displays the difference in normalized ranks (early minus late) for genes in the top 30 percentile of ranking in either early or late downstream genes. Genes with a rank difference > 0.5 (red) are preferentially regulated in the early phase, while those with a difference < -0.5 (tan) are preferentially regulated in the late phase.* ***(Right)*** *Bar plots display 3 top significantly enriched gene ontology terms for early or late downstream genes, using union of all downstream genes as background gene set.* ***(C)*** *The same scatterplot as in (B) overlaid with curated activation response genes and oxidative phosphorylation genes from literature.*

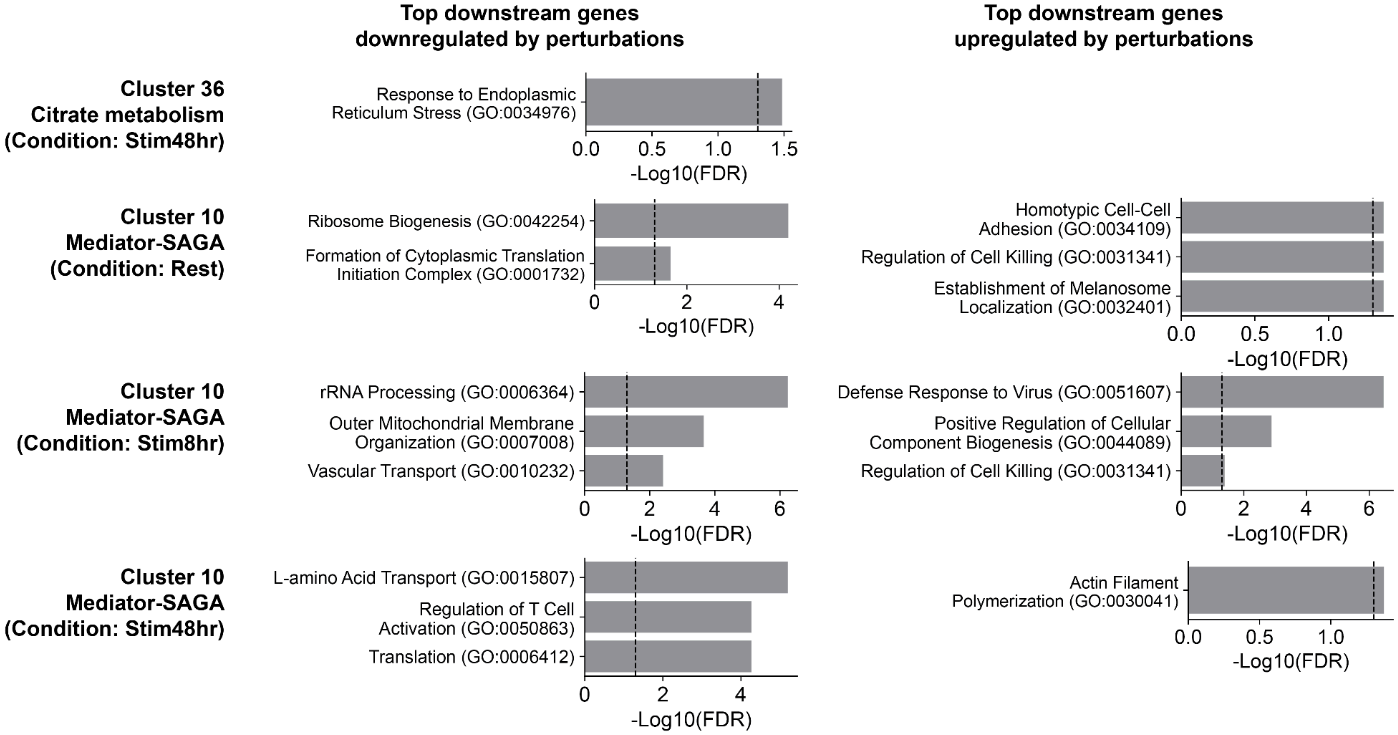


***Supplementary Figure 15**. **Downstream gene set annotation for selected clusters.** Gene Ontology (GO) enrichment analysis of downstream genes of selected clusters across indicated conditions. Bar plots display 3 top significantly enriched Biological Processes GO (2025 version) terms for downstream genes that are either downregulated (left panels) or upregulated (right panels) following the knockdown of the cluster's regulators. The x-axis represents the -Log10(FDR) significance of enrichment. The dashed vertical line denotes the significance threshold (FDR < 0.05).*

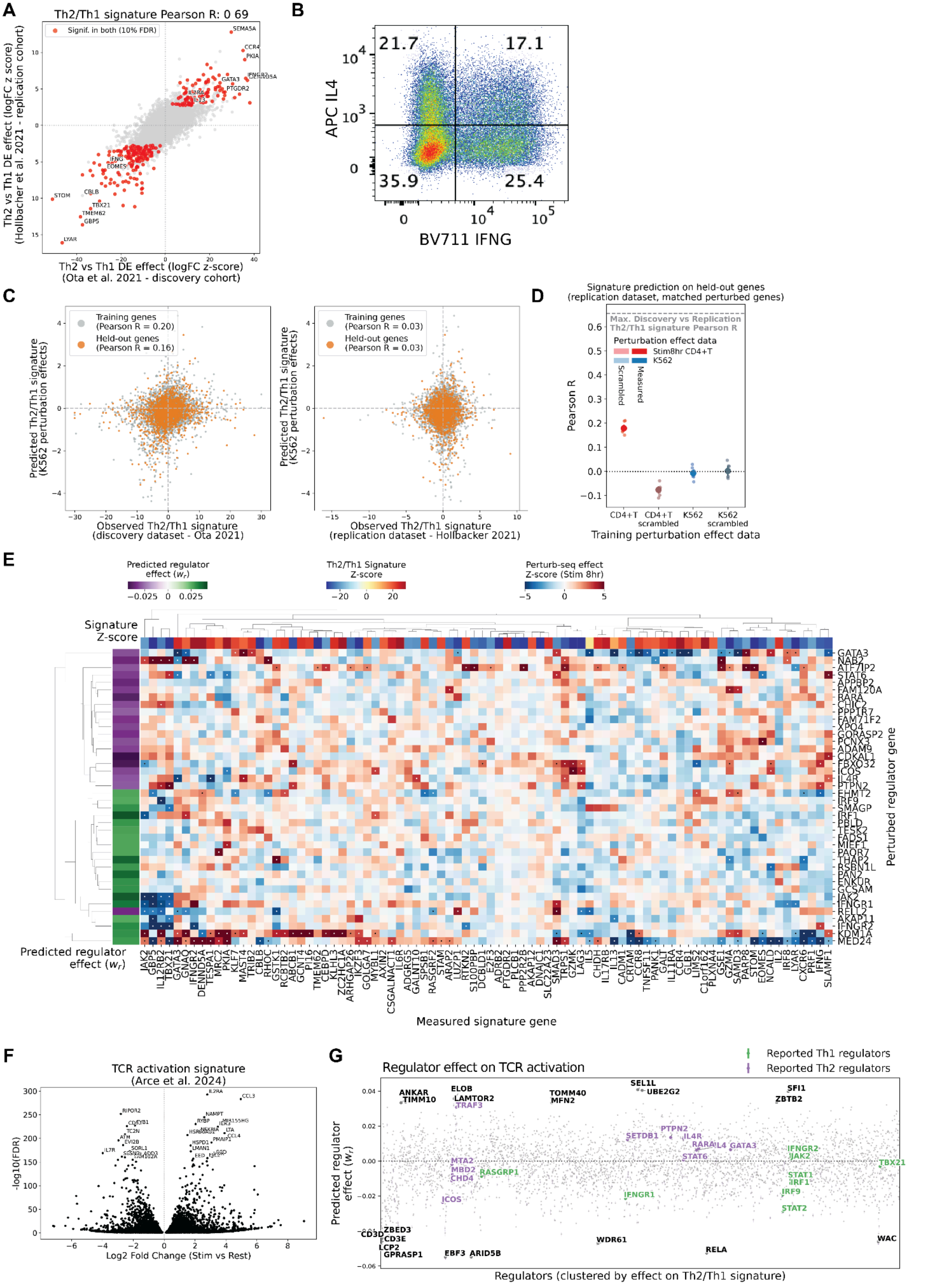
A
Th2/Th1 signature Pearson R: 0 69
Signif. in both (10% FDR)
Th2 vs Th1 DE effect (logFC z score) (Hollbacher et al. 2021 - replication cohort)
Th2 vs Th1 DE effect (logFC z-score) (Ota et al. 2021 - discovery cohort)
B
APC IL4
BV711 IFNG
21.7
17.1
35.9
25.4
C
Training genes (Pearson R = 0.20)
Held-out genes (Pearson R = 0.16)
Training genes (Pearson R = 0.03)
Held-out genes (Pearson R = 0.03)
Predicted Th2/Th1 signature (K562 perturbation effects)
Observed Th2/Th1 signature (discovery dataset - Ota 2021)
Observed Th2/Th1 signature (replication dataset - Hollbacker 2021)
D
Signature prediction on held-out genes (replication dataset, matched perturbed genes)
Max. Discovery vs Replication Th2/Th1 signature Pearson R
Perturbation effect data
Stim8hr CD4+T
K562
Scrambled
Measured
Pearson R
CD4+T
CD4+T scrambled
K562
K562 scrambled
Training perturbation effect data
E
Predicted regulator effect ($w_r$)
Th2/Th1 Signature Z-score
Perturb-seq effect Z-score (Stim 8hr)
Signature Z-score
Predicted regulator effect ($w_r$)
Perturbed regulator gene
Measured signature gene
GATA3
NAB2
ATF7IP2
STAT6
APPBP2
FAM120A
RARA
CHIC2
PPP1R7
FAM71F2
XPO4
GORASP2
PCNX3
ADAM9
CDKAL1
FBXO32
ICOS
IL4R
PTPN2
EHMT2
IRF9
SMAGP
IRF1
PBLD
TESK2
FADS1
MIEF1
PAQR7
THAP2
RSBN1L
PAN2
ENKUR
GCSAM
JAK2
IFNGR1
RELL2
AKAP11
IFNGR2
KDM1A
MED24
F
TCR activation signature (Arce et al. 2024)
-log10(FDR)
Log2 Fold Change (Stim vs Rest)
G
Regulator effect on TCR activation
Reported Th1 regulators
Reported Th2 regulators
Predicted regulator effect ($w_r$)
Regulators (clustered by effect on Th2/Th1 signature)

***Supplementary Figure 16. Predicting regulators of Th2/Th1 polarization.*** *(**A**) Correlation between Th2/Th1 differential expression results (logFC z-score) from the discovery cohort (Ota et al. 2021) and replication cohort (Hollbacher et al. 2021). Red points indicate significant genes in both cohorts (at 1% FDR); gray points represent all other genes. Selected signature genes are labeled. The Pearson correlation coefficient is reported. (**B**) Protein staining of IFN-γ (Brilliant Violet 711 anti-human IFN-γ Antibody, catalogue number 502540, Biolegend) and IL-4 (IL-4 Monoclonal Antibody (8D4-8), APC, catalogue number 17-7049-42, Life Tech) of NTC3 sample of donor 7 (*Suppl. Figure 15*). The experimental workflow is the same as "Arrayed validation of IL10/IL21 regulation" section in Methods except for the antibodies used for staining. (**C**) Reconstruction of polarization signatures from perturbation effects in CD4+ T cells (left) and K562 cells (right). The x-axis shows the observed Th2/Th1 signature; the y-axis shows the predicted signature from the model trained on the perturbation data. Orange points indicate held-out genes. Pearson correlation coefficients between true and predicted signatures are reported. Both models were trained only on 2190 regulators with significant effect in K562 perturb-seq (see Methods) (**D**) Evaluation of polarization signature prediction on held-out genes in the replication cohort for models trained only on 2190 regulators with significant effect in K562 perturb-seq: mean Pearson correlation coefficients (y-axis) across splits (5-fold cross-validation with 3 initializations) for model fit on Stim8hr CD4+T perturbation effects (red) and K562 perturbation effects dataset (blue, data from Replogle et al. 2022). Scrambled controls shown in light colors. Dotted line indicates maximum achievable correlation (inter-cohort agreement). Error bars: 95% CI across cross-validation splits. (**E**) Heatmap showing regulator knockdown effects on Th2/Th1 signature genes (showing top and bottom 50 significant genes by z-score, at 1% FDR) in perturb-seq data from Stim8hr condition. Rows represent regulators clustered by their effect on the Th2/Th1 signature; columns represent individual signature genes. Color intensity represents the magnitude and direction of predicted regulatory effects. Black dots denote where the trans effect is significant (10% FDR). Top annotations indicate signature Z-score. Left annotation shows the predicted regulator effect on the signature. (**F**) Volcano plot of TCR activation signature (Arce et al. 2025). Scatter plot shows log2 fold change (stimulated versus rest) plotted against -log10(FDR). (**G**) Predicted regulator effects (model coefficient $w_r$, y-axis) on TCR activation signature for all regulators. The mean and standard error for coefficients estimated on 15 different train-test splits are shown. Regulators are ordered by hierarchical clustering on the perturbation effects on Th2/Th1 signature genes (as in* Figure 4*F). Known Th1 and Th2 regulators highlighted in green and purple, respectively. Other top and bottom regulators without known effects on polarization are annotated in black.*

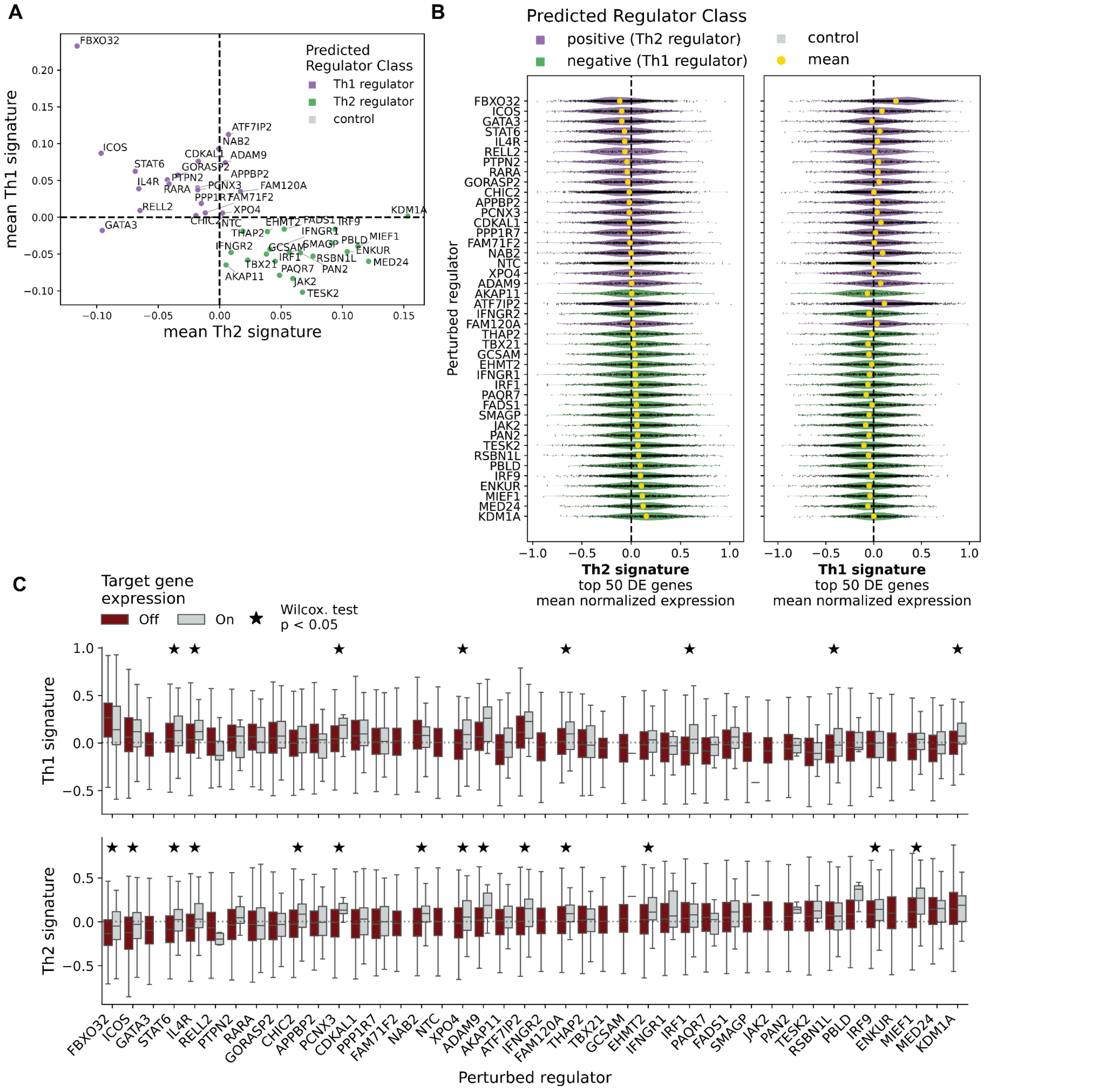


***Supplementary Figure 17. Effect of predicted polarization regulators on Th1 and Th2 genes.*** *(**A**) Mean Th2 signature (x-axis) and Th1 signature (y-axis) in cells harboring perturbation of putative Th1 and Th2 regulators. Color indicates predicted polarization effect (purple: Th2-promoting, green: Th1-promoting, gray: NTC) (**B**) Distributions of the Th2 (left) and Th1 (right) expression signature in cells with perturbation of top predicted polarization regulators. Regulators are ranked by mean signature score across cells (yellow dot). Color indicates predicted polarization effect (purple: Th2-promoting, green: Th1-promoting, gray: NTC). Signatures of NTC cells are shown only for cells with a subsample of non-targeting guides. (C) Distributions of the Th1 (top) and Th2 (bottom) expression signature in cells with perturbation of top predicted polarization regulators, stratified by expression of the target gene (where On: target gene normalized counts > 0). Stars denote where the difference in score is significant between cells with or without target gene expression (Wilcoxon rank-sum test).*

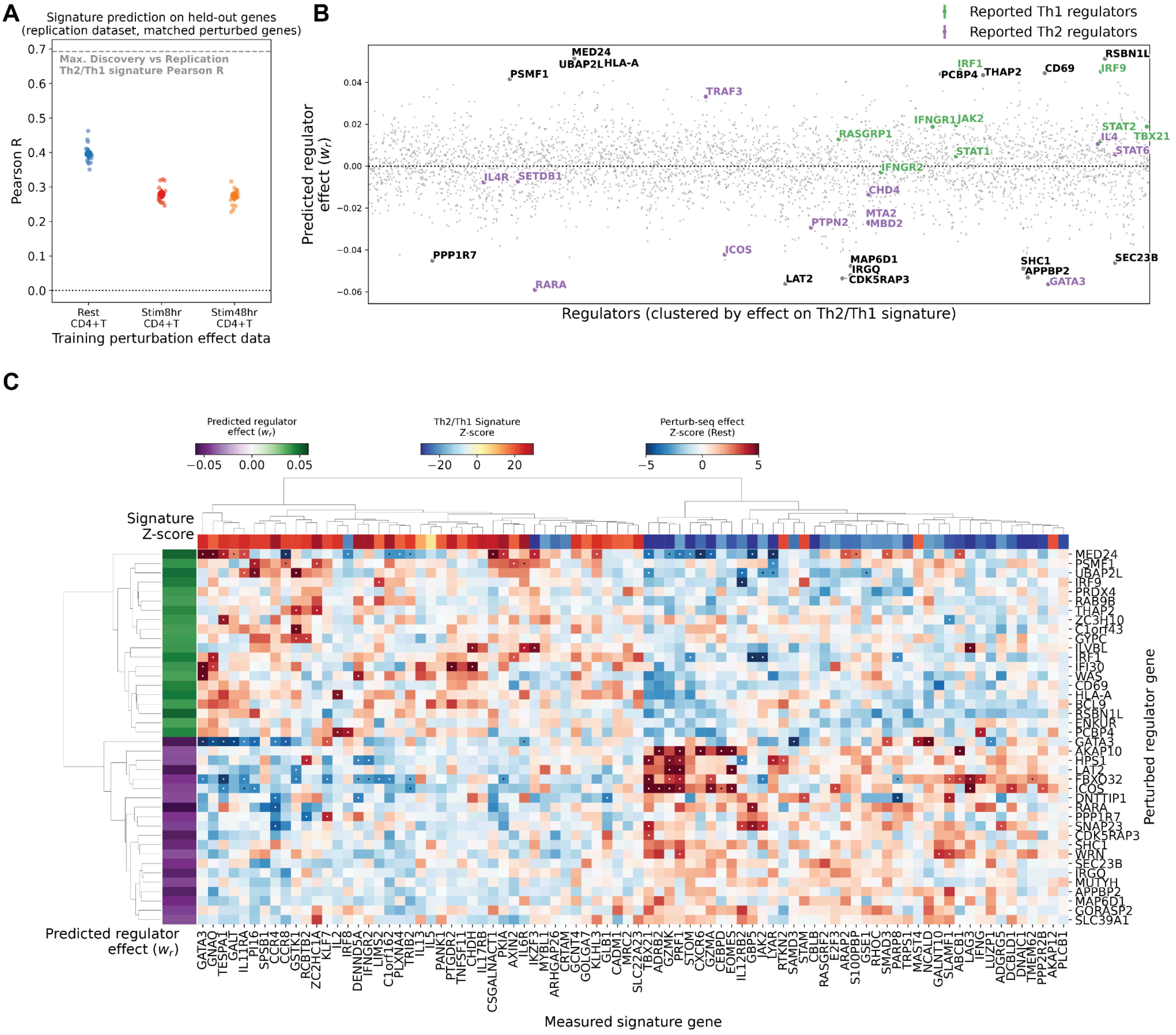


***Supplementary Figure 18**. **Predicting regulators of Th2/Th1 polarization across CD4+ T cell conditions.** (**A**) Evaluation of polarization signature prediction on held-out genes in the replication cohort for models trained on perturbation effects in different conditions (x-axis), measured with mean Pearson correlation coefficients (y-axis) across splits (5-fold cross-validation with 3 initializations). Dotted line indicates maximum achievable correlation (inter-cohort agreement). Error bars: 95% CI across cross-validation splits. (**B**) Predicted regulator effects (model coefficient $w_r$, y-axis) on polarization signature predicted from Rest condition. The mean and standard error for coefficients estimated on 15 different train-test splits are shown. Regulators are ordered by hierarchical clustering on the perturbation effects on Th2/Th1 signature genes (1% FDR). Known Th1 and Th2 regulators highlighted in green and purple, respectively. Top and bottom regulators without known effects on polarization are annotated in black. (**C**) As in* Suppl. Figure 16*E but showing predicted regulators and knockdown effects in perturb-seq data from Rest condition.*

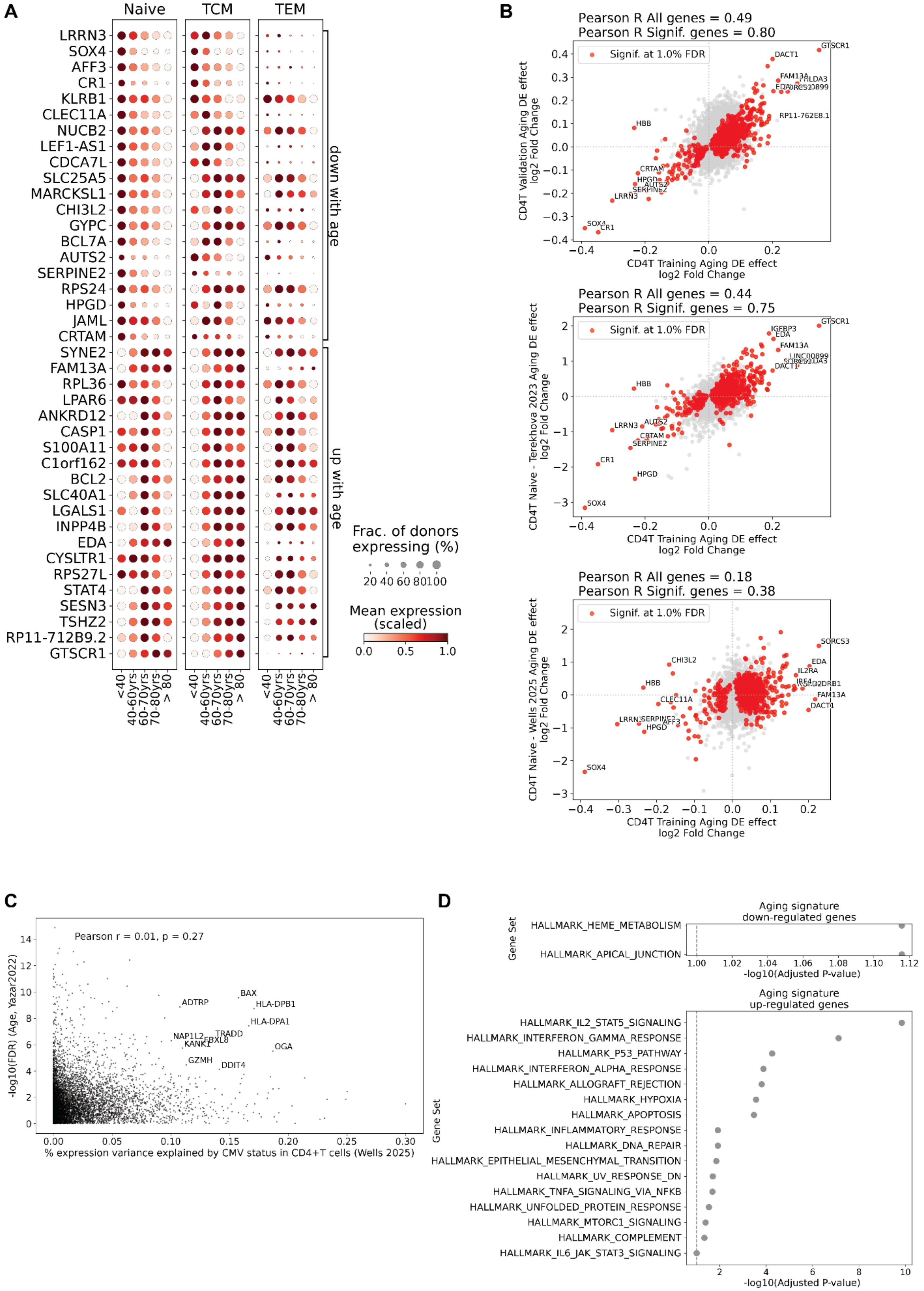
A
Naive
TCM
TEM
LRRN3
SOX4
AFF3
CR1
KLRB1
CLEC11A
NUCB2
LEF1-AS1
CDCA7L
SLC25A5
MARCKSL1
CHI3L2
GYPC
BCL7A
AUTS2
SERPINE2
RPS24
HPGD
JAML
CRTAM
SYNE2
FAM13A
RPL36
LPAR6
ANKRD12
CASP1
S100A11
C1orf162
BCL2
SLC40A1
LGALS1
INPP4B
EDA
CYSLTR1
RPS27L
STAT4
SESN3
TSHZ2
RP11-712B9.2
GTSCR1
down with age
up with age
<40
40-60yrs
60-70yrs
70-80yrs
> 80
Frac. of donors expressing (%)
20 40 60 80 100
Mean expression (scaled)
0.0 0.5 1.0
B
Pearson R All genes = 0.49
Pearson R Signif. genes = 0.80
Signif. at 1.0% FDR
CD4T Validation Aging DE effect log2 Fold Change
CD4T Training Aging DE effect log2 Fold Change
Pearson R All genes = 0.44
Pearson R Signif. genes = 0.75
CD4T Naive - Terekhova 2023 Aging DE effect log2 Fold Change
Pearson R All genes = 0.18
Pearson R Signif. genes = 0.38
CD4T Naive - Wells 2025 Aging DE effect log2 Fold Change
C
Pearson r = 0.01, p = 0.27
-log10(FDR) (Age, Yazar2022)
% expression variance explained by CMV status in CD4+T cells (Wells 2025)
BAX
ADTRP
HLA-DPB1
HLA-DPA1
NAP1L2
TRADD
FBXL8
KANK1
OGA
GZMH
DDIT4
D
Aging signature down-regulated genes
Gene Set
HALLMARK_HEME_METABOLISM
HALLMARK_APICAL_JUNCTION
-log10(Adjusted P-value)
Aging signature up-regulated genes
HALLMARK_IL2_STAT5_SIGNALING
HALLMARK_INTERFERON_GAMMA_RESPONSE
HALLMARK_P53_PATHWAY
HALLMARK_INTERFERON_ALPHA_RESPONSE
HALLMARK_ALLOGRAFT_REJECTION
HALLMARK_HYPOXIA
HALLMARK_APOPTOSIS
HALLMARK_INFLAMMATORY_RESPONSE
HALLMARK_DNA_REPAIR
HALLMARK_EPITHELIAL_MESENCHYMAL_TRANSITION
HALLMARK_UV_RESPONSE_DN
HALLMARK_TNFA_SIGNALING_VIA_NFKB
HALLMARK_UNFOLDED_PROTEIN_RESPONSE
HALLMARK_MTORC1_SIGNALING
HALLMARK_COMPLEMENT
HALLMARK_IL6_JAK_STAT3_SIGNALING

***Supplementary Figure 19*. Differential expression analysis of age-associated gene signature in CD4+ T cell subsets. (A)** *Dot plot showing expression patterns of top and bottom 20 age-associated DE genes (1% FDR) across CD4+ T cell subsets (Naive, TCM, TEM) and age bins (<40, 40-60, 60-70, 70-80, >80 years). Dot size represents the fraction of donors expressing each gene, and color intensity indicates mean log-normalized expression level, scaled between 0 and 1.* ***(B)*** *Comparison of age-associated differential expression signatures between the discovery cohort, the replication cohort and published datasets. Scatter plots show log2 fold changes for genes in the aging signature, comparing estimates from CD4+ T cells in the Yazar2022 replication cohort (top), CD4+ T cells from the Wells et al. 2025 (middle), and CD4+ T cells from Terekhova et al. 2023 (bottom) against the discovery cohort estimates. Red points indicate genes significant at 1% FDR in both datasets. Pearson correlation coefficients are shown for all genes and for significant genes in the discovery cohort.* ***(C)*** *Scatter plot comparing the percentage of expression variance explained by CMV status in CD4+ T cells ([64], x-axis) versus the FDR of differential expression with age (y-axis) for each gene. The Pearson correlation coefficient and p-value of the correlation are shown. Selected significant genes with high dependence on CMV status are labeled.* ***(D)*** *Gene set enrichment analysis showing Hallmark pathways significantly enriched among age-associated down-regulated genes (top) and up-regulated genes (bottom). The x-axis denotes -log10(adjusted p-value) for enrichment (using the EnrichR method as implemented in the gseapy package).*

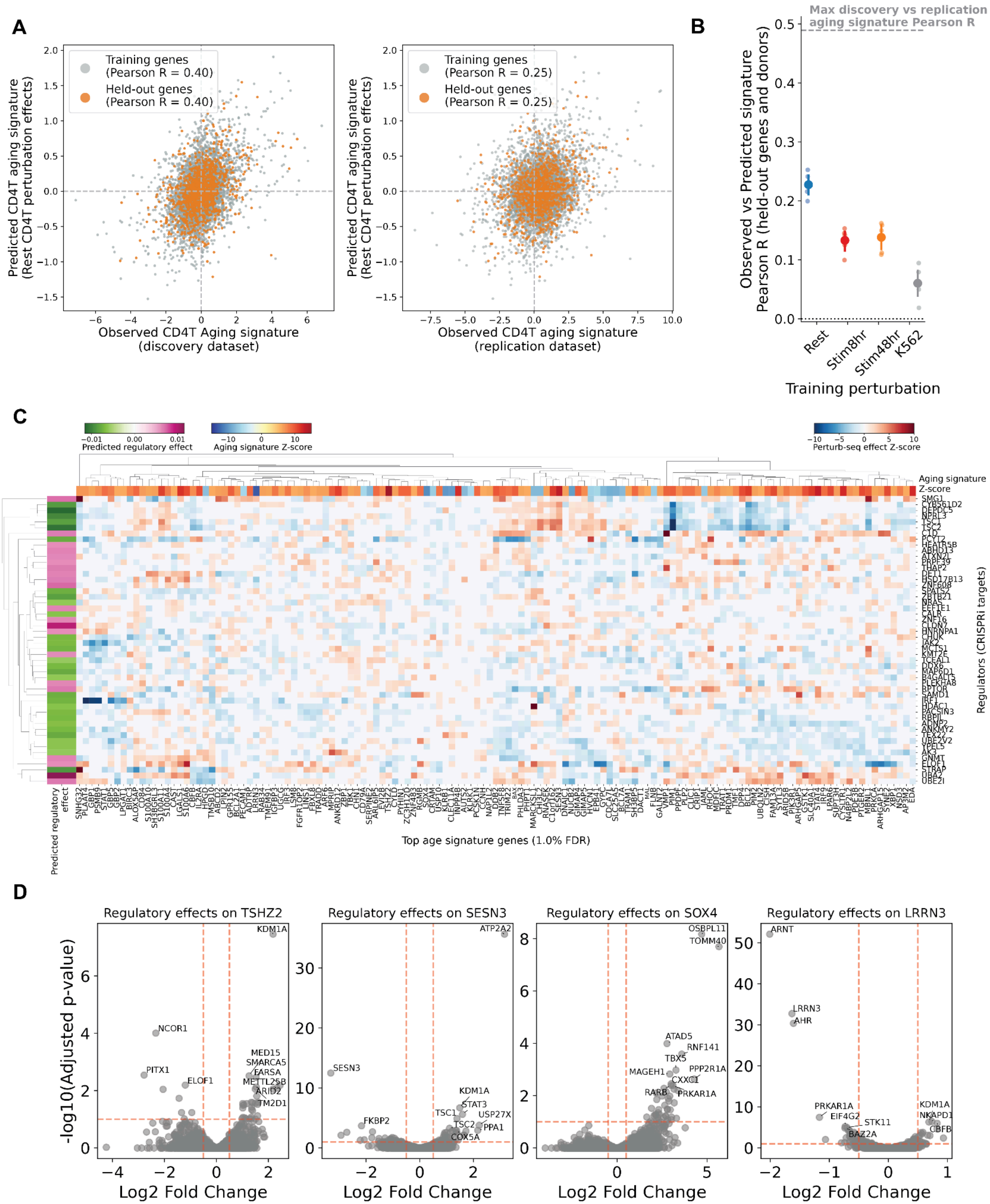
A
Training genes (Pearson R = 0.40)
Held-out genes (Pearson R = 0.40)
Predicted CD4T aging signature (Rest CD4T perturbation effects)
Observed CD4T Aging signature (discovery dataset)
Training genes (Pearson R = 0.25)
Held-out genes (Pearson R = 0.25)
Observed CD4T aging signature (replication dataset)
B
Max discovery vs replication aging signature Pearson R
Observed vs Predicted signature Pearson R (held-out genes and donors)
Rest
Stim8hr
Stim48hr
K562
Training perturbation
C
Predicted regulatory effect
Aging signature Z-score
Perturb-seq effect Z-score
Regulators (CRISPRi targets)
Top age signature genes (1.0% FDR)
D
Regulatory effects on TSHZ2
Regulatory effects on SESN3
Regulatory effects on SOX4
Regulatory effects on LRRN3
-log10(Adjusted p-value)
Log2 Fold Change

***Supplementary Figure 20. Predicting regulators of CD4+T cell aging gene signature.*** *(**A**) Reconstruction of aging signature from perturbation effects in Rest CD4+ T cells on the Discovery cohort signature (left) and Replication cohort signature (right). The x-axis shows the observed Th2/Th1 signature; the y-axis shows the predicted signature from the model trained on the perturbation data. Orange points indicate held-out genes. Pearson correlation coefficients between true and predicted signatures are reported. (**B**) Evaluation of aging signature prediction on held-out genes in the replication cohort for models trained on perturbation effects: mean Pearson correlation coefficients (y-axis) across splits (5-fold cross-validation) for model fit on Rest, Stim8hr and Stim48hr CD4+T perturbation effects and K562 perturbation effects (data from Replogle et al. 2022). Dotted line indicates maximum achievable correlation (inter-cohort agreement). Error bars: 95% CI across cross-validation splits. (**C**) Heatmap showing regulator knockdown effects on aging signature genes (showing top and bottom 50 significant genes by z-score, at 1% FDR) in perturb-seq data from Rest condition. Rows represent regulators clustered by their effect on the aging signature; columns represent individual signature genes. Color intensity represents the magnitude and direction of predicted regulatory effects. Top annotations indicate signature Z-score. Left annotation shows the predicted regulator effect on the signature. (**D**) Volcano plots of trans effects of tested regulators on top DE genes with age (TSHZ2, SESN3, SOX4, LRRN3) in Rest CD4+T perturb-seq. Each point represents a perturbed gene.*

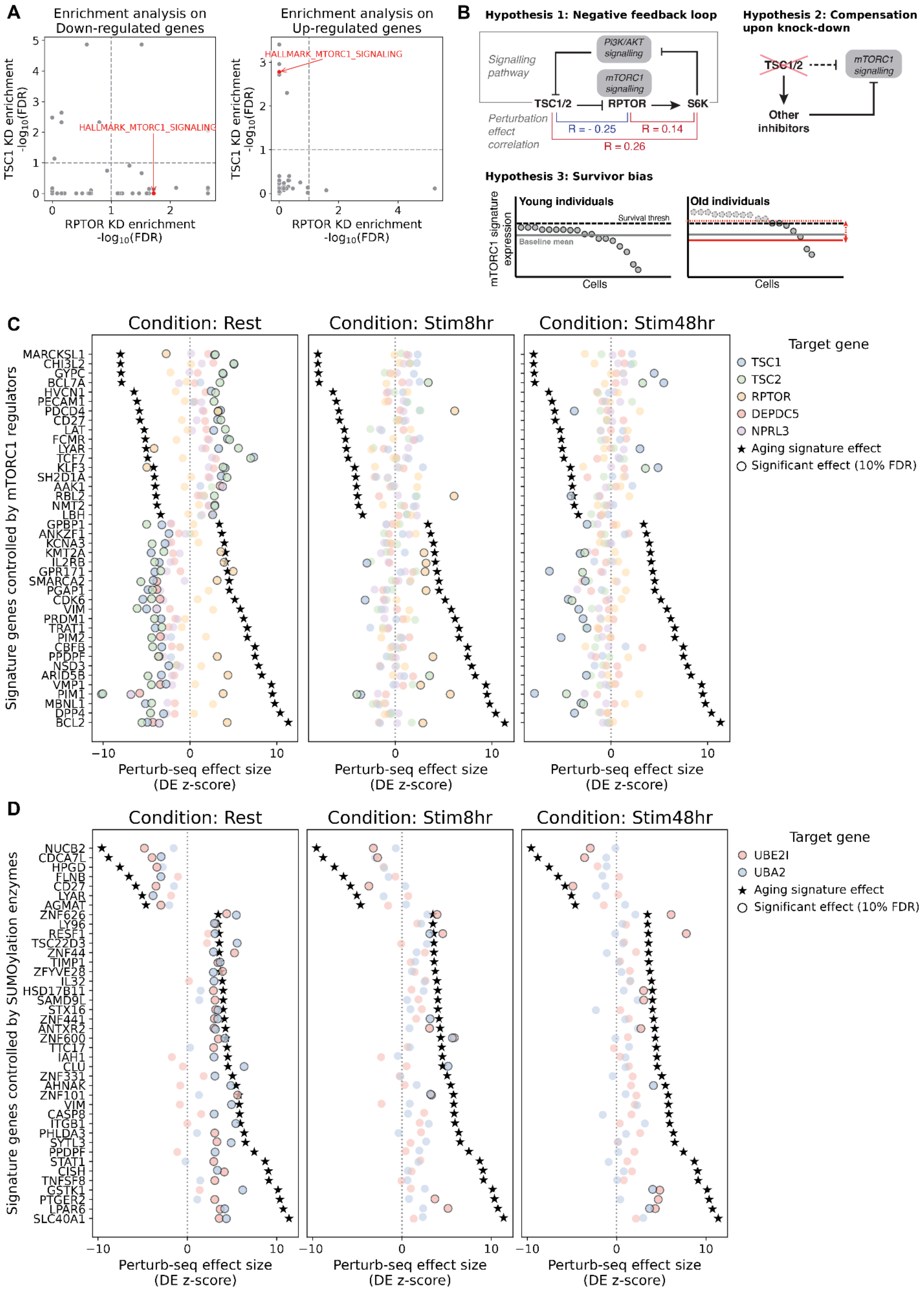
A
Enrichment analysis on Down-regulated genes
Enrichment analysis on Up-regulated genes
HALLMARK_MTORC1_SIGNALING
HALLMARK_MTORC1_SIGNALING
TSC1 KD enrichment -log10(FDR)
RPTOR KD enrichment -log10(FDR)
B
Hypothesis 1: Negative feedback loop
Signalling pathway
PI3K/AKT signalling
mTORC1 signalling
TSC1/2
RPTOR
S6K
Perturbation effect correlation
R = - 0.25
R = 0.14
R = 0.26
Hypothesis 2: Compensation upon knock-down
TSC1/2
mTORC1 signalling
Other inhibitors
Hypothesis 3: Survivor bias
mTORC1 signature expression
Young individuals
Survival thresh
Baseline mean
Cells
Old individuals
Cells
C
Condition: Rest
Condition: Stim8hr
Condition: Stim48hr
Signature genes controlled by mTORC1 regulators
MARCKSL1
CHI3L2
GYPC
BCL7A
HVCN1
PECAM1
PDCD4
CD27
LAT
FCMR
LYAR
TCF7
KLF3
SH2D1A
AAK1
RBL2
NMT2
LBH
GPBP1
ANKZF1
KCNA3
KMT2A
IL2RB
GPR171
SMARCA2
PGAP1
CDK6
VIM
PRDM1
TRAT1
PIM2
CBFB
PPDPF
NSD3
ARID5B
VMP1
PIM1
MBNL1
DPP4
BCL2
Perturb-seq effect size (DE z-score)
Target gene
TSC1
TSC2
RPTOR
DEPDC5
NPRL3
Aging signature effect
Significant effect (10% FDR)
D
Condition: Rest
Condition: Stim8hr
Condition: Stim48hr
Signature genes controlled by SUMOylation enzymes
NUCB2
CDCA7L
HPGD
FLNB
CD27
LYAR
AGMAT
ZNF626
LY96
RESF1
TSC22D3
ZNF44
TIMP1
ZFYVE28
IL32
HSD17B11
SAMD9L
STX16
ZNF441
ANTXR2
ZNF600
TTC17
IAH1
CLU
ZNF331
AHNAK
ZNF101
VIM
CASP8
ITGB1
PHLDA3
SYTL3
PPDPF
STAT1
CISH
TNFSF8
GSTK1
PTGER2
LPAR6
SLC40A1
Perturb-seq effect size (DE z-score)
Target gene
UBE2I
UBA2
Aging signature effect
Significant effect (10% FDR)

***Supplementary Figure 21. Predicted regulators of CD4+ T cell aging signature.*** *(**A**) Comparison of gene set enrichment analysis results (-log10(FDR)) on knockdown of mTORC1 activator RPTOR (x-axis) and mTORC1 inhibitor TSC1 (y-axis). Points represent tested gene sets from the MSigDB Hallmark gene sets. Enrichments on down-regulated genes (left) and up-regulated genes (right) are shown. (**B**) Illustration of hypotheses for discordant regulator effect prediction on mTORC1 pathway components. (**C-D**) Scatterplots of perturbation effects (DE log-fold change z-score, x-axis) of predicted regulators of aging signatures (y-axis) on aging signature genes controlled by regulators (see Methods). Effects across conditions are shown. The star denotes the DE effect (z-score) associated with age in the Discovery cohort. The black outline denotes where the trans effects are significant. We report effects on genes controlled by mTORC1 pathway regulators (C) and SUMOylation enzymes (D).*

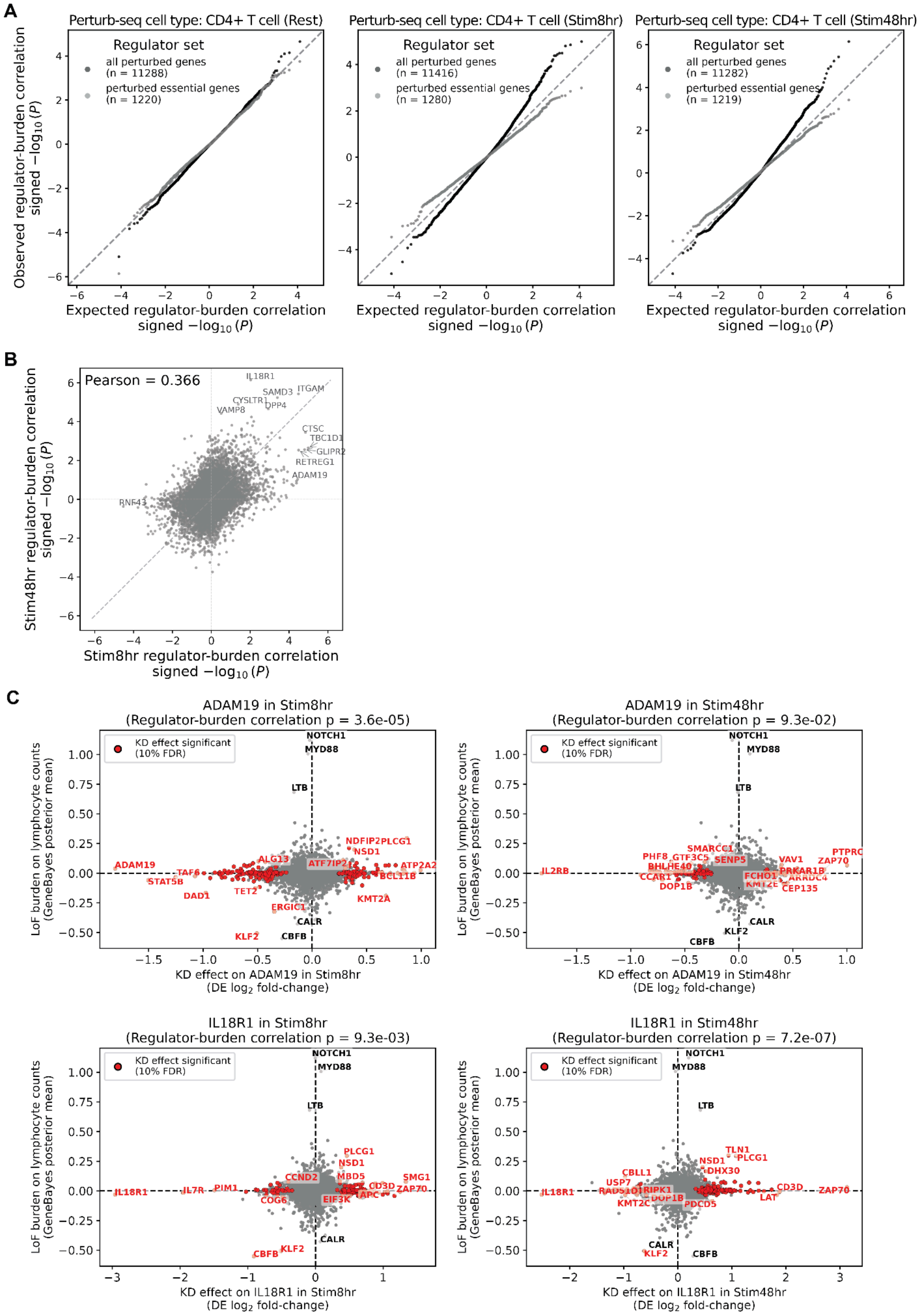
A
Perturb-seq cell type: CD4+ T cell (Rest)
Perturb-seq cell type: CD4+ T cell (Stim8hr)
Perturb-seq cell type: CD4+ T cell (Stim48hr)
Regulator set
all perturbed genes (n = 11288)
perturbed essential genes (n = 1220)
all perturbed genes (n = 11416)
perturbed essential genes (n = 1280)
all perturbed genes (n = 11282)
perturbed essential genes (n = 1219)
Observed regulator-burden correlation signed −log10 (P)
Expected regulator-burden correlation signed −log10 (P)
B
Pearson = 0.366
IL18R1
SAMD3
ITGAM
CYSLTR1
DPP4
VAMP8
CTSC
TBC1D1
GLIPR2
RETREG1
ADAM19
RNF43
Stim48hr regulator-burden correlation signed −log10 (P)
Stim8hr regulator-burden correlation signed −log10 (P)
C
ADAM19 in Stim8hr (Regulator-burden correlation p = 3.6e-05)
ADAM19 in Stim48hr (Regulator-burden correlation p = 9.3e-02)
IL18R1 in Stim8hr (Regulator-burden correlation p = 9.3e-03)
IL18R1 in Stim48hr (Regulator-burden correlation p = 7.2e-07)
KD effect significant (10% FDR)
LoF burden on lymphocyte counts (GeneBayes posterior mean)
KD effect on ADAM19 in Stim8hr (DE log2 fold-change)
KD effect on ADAM19 in Stim48hr (DE log2 fold-change)
KD effect on IL18R1 in Stim8hr (DE log2 fold-change)
KD effect on IL18R1 in Stim48hr (DE log2 fold-change)

***Supplementary Figure 22*. Regulator-burden correlation on lymphocyte traits.** *(**A**) QQ plot comparing expected vs observed signed p-values for regulator-burden correlations for LoF burden on lymphocyte counts in the UK biobank (UKBB) and regulatory effects estimated from perturb-seq data using all perturbed genes (black) or perturbed essential genes (grey, matching Jurkat dataset from Nadig et al. [106]). The sign is determined by the direction of correlation. Each dot represents one measured gene. Analysis on perturb-seq data from different CD4+T conditions are shown. (**B**) Comparison of signed p-values for regulator-burden correlations estimated on perturbation effects in Stim8hr condition (x-axis) and Stim48hr condition (y-axis). Top significant genes are annotated. The Pearson correlation coefficient is reported on top. (**C**) Scatterplots of knock-down (KD) effects (x-axis) and loss-of-function burden on Lymphocyte Counts (y-axis) of all regulators on a measured gene of interest. Results for ADAM19 (top row, putative core gene in Stim8hr condition) and IL18R1 (bottom row, putative core gene in Stim48hr condition) are shown, for KD effects in both conditions (columns). The p-value of the regulator-burden correlation for each comparison is reported. Red dots denote where the knock-down effects are significant.*

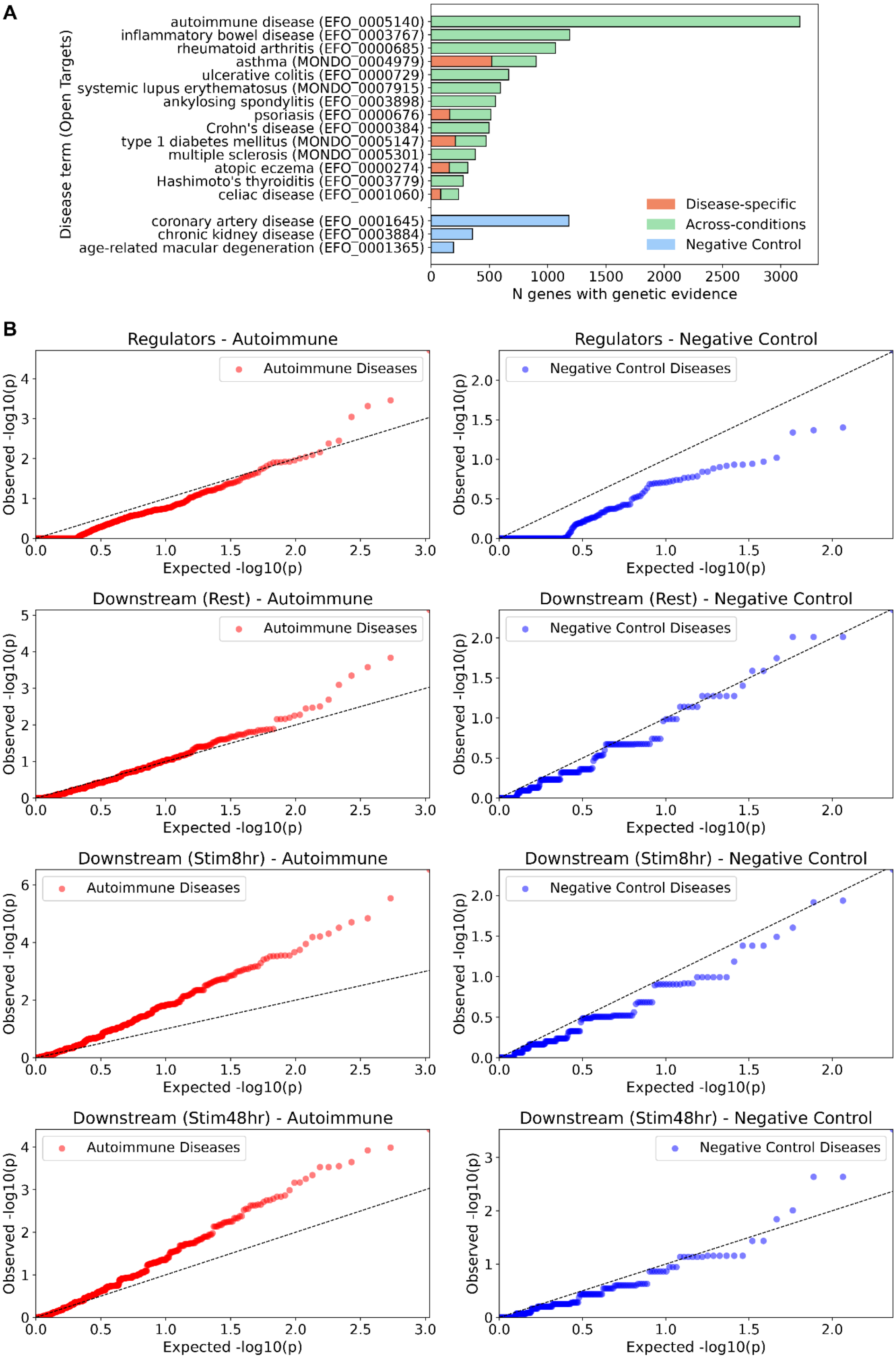
A
Disease term (Open Targets)
autoimmune disease (EFO_0005140)
inflammatory bowel disease (EFO_0003767)
rheumatoid arthritis (EFO_0000685)
asthma (MONDO_0004979)
ulcerative colitis (EFO_0000729)
systemic lupus erythematosus (MONDO_0007915)
ankylosing spondylitis (EFO_0003898)
psoriasis (EFO_0000676)
Crohn's disease (EFO_0000384)
type 1 diabetes mellitus (MONDO_0005147)
multiple sclerosis (MONDO_0005301)
atopic eczema (EFO_0000274)
Hashimoto's thyroiditis (EFO_0003779)
celiac disease (EFO_0001060)
coronary artery disease (EFO_0001645)
chronic kidney disease (EFO_0003884)
age-related macular degeneration (EFO_0001365)
Disease-specific
Across-conditions
Negative Control
N genes with genetic evidence
B
Regulators - Autoimmune
Regulators - Negative Control
Downstream (Rest) - Autoimmune
Downstream (Rest) - Negative Control
Downstream (Stim8hr) - Autoimmune
Downstream (Stim8hr) - Negative Control
Downstream (Stim48hr) - Autoimmune
Downstream (Stim48hr) - Negative Control
Autoimmune Diseases
Negative Control Diseases
Observed -log10(p)
Expected -log10(p)

***Supplementary Figure 23. GWAS gene enrichment analysis across regulator clusters.*** *(**A**) Number of genes with genetic evidence for association with each disease term retrieved from the OpenTargets Platform. Bars show gene counts for genes with disease-specific evidence (orange) and evidence shared across multiple autoimmune conditions (captured in general "autoimmune disease" term, in green). Disease identifiers from the Experimental Factor Ontology (EFO) or Mondo Disease Ontology (MONDO) are indicated in parentheses. (**B**) Quantile-quantile (QQ) plots comparing observed versus expected −log10(p-values) from Fisher's exact tests for enrichment of disease-associated genes. Left column: autoimmune diseases (red); right column: negative control diseases (blue). Rows show enrichment tests for cluster regulators (top) and downstream target genes in Rest condition, Stim8hr condition, and Stim48hr condition. Dashed diagonal lines indicate the expected distribution under the null hypothesis of no enrichment.*

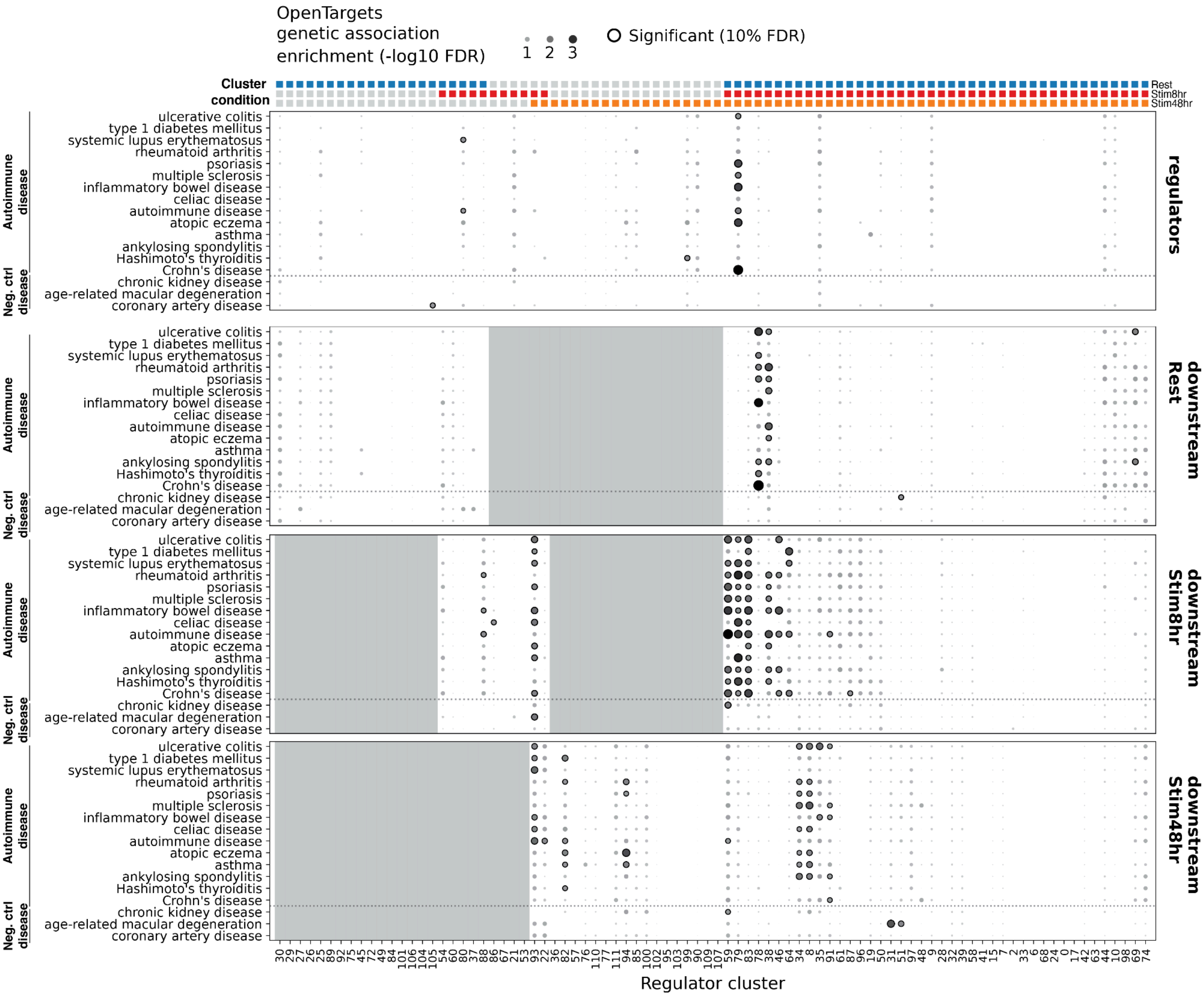


***Supplementary Figure 24. Enrichment of autoimmune disease GWAS genes across regulatory clusters and their downstream targets (related to*** Figure 7***A).*** *Dot plot showing enrichment of disease-associated genes within regulatory clusters (n=77) and their downstream target genes. Rows represent disease terms, grouped into autoimmune diseases and negative control non-immune diseases. Columns represent regulatory clusters, ordered by the condition in which each cluster was identified (indicated by the colored bar at top). Panels show enrichment results for cluster regulators (top) and downstream genes in each condition. Dot size corresponds to enrichment significance (−log10 FDR-adjusted p-value). Black-outlined dots indicate significant enrichments at 10% FDR. Gray shading indicates conditions where downstream gene enrichment was not tested due to lack of coordinated regulator effects.*

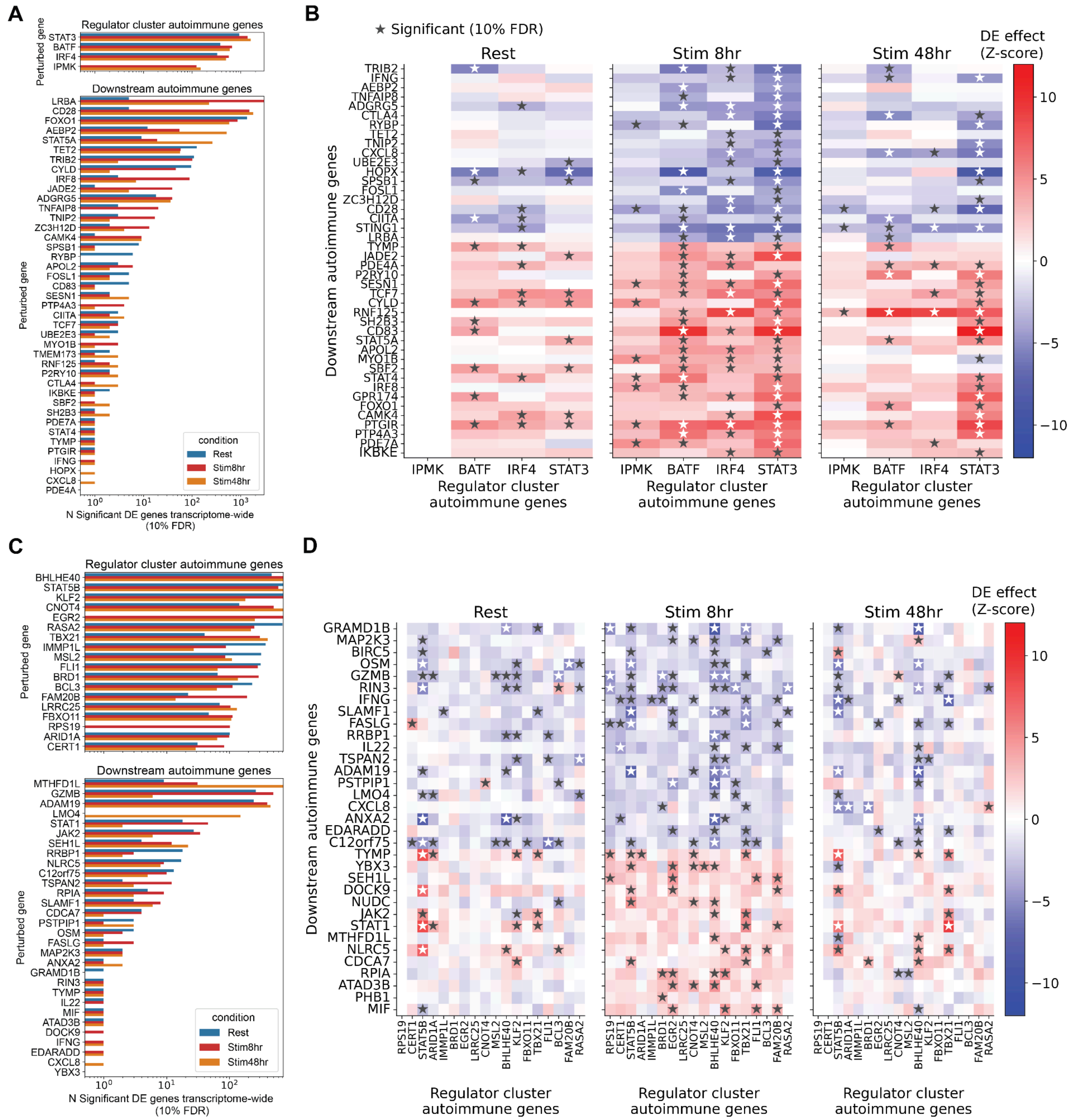


***Supplementary Figure 25. Effects of perturbations of autoimmune disease-associated genes in regulator clusters (related to*** Figure 7***C-D).*** *(**A, C**) perturbation effects of autoimmune disease-associated genes identified through enrichment analysis for cluster 79 (A) and cluster 80 (C), showing the number of significant DE genes (10% FDR) per perturbed gene across all three culture conditions (Rest, Stim8hr, Stim48hr). Only regulators (top) or downstream genes (bottom) from the cluster that are also associated with autoimmune diseases (based on genetic evidence from OpenTargets) are shown. (**B, D**) Heatmaps showing the differential expression effect (LFC z-score) of regulator autoimmune genes perturbations (x-axis) on downstream autoimmune disease genes (y-axis) across three*

*culture conditions (Rest, Stim8hr, Stim48hr) for cluster 79 (B) and cluster 80 (D). Stars indicate statistically significant differential expression (10% FDR). Missing values are filled with zero.*

#### Supplementary tables

**Supplementary Table 1** Sample metadata

- **cell_sample_id**: Unique identifier for each cell sample, combining 10x run ID, donor number, and culture condition (e.g., CD4i_R1_D1_Rest)
- **10xrun_id**: Identifier for the 10x Genomics sequencing run batch (e.g., CD4i_R1, CD4i_R2)
- **donor_id**: Unique identifier for the blood donor (format: CE#######)
- **culture_condition**: Experimental condition applied to cells (Rest, Stim8hr, or Stim48hr)
- **library_id**: Unique library identifier combining sample ID and sequencing run information
- **library_prep_kit**: 10x Genomics library preparation kit used (GEMX_flex_v2)
- **probe_hyb_loading**: Details of probe hybridization including cell count, probe volume, and barcode IDs for GEX and CRISPR probes
- **GEM_loading**: Number of cells loaded per Gel Bead-in-Emulsion (GEM)
- **sequencing_platform**: Sequencing platform used (Ultima)
- **age**: Donor age in years
- **sex**: Donor biological sex (Male/Female)
- **ethnicity**: Donor self-reported ethnicity
- **weight_kg**: Donor weight in kilograms
- **height_cm**: Donor height in centimeters
- **smoker**: Donor smoking status (Yes/No)
- **blood_type**: Donor ABO and Rh blood type
- **anticoagulant**: Anticoagulant used during blood collection (ACDA)
- **harvest_date**: Date of blood sample collection (M/D/YY format)

**Supplementary Table 2** Plasmid constructs used in this study

**Supplementary Table 3** gRNA guide library metadata

- **sgRNA**: Unique identifier for the guide RNA
- **chromosome**: Chromosome of the target site
- **pos**: Genomic position of the guide target site
- **strand**: DNA strand orientation of the target site (+ or -)
- **seq**: Full guide RNA sequence
- **seq_last19bp**: Last 19 base pairs of the guide sequence
- **PAM**: boolean flag for presence of Protospacer Adjacent Motif sequence
- **note**: Additional notes about the guide alignment to the genome
- **flag**: Quality control flag
- **target_gene_name_from_sgRNA**: Target gene name derived from the gRNA identifier
- **designed_target_gene_id**: Ensembl gene ID of the intended target gene (as designed)
- **designed_target_gene_name**: Gene name of the intended target gene (as designed)
- **target_gene_id**: Ensembl gene ID of the target gene, empty if intended target gene was not found near alignment site
- **target_gene_name**: Gene name of the target gene, empty if intended target gene was not found near alignment site
- **distance_to_closest_target_tss**: Distance (in base pairs) from guide to the closest transcription start site (TSS) of the target gene

- **nearby_gene_within_2kb**: list of genes within 2 kb of the guide target site
- **nearby_gene_within_30kb**: list of genes within 30 kb of the guide target site
- **nearest_within2kb_gene_id**: Ensembl gene ID of the nearest gene within 2 kb
- **nearest_within2kb_gene_name**: Gene name of the nearest gene within 2 kb
- **nearest_within2kb_gene_dist**: Distance to the nearest gene within 2 kb
- **nearest_within2kb_nontarget_gene_id**: Ensembl gene ID of the nearest non-target gene within 2 kb
- **nearest_within2kb_nontarget_gene_name**: Gene name of the nearest non-target gene within 2 kb
- **nearest_within2kb_nontarget_gene_dist**: Distance to the nearest non-target gene within 2 kb
- **putative_bidirectional_promoter**: Flag indicating potential bidirectional promoter region (may affect multiple genes)
- **other_alignment_chromosome**: Chromosome with potential off-target alignment
- **other_alignment_pos**: Genomic position of potential off-target alignment

**Supplementary Table 4** Summary statistics on knockdown efficiency of each gRNA guide across three culture conditions.

- **index**: gRNA ID
- **guide_mean_expr**: Mean log-normalized expression of the target gene in cells carrying this guide
- **guide_std_expr**: Standard deviation of log-normalized target gene expression in cells carrying this guide (set to 0.01 for guides with zero variance, 100 for guides with only one cell)
- **guide_n**: Number of cells carrying this guide
- **ntc_mean_expr**: Mean log-normalized expression of the target gene in non-targeting control cells
- **ntc_std_expr**: Standard deviation of log-normalized target gene expression in non-targeting control cells
- **ntc_n**: Total number of non-targeting control cells across all samples
- **t_statistic**: Welch's t-test statistic comparing guide expression vs NTC expression (negative values indicate knockdown)
- **p_value**: Nominal p-value from Welch's t-test
- **adj_p_value**: Benjamini-Hochberg FDR-adjusted p-value (minimum value capped at 1e-16)
- **signif_knockdown**: Boolean indicating significant knockdown (adj_p_value < 0.1 AND t_statistic < 0)
- **perturbed_gene_id**: Ensembl gene ID of the target gene
- **rank**: Rank of the target gene based on mean expression in NTC cells (1 = lowest expressed)
- **high_confidence_no_effect_guides**: Boolean indicating guides with high confidence of having no knockdown effect (criteria: non-significant knockdown, >10 cells with guide, target expression in NTCs >0.001)
- **culture_condition**: Culture condition for this measurement (Rest, Stim8hr, or Stim48hr)

**Supplementary Table 5** Perturbation effects summary statistics

- **target_contrast_gene_name**: Name of the perturbed gene
- **culture_condition**: culture condition (Rest, Stim8hr, Stim48hr)
- **target_contrast**: Unique identifier for the perturbed gene
- **chunk**: differential expression processing group identifier
- **n_cells_target**: Number of cells with targeting guide for the perturbed gene
- **n_up_genes**: Count of significantly upregulated genes (10% FDR)
- **n_down_genes**: Count of significantly downregulated genes (10% FDR)
- **n_total_de_genes**: Total number of significantly differentially expressed genes (10% FDR)
- **ontarget_effect_size**: Effect size of the perturbation on its intended target gene
- **ontarget_significant**: Boolean indicating whether on-target knockdown was significant (10% FDR)
- **target_baseMean**: Mean baseline expression of the target gene
- **offtarget_flag**: Flag indicating potential off-target effects (TSS within 10 kb with significant down-regulation)
- **n_total_genes_category**: Category based on number of *trans*-effects

- **ontarget_effect_category**: Category based on on-target / off-target effects
- **n_downstream**: Number of genes significantly affected by this perturbation, excluding on-target effect (incoming *trans*-effects)
- **crossdonor_correlation_mean**: Mean cross-donor correlation (Pearson correlation between logFC effects estimated in disjoint pairs of donors). If NA, the perturbation was not tested across donors.
- **crossdonor_correlation_min**: Minimum cross-donor correlation (Pearson correlation between logFC effects estimated in disjoint pairs of donors). If NA, the perturbation was not tested across donors.
- **crossguide_correlation**: Cross-guide correlation (Pearson correlation between logFC effects estimated with individual gRNAs). If NA, the perturbation was not tested across guides.

**Supplementary Table 6** Cross-cell-type comparison of perturbation effects between K562 cells and CD4+ T cells. Each row represents a gene perturbed in both cell types, with correlation analysis of differential expression profiles.

- **target_contrast_gene_name**: Name of the perturbed gene being compared between cell types
- **logfc_pearson_r**: Pearson correlation coefficient comparing log fold change profiles between K562 and CD4+ T cells
- **logfc_pearson_pval**: P-value for the Pearson correlation
- **random_r1**: Pearson correlation with first random perturbation (negative control)
- **random_r2**: Pearson correlation with second random perturbation (negative control)
- **random_r3**: Pearson correlation with third random perturbation (negative control)
- **comparison**: Comparison identifier (e.g., "K562 vs CD4+T (Rest)")
- **condition**: Culture condition for the CD4+ T cell dataset (Rest, Stim8hr, or Stim48hr)
- **donor_correlation_mean**: Mean correlation of log fold change profiles across donors (measure of reproducibility within CD4+T data)
- **n_degs_MASH_K562**: Number of differentially expressed genes (DEGs) at MASH 5% LFSR in K562 cells
- **n_degs_MASH_Rest**: Number of DEGs identified at MASH 5% LFSR in CD4+ T cells (Rest condition)
- **n_degs_MASH_Stim48hr**: Number of DEGs identified at MASH 5% LFSR in CD4+ T cells (48-hour stimulation condition)
- **n_degs_MASH_Stim8hr**: Number of DEGs identified at MASH 5% LFSR in CD4+ T cells (8-hour stimulation condition)

**Supplementary Table 7** DESeq2 results for bulk RNAseq validation of IL10/IL21 regulation.

**Supplementary Table 8** Summary of flow cytometry results for the validation of IL10/IL21 regulation.

- **Sample**: Biological replicates of knockdown sample (format: donor_regulator)
- **IL10_perc**: Percentage of IL10+ cells in the sample
- **IL21_perc**: Percentage of IL21+ cells in the sample
- **Donor**: Unique identifier for the blood donor
- **Perturbation**: The regulator that are knocked down or non-targeting control (NTC, NTC1, NTC2, NTC3)

**Supplementary Table 9** Perturbation clustering results and annotations.

- **cluster**: Unique numeric identifier for the cluster from HDBSCAN.
- **manual_annotation**: Manual annotation for the cluster based on database enrichment, gene ontology analysis, LLM lookup, and manual literature search.
- **intracluster_corr**: Mean intraclass correlation of perturbation effects within the cluster.
- **cluster_size**: Total count of perturbations in the cluster.
- **cluster_gene_size**: Count of unique genes in the cluster.
- **cluster_member**: Unique genes in the cluster.
- **rest_count**: Count of perturbations in 'Rest' condition.
- **stim8hr_count**: Count of perturbations in 'Stim8hr' condition.

- **stim48hr_count**: Count of perturbations in 'Stim48hr' condition.
- **cluster_member_with_condition**: List of specific Gene_Condition pairs.
- **complex_corum**: Top enriched CORUM complex name.
- **overlap_genes_corum**: Genes overlapping with the CORUM complex.
- **overlap_fraction_corum**: Fraction of cluster genes in the CORUM complex.
- **raw_p_value_corum**: Hypergeometric p-value for CORUM enrichment.
- **complex_size_corum**: Size of the CORUM complex.
- **overlap_size_corum**: Count of overlapping genes (CORUM).
- **fdr_corum**: Benjamini-Hochberg FDR for CORUM enrichment.
- **complex_stringdb**: Top enriched STRING cluster ID.
- **best_described_by**: Functional description of the STRING cluster.
- **overlap_genes_stringdb**: Genes overlapping with the STRING cluster.
- **overlap_fraction_stringdb**: Fraction of cluster genes in the STRING cluster.
- **raw_p_value_stringdb**: Hypergeometric p-value for STRING enrichment.
- **complex_size_stringdb**: Size of the STRING cluster.
- **overlap_size_stringdb**: Count of overlapping genes (STRING).
- **fdr_stringdb**: Benjamini-Hochberg FDR for STRING enrichment.
- **complex_kegg**: Top enriched KEGG pathway name.
- **overlap_genes_kegg**: Genes overlapping with the KEGG pathway.
- **overlap_fraction_kegg**: Fraction of cluster genes in the KEGG pathway.
- **raw_p_value_kegg**: Hypergeometric p-value for KEGG enrichment.
- **complex_size_kegg**: Size of the KEGG pathway.
- **overlap_size_kegg**: Count of overlapping genes (KEGG).
- **fdr_kegg**: Benjamini-Hochberg FDR for KEGG enrichment.
- **complex_reactome**: Top enriched Reactome pathway name.
- **overlap_genes_reactome**: Genes overlapping with the Reactome pathway.
- **overlap_fraction_reactome**: Fraction of cluster genes in the Reactome pathway.
- **raw_p_value_reactome**: Hypergeometric p-value for Reactome enrichment.
- **complex_size_reactome**: Size of the Reactome pathway.
- **overlap_size_reactome**: Count of overlapping genes (Reactome).
- **fdr_reactome**: Benjamini-Hochberg FDR for Reactome enrichment.
- **corr_rest**: mean correlation of regulator perturbation effects in Rest condition.
- **corr_stim8hr**: mean correlation of regulator perturbation effects in Stim8hr condition.
- **corr_stim48hr**: mean correlation of regulator perturbation effects in Stim48hr condition.
- **corr_shared**: mean correlation of regulator perturbation effects for all pairwise perturbations that are not between the same condition.
- **condition_specificity**: condition specificity of regulator clusters.

**Supplementary Table 10** Downstream genes of regulator clusters.

- **hdbscan_cluster:** Unique numeric identifier for the cluster from HDBSCAN.
- **downstream_gene:** Name of the downstream target gene identified as differentially expressed (fdr < 0.1) for at least one cluster member regulator.
- **downstream_gene_ids:** Unique gene identifier corresponding to the downstream gene name.
- **num_of_upstream:** Count of cluster member regulators that significantly (fdr < 0.1) perturb the downstream gene.
- **sign_coherence:** Measure of the consistency of regulation direction among significant upstream regulators (where +1 indicates consistent upregulation and -1 indicates consistent downregulation).
- **zscore_rank_negative_regulation:** Rank-based ranking of the downstream gene based on summation of ranks of z-scores across cluster members, prioritizing strong downregulation.

- **zscore_rank_positive_regulation:** Rank-based ranking of the downstream gene based on summation of inverted ranks of z-scores across cluster members, prioritizing strong upregulation.
- **condition:** Experimental condition under which the downstream effects were observed (Rest, Stim8hr, or Stim48hr).

**Supplementary Table 11** DESeq2 results for comparison of Th2 and Th1 gene expression profiles from discovery cohort (Ota 2021) and replication cohort (Hollbacher 2020).

**Supplementary Table 12** Signature prediction model coefficients encoding predicted regulator effects on Th2/Th1 polarization and TCR activation (related to Figure 4F, Suppl. Figure 16G, Suppl. Figure 18B).

**Supplementary Table 13** DESeq2 results for comparison of aging CD4+T cells from discovery and replication cohorts (Yazar 2022).

**Supplementary Table 14** Signature prediction model coefficients encoding predicted regulator effects on CD4+T aging (related to Figure 5D).

**Supplementary Table 15** Enrichment analysis results for autoimmune disease-associated genes within gene clusters derived from perturbation-response profiles.
